# Burst-like Secretion of Platelet Dense Granules Promotes Thrombus Shell Expansion

**DOI:** 10.64898/2026.05.26.727941

**Authors:** Taisia O. Shepeliuk, Anastasiia A. Masaltseva, Roman R. Kerimov, Fazly I. Ataullakhanov, Ekaterina L. Grishchuk

**Affiliations:** Department of Physiology, Perelman School of Medicine, University of Pennsylvania, Philadelphia, PA, USA

## Abstract

Vessel-wall injury triggers platelet recruitment and aggregation with exquisite spatiotemporal regulation. While secreted agonists from thrombin-activated platelets play a crucial role in thrombus formation, the underlying mechanisms remain elusive. Using real-time imaging of isolated human platelets, we demonstrate that dense granules, which are enriched with agonists, are released in brief, stochastic bursts driven by intracellular calcium spikes, which are necessary but not individually sufficient to trigger secretion events. This burst-like secretion is sustained through extracellular feedback, establishing a cooperative, probabilistic mechanism of granule release in which released agonists amplify thrombin-induced granule exocytosis and increase the likelihood of secretion bursts. Computational modeling of whole-thrombus growth reveals that these transient bursts generate localized microdomains of high agonist concentration, facilitating expansion of the outer thrombus layers. Our findings establish burst-like secretion as a distinct hemostatic mechanism that enhances platelet recruitment and orchestrates thrombus architecture through localized, self-reinforcing activation.

**Key points:**

1. Live imaging reveals dense granules secretion in discrete bursts.
2. Intracellular calcium spikes are necessary but not sufficient for triggering secretion events.
3. Burst-like secretion arises from cooperativity, driven by the feedback amplification from the extracellularly released granule content.
4. Burst-like dense granule secretion promotes dynamic expansion of the thrombus shell in silico.

## Introduction

Different types of blood vessel disruption, including direct endothelial damage^1^, denudation due to trauma or compression^2^, stenosis^3^ or ligation^4^, and vessel wall rupture following puncture injury^5–8^, can all induce thrombus formation to limit blood loss and restore vascular integrity. While the mechanisms underlying thrombus formation vary with the nature of the injury, vessel type, and flow conditions, circulating platelets play a universal role by rapidly accumulating at the affected site. This response is tightly regulated by multiple activators. Thrombin generated at the injury site serves as a primary regulator by inducing irreversible platelet–platelet bonding^9^. Consistently, studies of thrombus architecture in the microcirculation of mice, including both laser-induced deep injury and pipette puncture models, show that the thrombus core, where thrombin concentration is highest, consists of tightly packed platelets^9–11^ (Figure 1A). In contrast, the outer layers of the growing thrombus, referred to as the thrombus shell, experience lower thrombin levels due to dilution by blood flow^10^. While some platelets in these layers may undergo irreversible thrombin-induced bonding, many interact through reversible adhesion mediated by secondary agonists secreted from thrombin-activated platelets^12^ (Figure 1A). The core–shell organization has provided a useful conceptual framework for understanding platelet activation gradients, particularly in nonpenetrating or surface-limited injury models. Within this framework, the expansion and remodeling of the thrombus shell remain among the least explored aspects of thrombus formation, especially in humans, where direct observation and manipulation are limited.

**Figure 1.**
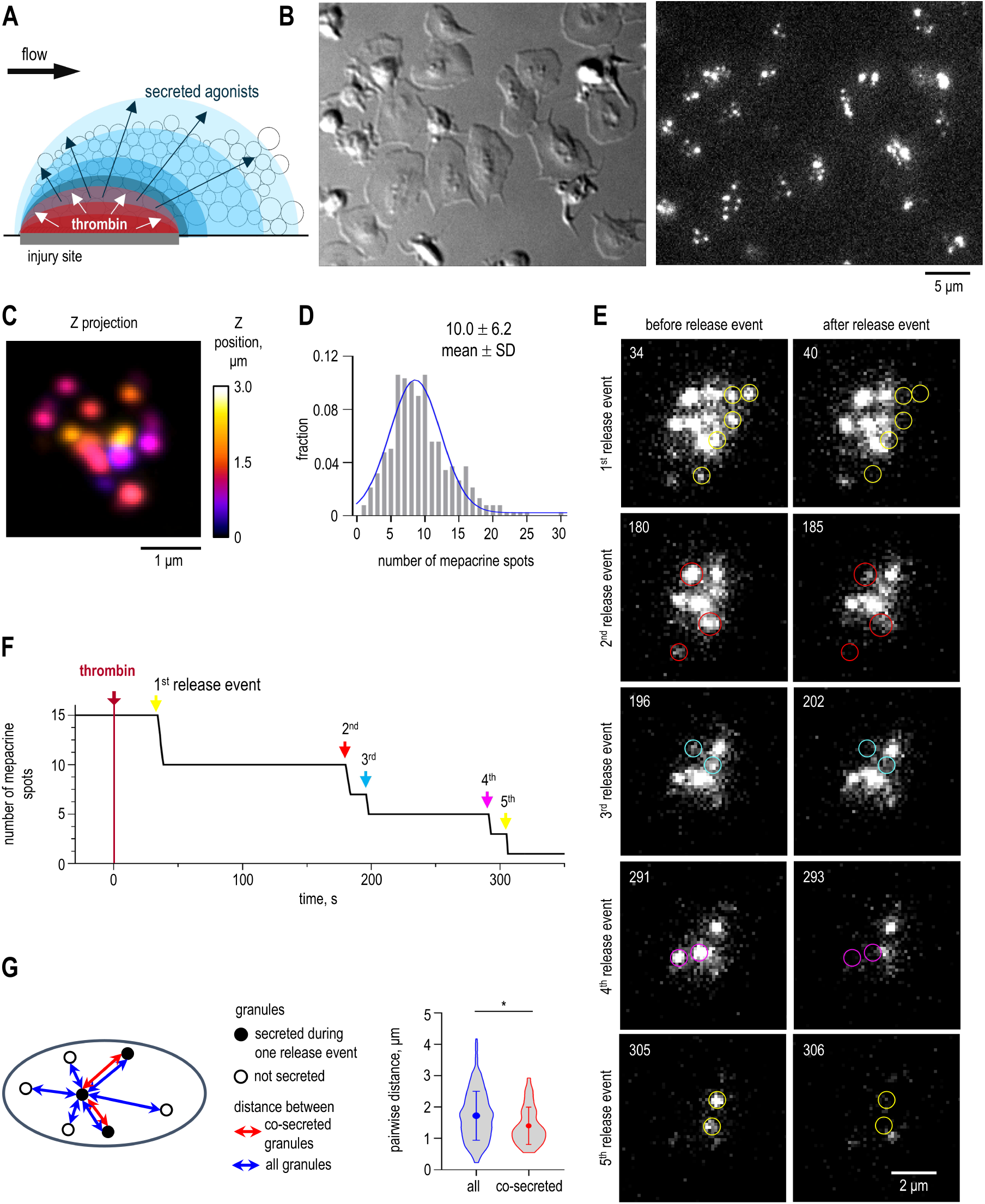
Live imaging of DG secretion in individual platelets. (A) Platelets are represented as empty white circles, with the central thrombus core shown in red and the surrounding shell layers in blue. The thrombin gradient is centered at the site of vascular injury, where thrombin-activated platelets release secondary agonists that are often presumed to establish a secondary, core-centered gradient. However, given the finite nature of granule stores and the strong dilution effects of flow, the actual spatiotemporal distribution of secreted agonists is likely to diverge from this simplified, concentric model. (B) Example field of adhered platelets imaged via DIC (left) and mepacrine fluorescence (right) on a fibrinogen-coated coverslip. (C) Z projection of mepacrine-labeled DGs was captured using a VT-iSIM microscope. Z projection was generated from z-stack with intervals of 150 nm between planes. Each plane was color coded to visualize the z-position of DGs. (D) Distribution of mepacrine spot number in individual platelets measured by VT-iSIM microscope (*N* = 3 independent experiments, *n* = 377 platelets, bin size = 1). The blue curve represents a Gaussian fit. (E) Still images of a representative platelet activated by 0.1 U/ml thrombin at 0 s. Epifluorescence images show mepacrine-labeled DGs. The numbers show time in seconds after thrombin addition. Each pair of images, from top to bottom, show start and end of a release event. Circles indicate mepacrine spots released during same release event. (F) Time dependence of number of mepacrine spots of platelet shown in panel E. Arrows indicate release event shown in panel E. (G) Pairwise distances between platelet DGs are illustrated relative to one secreted granule. In the graph, dots with whiskers represent mean ± SD for *N* = 3, and contours show the distance distribution based on *n* = 16. Mann-Whitney U test: * is p < 0.05.

Among the granules secreted by thrombin-activated platelets, dense granules (DGs) play a particularly important role. DG deficiency, as seen in Hermansky-Pudlak syndrome, leads to significantly reduced platelet accumulation at sites of vessel injury and excessive bleeding^13–17^. DGs contain ADP, ATP, serotonin, and other physiologically important compounds^18,19^. The most extensively studied DG-derived agonist, ADP, promotes further DG secretion from platelets in vitro, suggesting a feedback activation mechanism operating in both autocrine and paracrine manners^20–25^. Notably, ADP induces reversible platelet-platelet adhesion and therefore may serve as a key regulator of shell formation^9,15^. Current models assume that in a growing thrombus, secreted agonist concentration follows the thrombin gradient, which is highest at the injury site (Figure 1A), but the spatiotemporal dynamics of DG secretion and ADP-mediated platelet adhesion remain poorly understood^9,12,15^. If DG-derived agonists, rather than thrombin, primarily regulate platelet adhesion in the thrombus shell, mechanisms must exist to sustain their levels despite DG exhaustion due to secretion, as well as flow-mediated dilution.

Despite the critical role of ADP and other secreted agonists in thrombus formation, the timing and regulatory logic of DG secretion in individual human platelets remain unclear. Current models rely on bulk platelet suspension studies, where kinetic data from millions of cells yield population-averaged responses rather than insights into individual platelet behaviors. In these assays, thrombin addition induces a steady, plateau-limited DG secretion^26–28^, suggesting a monotonic release pattern at the cellular level. However, population-based approaches may obscure key aspects of secretion^29,30,39,31–38^, as seen in other cell types where direct observations reveal granule release in discrete bursts rather than as a continuous process^40–42^. Understanding DG secretion kinetics at the single-cell level is crucial for elucidating how secreted agonists regulate their own release. Additionally, the extent to which DG secretion and feedback activation depend on thrombin and ADP concentrations—both of which fluctuate significantly within a growing thrombus—remains unknown. Agonist concentration may influence not only the fraction of responsive platelets but also the number of DGs released and the timing of secretion, underscoring the need for a quantitative live imaging approach.

Live imaging has already been instrumental in revealing calcium signaling dynamics in activated platelets. While studies in cell suspensions demonstrate a general increase in intracellular calcium upon thrombin activation^43^, single-cell analyses reveal irregular calcium oscillations rather than a steady rise^44–46^. Calcium currents are necessary for DG secretion but how these oscillations relate to granule release and feedback regulation remains unclear^43,47^. Here, we directly visualized DG secretion and calcium dynamics in individual human platelets activated by thrombin and ADP at physiologically relevant concentrations. Our findings reveal, for the first time, that DGs are secreted in distinct bursts comprising several spatially separated granules, strongly supporting the cooperative nature of DG release. We show that feedback regulation increases the number of contemporaneously released granules and increases burst frequency by modulating calcium-dependent triggering. Using an advanced mathematical model of whole-thrombus growth that integrates agonist-modulated platelet adhesion, DG secretion, and flow-dependent agonist diffusion in real time, we demonstrate that secretion bursts play a crucial role in thrombus shell expansion and dynamic remodeling.

## Results

### Several spatially separated DGs are released during a brief burst of secretion

To investigate the kinetics of DG secretion at a single-cell level, we perfused human blood through a custom-made microfluidic flow chamber, which permits live observations under controlled conditions (Supplementary Figure 1). After the platelets settled on the fibrinogen-coated coverslip, the unbound components were removed using a physiological buffer supplemented with the fluorescent dye mepacrine (Figure 1B). Prior reports have established that mepacrine binds to adenosine nucleotides, which are specifically present in dense granules^48,49^. Imaging via epifluorescence microscopy (optical resolution 200 nm) revealed on average 5 mepacrine spots of varying size and brightness per cell. However, a microscope with super-resolution structured illumination capabilities (optical resolution 100 nm) detected 10.0 ± 6.2 mepacrine-positive spots per cell (Figure 1C,D, Supplementary Movie 1), consistent with the upper range of the reported number of DGs in human cells^48^ (Supplementary Note 1). Following other studies that used mepacrine for DG visualization^50,51^, we refer to these dots as DGs.

In chambers with no added activators, 67% of cells retained the same total brightness, and in 33% of cells the brightness decreased by only 20% after 10 mins, indicating little background activation and minimal photobleaching (Supplementary Figure 1C,D, Supplementary Movie 2). However, addition of 0.1 U/ml thrombin led to the disappearance of fluorescent dots in most cells, implying active secretion (Supplementary Figures 1C,E, Supplementary Movie 3). Live imaging revealed that several DGs are often released simultaneously (Figure 1E,F, Supplementary Figures 1F,G and 2), suggesting the coordinated secretion bursts.

Curiously, DGs released simultaneously were not clustered within the same region but were separated by an average pairwise distance of 1.4 ± 0.6 μm (Figure 1G, Supplementary Figure 3A). Although this distance was statistically lower than that between all visible granules in the same cells (1.7 ± 0.8 μm), the distributions were broad and largely overlapping, indicating no clear spatial correlation, and suggesting that co-secreted DGs were only marginally, if at all, closer to each other. In a separate analysis, we found no relationship between the timing of DG release and their distance from the cell boundary (Supplementary Figure 3B,C, Supplementary Note 2). Furthermore, similar stepwise kinetics were observed in both well-spread and rounded cells (Supplementary Figure 3D–H), suggesting that this phenomenon is intrinsic to thrombin-activated platelets and not influenced by cell morphology or inadvertent activation during sample preparation.

### Platelet DG secretion is governed by dual stochastic mechanisms

To gain insight into the temporal regulation of dense granule (DG) secretion, we quantified secretion bursts by tracking changes in total platelet brightness (Supplementary Figures 4A–C). At 0.1 U/ml thrombin, the first secretion burst peaked after a lag of approximately 20 s, consistent with activation delays observed in platelet suspension^28,52,53^. During this initial burst, the decrease in mepacrine fluorescence indicated that platelets released about one-third of their total DG content (Figure 2A,B). In subsequent bursts, the amplitude of brightness decreases diminished, but each event released roughly one-third of the remaining DGs. This pattern suggests a constant probability of release per event, implying similar regulatory mechanisms across sequential secretion bursts.

**Figure 2.**
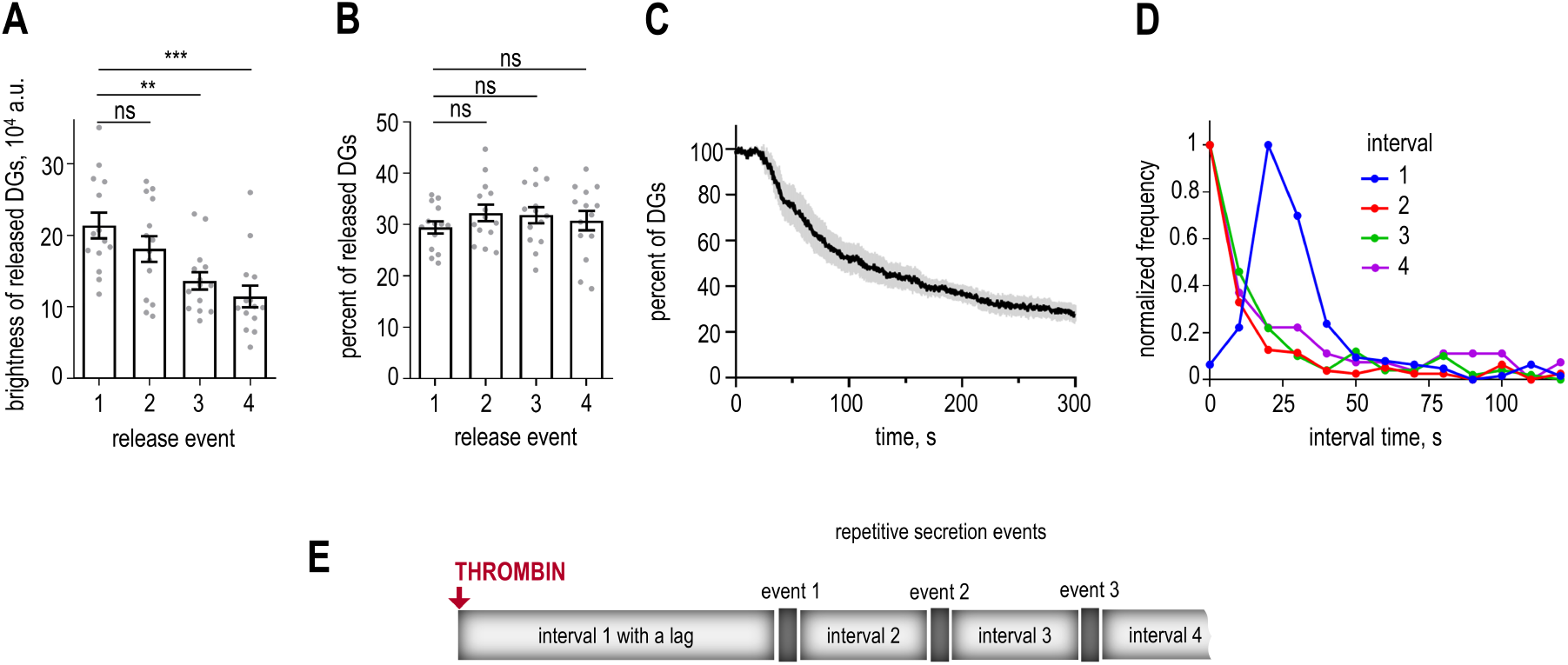
Quantification of DG release kinetics. (A) Change in total mepacrine brightness of platelets activated by 0.1 U/ml thrombin during one release event, representing amount of released DGs during the indicated event. Here and in other graphs, each dot represents the mean result for *n* platelets for each condition. Bars show their mean ± SEM for *N* independent experiments. Here and in panel D: *N* = 14, *n* = 171, 159, 129 and 89 platelets for consecutive events, see Data Source File for more details. Mann-Whitney U test: *** is p < 0.001, ** is p < 0.01, ns – not significant. (B) Relative change in mepacrine brightness of platelets activated by 0.1 U/ml thrombin during one release event, representing percent of released DGs from the remaining DG pool for each secretion event. See legend to panel A for other details. (C) Population secretion curve of percent of the remaining granules over time for secreting platelets activated by 0.1 U/ml thrombin. The black curve is the mean of *N* = 5, *n* = 6, while the grey color is SEM. (D) Distributions of time intervals between the time of thrombin addition and the 1^st^ release event (interval 1), and the intervals between subsequent release events. For easier comparison, all durations were normalized based on the maximum value for each curve. (E) Summary of the DG secretion kinetics in thrombin-activated platelets. Following a lag time, the brief secretion events occur, during which the DGs are released with a constant probability, and these events are interspersed with intervals of stochastic durations.

To assess overall secretion dynamics, we averaged data from multiple cells to generate a population-level secretion curve (Figure 2C). As in previous suspension studies^26–28^, this averaged curve exhibited smooth secretion kinetics. Such smooth population trace results from averaging asynchronous, burst-like secretion events across individual cells, emphasizing the importance of our single-cell analysis.

We next quantified the time intervals between consecutive bursts (Figure 2D). After the initial lag, the first secretion burst followed an exponential timing distribution with a characteristic time of 14 s (Supplementary Figure 4D,E). Subsequent bursts also exhibited exponential timing distributions but without a lag. The similarity in characteristic times across intervals strongly supports a stochastic triggering mechanism. Therefore, DG secretion in platelets is governed by two stochastic processes on different time scales: a relatively infrequent (∼15 s) stochastic triggering of brief (2–4 s) bursting events, during which multiple DGs are released with a constant probability (Figure 2E). The dual stochastic nature of bursting secretion may provide multiple regulatory opportunities for different agonists to modulate the timing and extent of DG release.

### Thrombin and ADP induce secretion bursts with distinct regulatory profiles

Within the developing thrombus, incoming platelets adhere and become activated at various sites, encountering different concentrations of thrombin and other agonists (Figure 1A). To investigate how DG secretion bursts respond to thrombin levels, we systematically varied thrombin concentration over a 1,000-fold range. As thrombin concentration increased from 10⁻⁴ to 1 U/ml, the percentage of secreting cells rose accordingly. Nevertheless, DGs were released intermittently in distinct sets across all concentrations (Figure 3A,B, Supplementary Figure 5A,B). Higher thrombin levels led to an increased number of secretion events (Figure 3C) and shortened time to the first release event in a concentration-dependent manner (Figure 3D). Thrombin’s ability to modulate both the number and timing of secretion events suggests it influences activation pathways governing burst initiation. Notably, higher thrombin concentrations also increased the number of DGs released per event (Figure 3E). Thus, thrombin also regulates pathways affecting the likelihood of DG release within each burst, underscoring its central role in platelet activation^54^.

**Figure 3.**
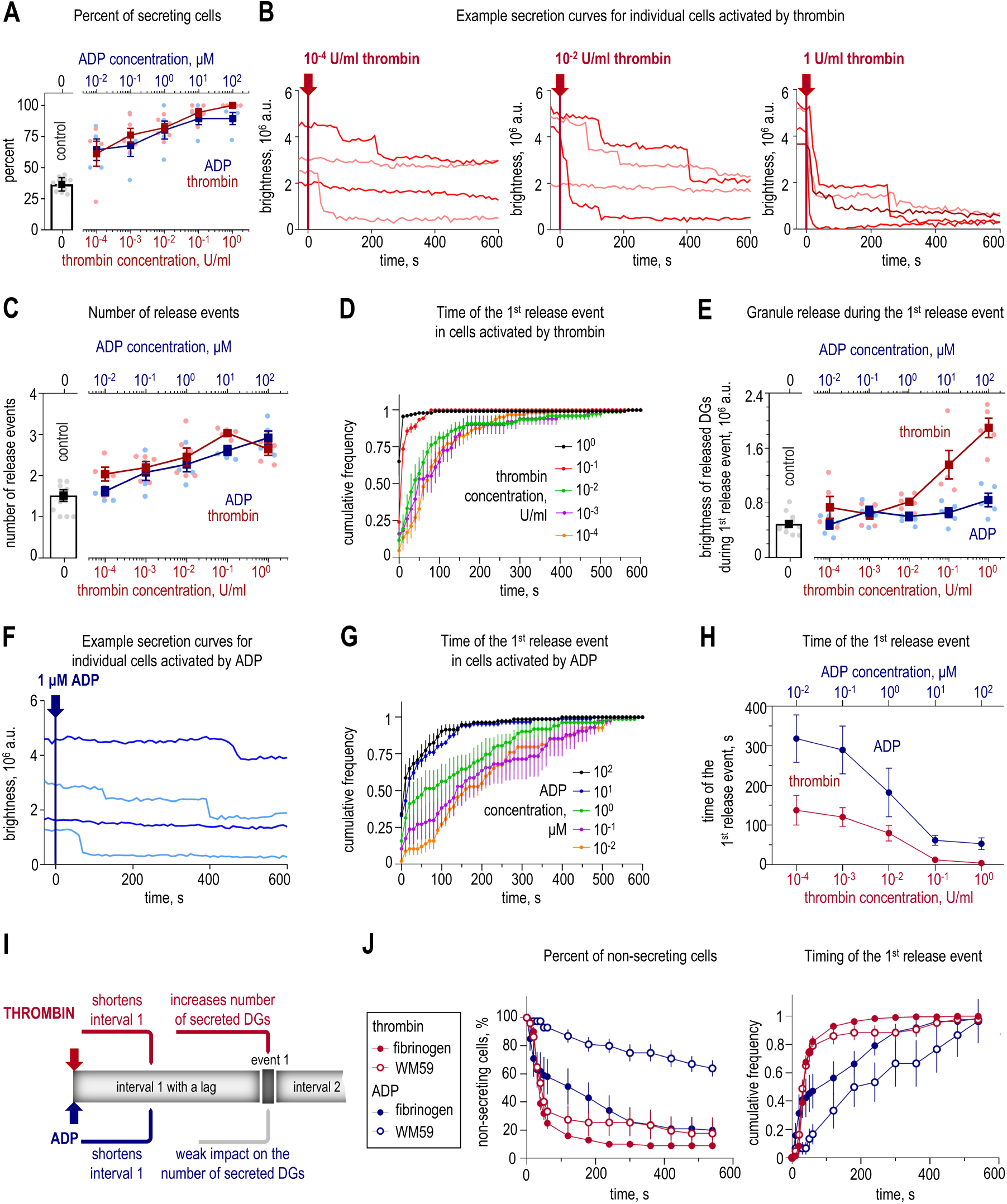
Temporally-coordinated DG secretion in platelets activated by different agonists and adhesive surfaces. (A) Cells that secreted DGs within 600 s after activation with thrombin (bottom axis, light red dots and curve) or ADP (top axis, light blue dots and curve); concentrations are plotted on a semi-log scale. Data for control platelets with no thrombin or ADP are in grey. The number of experiments (*N*) and cells (n) is provided in Supplementary Data Source File, with minimum 4 independent experiments and 58 cells for each condition. Dots represent mean for *n* cells in each experiment; squares with errors represent mean ± SEM based on averaged data from *N* experiments. (B) Changes in total cell brightness with each curve corresponding to one cell. (C) Average number of release events within 600 s after the activation, see legend to panel A. (D) Cumulative frequency distributions of the secretion timing. Each curve was obtained by averaging results of *N* independent experiments, errors are SEM, see legend to panel A. (E) Brightness of released DGs during the 1^st^ release event, see legend to panel A. (F) Changes in total cell brightness. (G) Cumulative frequency distributions as in panel D; *N* = 4 – 5 and *n* = 58 – 94 for each ADP concentration, see Data Source file for more details. (H) Each point shows the time by which 75% of 1^st^ release events occurred, averaged across individual experiments (mean ± SEM) from panels D and G. (I) Summary of the effects of thrombin and ADP on DG secretion in adhered platelets. Thrombin influences both the timing and the quantity of released granules, whereas the influence of ADP is milder, predominantly affecting the timing of events. (J) DG-containing platelets within 600 s after the addition of 0.1 U/mL thrombin or 1 μM ADP (left graph) and cumulative frequency distributions of the secretion timing (right graph); fibrinogen and thrombin (*N* = 22, *n* = 267), WM59 and thrombin (*N* = 5, *n* = 51), fibrinogen and ADP (*N* = 5, *n* = 82), WM59 and ADP (*N* = 9, *n* = 80).

We next asked whether the bursting pattern of secretion was specific to thrombin. In chambers without thrombin, approximately 30% of platelets spontaneously released a small portion of their DGs (Figure 3A, Supplementary Figure 1D). This spontaneous secretion was in distinct steps rather than continuous (Supplementary Figure 5C). To increase secretion frequency, we added ADP. Although on its own, ADP is not a strong inducer of DG secretion in platelet suspension^55^, it triggered robust secretion in our chambers, likely due to the potentiating effect of fibrinogen. Across a wide ADP concentration range (10⁻² – 10² μM), the bursting pattern of secretion was evident (Figure 3F, Supplementary Figure 5D,E). At higher concentrations, ADP promoted DG release comparable to thrombin, ultimately resulting in similar percentages of secreting cells, numbers of secretion events, and DG release per activated cell (Figure 3A,C, Supplementary Figure 5F). Although both 1 μM ADP and 0.1 U/ml thrombin reached similar total secretion levels by 10 minutes after stimulation (Figure 3A,C,E), the rate of secretion was significantly slower with ADP (Supplementary Figure 5G). For example, 75% of first release events occurred within 3 minutes for ADP, compared to just 12 seconds for thrombin (Figure 3D,G,H). Furthermore, while thrombin enhanced the number of DGs released per event with increasing concentration, ADP only modestly increased DG release per event (Figure 3E), suggesting that ADP and thrombin activate secretion through distinct pathways. It appears that thrombin controls both burst triggering and DG release per event, whereas ADP primarily influences burst triggering (Figure 3I).

### Burst-like DG secretion is preserved across distinct adhesion contexts

To test the generality of our findings on burst-like secretion, we next examined dense granule release in platelets adhered via anti-PECAM-1 antibody. Unlike fibrinogen, which promotes platelet spreading and partial activation through integrin αIIbβ3 engagement, adhesion via PECAM-1 antibody (WM59) provides neutral attachment method^56,57^. Platelets adhered to WM59-coated coverslips remained rounded with only occasional filopodia, in contrast to the spread morphology observed on fibrinogen-coated surfaces (Supplementary Figure 5H). The WM59-adhered platelets exhibited almost no spontaneous DG release (Supplementary Figure 5I). Despite these differences, activation of WM59-adhered platelets with either thrombin or ADP induced cell spreading comparable to fibrinogen-adhered platelets. Thrombin-induced DG secretion was only modestly reduced compared to fibrinogen-adhered platelets (Supplementary Figure 5I). In contrast, ADP-induced DG secretion was significantly lower in WM59-adhered platelets (Figure 3J, Supplementary Figure 5J,K), suggesting that fibrinogen enhances ADP-, but not thrombin-induced, secretion. Importantly, under all examined conditions, the burst-like pattern of DG release was preserved (Supplementary Figure 5I), indicating that this dynamic mode of secretion is an intrinsic feature of platelet activation rather than a consequence of a specific agonist or the adhesion context.

### Calcium spikes are necessary but not sufficient to trigger a secretion burst

Intracellular calcium concentration represents a strong candidate for a global signal that may initiate secretion bursts^31,33,36^. Elevated calcium levels are required for DG secretion in platelet suspension^43^, and calcium spiking has been reported in activated platelets^44–46,58^. To examine whether calcium spikes correlate with secretion events, we imaged DG secretion simultaneously with cytosolic calcium using the fluorescent dye Calbryte-590^AM^ (Figure 4A, Supplementary Movie 4). Both thrombin and ADP induced irregular calcium spikes, as expected (Figure 4B, Supplementary Figure 6A-D)^44–46,58^. These spikes originated from intracellular stores, as chelating extracellular calcium with EGTA did not alter the frequency of calcium oscillations or the nature of DG secretion (Supplementary Figure 6E-H), consistent with findings in platelet suspensions^59^.

**Figure 4.**
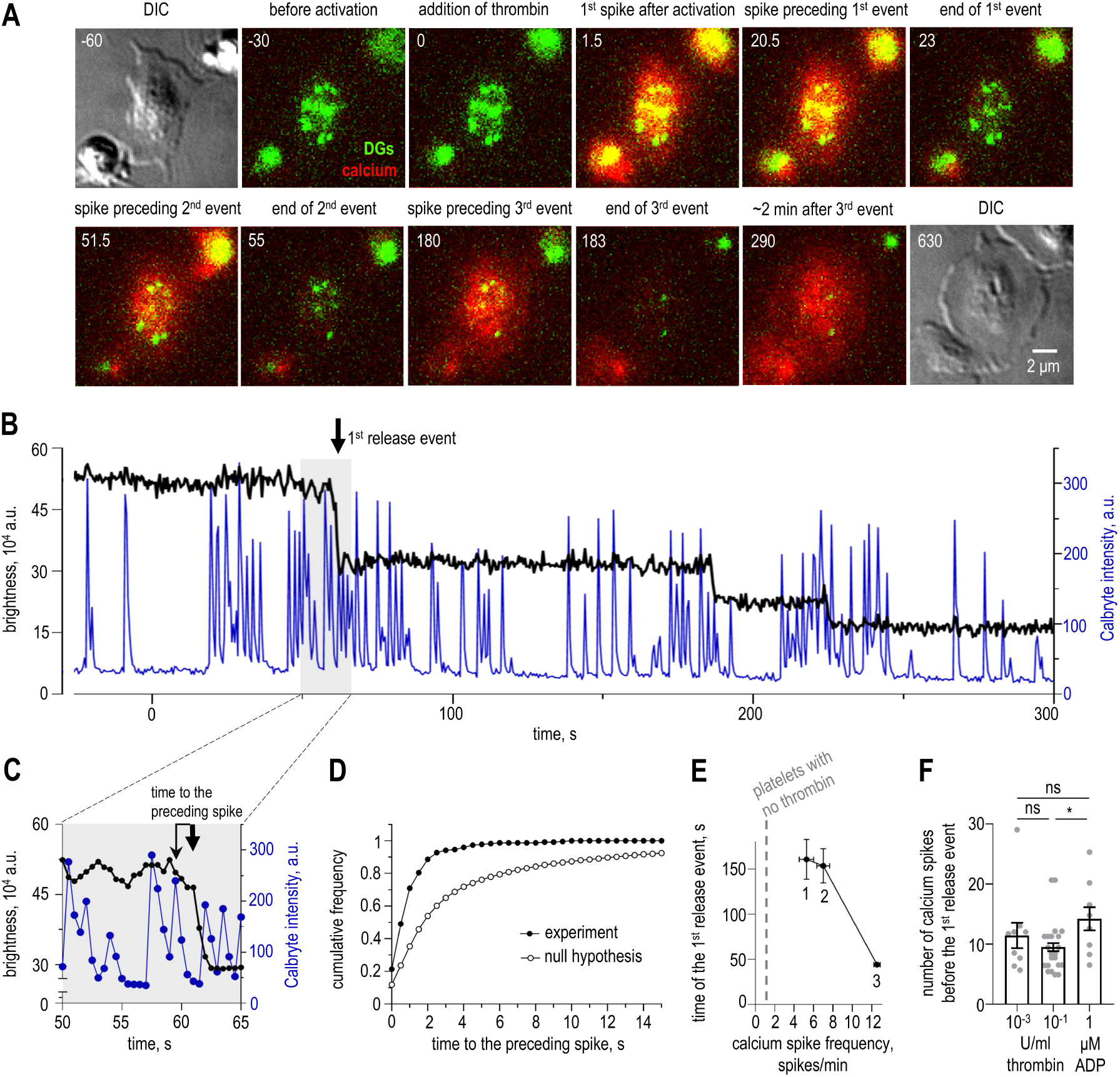
Calcium signaling and DG secretion. (A) Still images of a representative platelet activated by 0.1 U/ml thrombin at 0 s. The first and last images show cell morphology. Epifluorescence images show mepacrine-labeled DGs (green) and intracellular calcium concentration via Calbryte-590^AM^ dye (red). Numbers show time in seconds before (negative numbers) and after (positive numbers) thrombin addition. (B) Changes in the total cell brightness (black, left y axis) and calcium concentration (blue, right y axis) in a platelet activated by 0.1 U/ml thrombin at 0 s. The grey area marks the time window encompassing the 1^st^ release event. (C) Enlargement of the grey time window from panel B. Vertical arrows depict the time span from the initiation of the first DG release event to the occurrence of the calcium spike preceding the event. (D) Cumulative frequency distributions of the time interval between the secretion event and the preceding spike. Black circles correspond to experimental results using platelets activated by 0.1 U/ml thrombin (*N* = 20, *n* = 230). Open circles show expected distribution of intervals between the release event and the preceding spike from the randomly generated oscillations with the same spiking frequency as in the experiment, see Materials and Methods for details. (E) Inverse correlation between the frequency of calcium spiking and the time of the 1^st^ release event (mean ± SEM) in cells activated via different regimes. On the graph, label “1” corresponds to platelets activated by 10^-3^ U/ml thrombin (*N* = 17, *n* = 36), “2” - 1 μM ADP (*N* = 9, *n* = 49), “3” – 0.1 U/ml thrombin *N* = 20, *n* = 230. (F) Number of calcium spikes from the start of activation and until the 1^st^ release event. Each dot corresponds to the average result from each individual experiment, bars show mean ± SEM. Mann-Whitney U test: * is p < 0.05, ns – not significant.

Single-cell imaging revealed that activated platelets exhibited far more calcium spikes than secretion bursts (Figure 4B), indicating that a transient increase in intracellular calcium alone is not sufficient for a bursting event. However, approximately 90% of secretion events occurred within 2 seconds following a spike (Figure 4C,D).

The high frequency and irregularity of calcium spikes made it challenging to visually assess whether this timing was coincidental. For unbiased quantitative analysis, we applied a mathematical algorithm to test the null hypothesis that there is no correlation between spike occurrence and secretion events (Materials and Methods). Using a bootstrap approach, we calculated time intervals between randomly selected time points and preceding calcium spikes. The resulting cumulative distribution for these randomly selected intervals differed significantly from the experimentally measured distribution of times between secretion events and their preceding spikes (Figure 4C,D), thus rejecting the null hypothesis and demonstrating a strong temporal association between calcium spikes and secretion events. These results support the conclusion that calcium spikes are necessary but that individual spikes are not sufficient to trigger secretion bursts.

### Secretion bursts are triggered stochastically by calcium spikes

Since not every calcium spike was followed by a DG burst, we examined whether calcium spikes that directly preceded the first secretion event had distinguishing features. There was no correlation between DG secretion and the amplitude of the preceding spike or the frequency of spiking between the addition of the activator and the first secretion event, suggesting that calcium spikes induce secretion events in a stochastic manner (Supplementary Figure 7, Supplementary Note 3). At 0.1 U/ml thrombin, only 1 in 10 calcium spikes was followed within 2 s by a secretion event, giving a triggering probability p_1_ = 0.1. We then investigated whether this probability varies with the degree or method of platelet activation. At a lower thrombin concentration (10⁻³ U/ml), which induces slower DG secretion, calcium spikes occurred significantly less frequently (Supplementary Figure 6B,D), revealing an inverse relationship between spike frequency and the timing of the first release event (Figure 4E), such that higher spike frequency was associated with reduced time before secretion onset. Notably, the number of spikes preceding a secretion event did not change significantly (Figure 4F), implying that the occurrence probability p_1_ remains independent of thrombin concentration. Moreover, in cells activated with 1 μM ADP, which induces secretion with kinetics similar to 10⁻³ U/ml thrombin, the spike frequency before the first release event was only slightly lower, resulting in a similar p_1_ = 0.07 (Figure 4E,F, Supplementary Figure 6C,D). Thus, secretion events are triggered with a consistent probability across different activation conditions that elicit irregular calcium oscillations with distinct frequencies.

We next examined whether the amount released during a burst depends on features of the calcium spike. There was no significant correlation between the number of DGs released per burst and either the amplitude of the preceding spike or the cumulative calcium signal, measured as the area under the calcium curve prior to the release event (Supplementary Figure 7B,D). Together, these findings indicate that once calcium spikes are induced in activated platelets, the occurrence of subsequent secretion bursts is governed by stochastic calcium-dependent triggering rather than by specific upstream molecular events that generate the oscillatory spikes. Moreover, calcium spikes do not regulate the number of granules released, which is instead largely controlled by thrombin level.

### DG-derived agonists accelerate thrombin-induced DG secretion and increase the likelihood of calcium-dependent secretion bursts

The reported here results illuminated the regulatory principles underlying the triggering of secretion bursts but did not clarify why multiple granules are released within a single burst. Such coordinated secretion may result from release cooperativity, where the secretion of an initial granule facilitates the release of additional DGs. This positive feedback could originate intracellularly or involve the secreted content acting extracellularly on platelets. We focused on the latter, given that DG secretion is known to cause a rapid, transient increase in the local extracellular concentrations of platelet activators, including ADP and ATP, and serotonin^19,60,61^.

To experimentally assess how released DGs amplify secretion, we activated a suspension of washed platelets with 0.1 U/ml thrombin, centrifuged the cells, and collected the supernatant, referred to here as the “secreted content” (SC) (Figure 5A). We then added the SC to coverslip-adhered platelets in our real-time microscopy assay. In the presence of SC, both the number of DGs secreted during the first bursting event and the speed of secretion initiation increased relative to thrombin activation alone (Figure 5B,C). Additionally, the first release event occurred after only half the number of calcium spikes compared to activation with thrombin alone (Figure 5D), indicating that SC significantly enhances the probability of calcium-dependent triggering of secretion (∼0.22). In contrast, the control supernatant from washed platelets had no detectable effect (Figure 5A-D). We also confirmed that the amplification observed with SC was not due to residual thrombin in the preparation (Figure 5A-D). These findings strongly support an extracellular feedback mechanism in which thrombin-activated platelets release factors that amplify DG secretion. Specifically, in thrombin-activated platelets, addition of the released content accelerates secretion onset and increases its magnitude by raising calcium spike frequency and enhancing platelet sensitivity to calcium.

**Figure 5.**
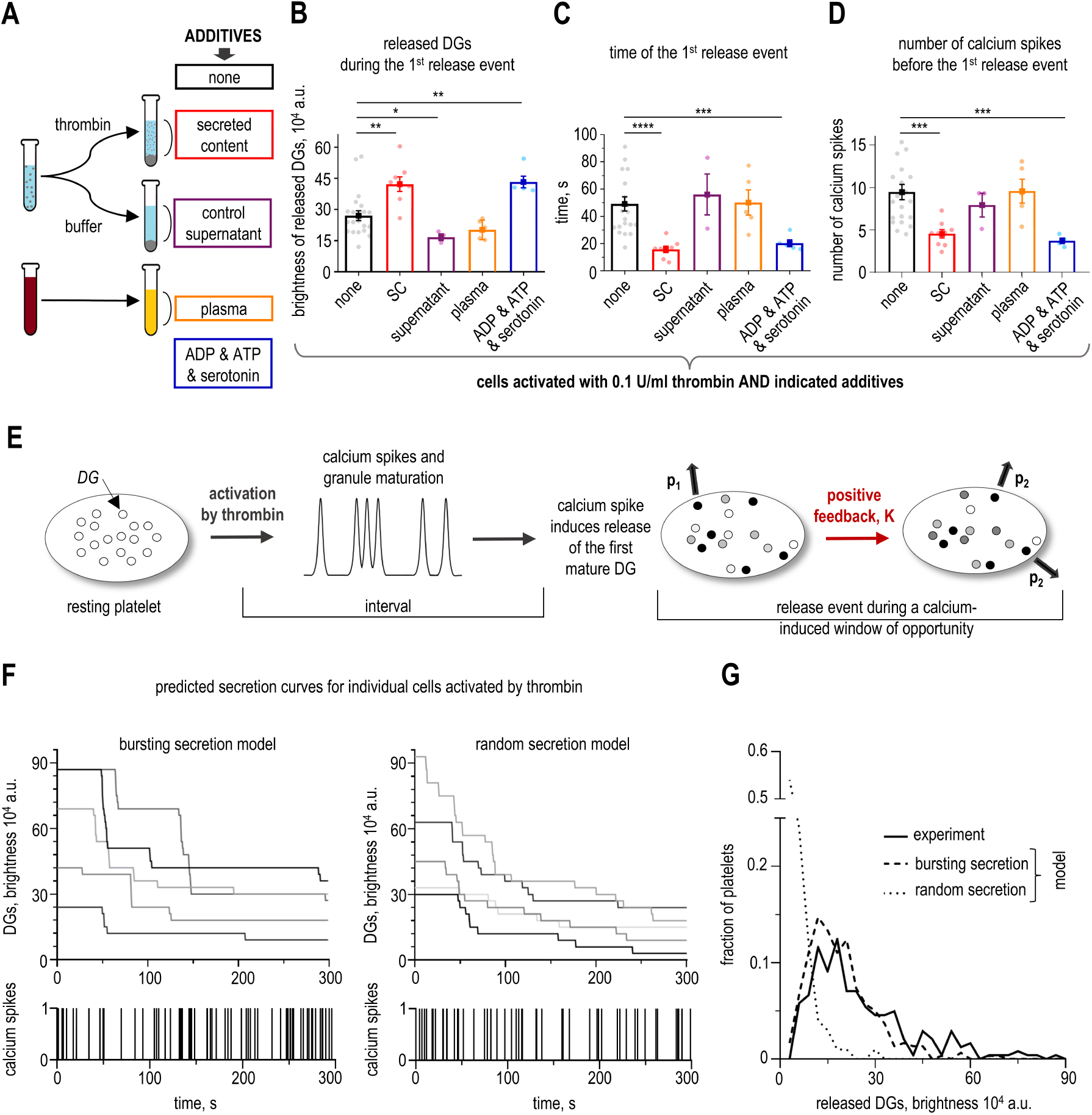
Cooperativity of DG secretion. (A) Experimental design to generate different additives to test their effect on thrombin-activated secretion. (B-D) Secretion and calcium spikes in isolated platelets activated by 0.1 U/ml thrombin and indicated additives: none – no additives (*N* = 21, *n* = 230), SC – secreted content (*N* = 9, *n* = 98), “supernatant” (*N* = 3, *n* = 38), “plasma” (*N* = 5, *n* = 84), “ADP & ATP & serotonin” (*N* = 5, *n* = 75). Mann-Whitney U test: ** is p < 0.01; * is p < 0.05. If columns are not compared, there is no significant difference. (B) Brightness of released DGs during the 1^st^ release event. (C) Time of the 1^st^ release event. Here and in panel E, Mann-Whitney U test: **** is p < 0.0001, *** is p < 0.001. If columns are not compared, there is no significant difference. (D) Number of calcium spikes before the 1^st^ release event. (E) Activation induces irregular calcium spikes and granule maturation. Shades of gray indicate granule release probability, with black corresponding to maximum probability. Once a mature granule is stochastically released with probability p_1_, positive feedback amplifies this probability by a factor K, increasing the secretion probability p_2_ for the remaining mature granules, which are then secreted during the same release event. (F) Comparison of the predictions for granule secretion in different secretion models. Each curve shows change in DG brightness in one modelled cell. The bottom graphs depict the corresponding calcium spikes. (G) Distributions of the brightness of granules released during the 1^st^ release event in platelets activated by 0.1 U/ml thrombin. Experimental distribution is for *N* = 20, *n* = 230; theoretical results are based on 100 simulations for each model.

Using this novel single-cell activation assay, we investigated potential molecular mediators of this regulatory feedback. We hypothesized that DG-derived agonists might be sufficient to amplify secretion or, alternatively, that secretion of other granule types could also contribute. To distinguish between these possibilities, we activated platelets with 0.1 U/ml thrombin in combination with selected agonists highly enriched in DGs. ADP and serotonin were applied at 20 μM, approximating their concentrations in the supernatant of secreting platelets (Supplementary Note 4). Neither ADP nor serotonin alone enhanced thrombin-induced DG secretion, nor did ATP, even when applied in excess (1 mM) (Supplementary Figure 8A–C). Strikingly, the combination of all three compounds (20 μM ADP, 20 μM serotonin, and 13 μM ATP) elicited a strong secretion response that was statistically indistinguishable from the effect of SC (Figure 5A–D, Supplementary Figure 8D). The frequency of calcium spikes in cells activated by this mixture closely matched that observed with SC (Supplementary Figure 8E), and the probability of calcium-dependent triggering of secretion was also similar (0.27 vs. 0.22). These results suggest that DGs themselves can drive cooperative amplification through a combination of ADP, serotonin, and ATP, leading to a secretion burst during which multiple DGs are released.

### Burst-like DG secretion promotes rapid expansion of the thrombus shell *in silico*

Previous work has proposed that at the site of vascular injury, platelets aggregate and initiate DG release in response to local thrombin concentration, with the kinetics and amount of DG secretion further modulated by local agonists^9,10,28^ (Figure 1A). Studying these processes in humans is challenging. To overcome this barrier, we developed a mathematical model of thrombus formation, which incorporates our quantitative findings on DG secretion and its calcium-dependent triggering.

First, a model of granule secretion in individual cells was constructed (Figure 5E, Supplementary Materials, Part 1. Modeling of DG secretion in platelets). Since our experiments revealed that DGs are not secreted all at once, we included in the model an explicit mechanism that limits secretion burst size. We hypothesized that similar to other secretory cells, DGs likely differ in their readiness for release, a process often referred to as maturation, and regulated by calcium levels^31,35,36,62,63^. In the model, DG maturation is a gradual process that increases the pool of release-ready granules after activation. Additionally, the model incorporates a brief “window of opportunity” following each calcium spike, during which only these mature granules can be released. The cooperativity of DG release was implemented with a positive feedback loop, which promotes additional granule secretion with increased probability p_2_=Kp_1_, where K is the feedback factor. Using these experimentally derived assumptions and parameter values (Supplementary Table 1), our model effectively reproduced secretion in individual human platelets, including the number of DGs secreted during the initial release event (Figure 5F,G), the characteristics of DG secretion in response to different agonists and concentrations, and secretion kinetics across the platelet population (Supplementary Figures 9–12). In contrast, a model variant in which calcium spikes triggered random DG secretion predicted continuous, non-bursting secretion and did not match our experimental data (Figure 5F,G).

Next, we integrated this cellular DG secretion model with a published model of microvascular thrombus formation^64^. At the start of each simulation, platelets flow along the vessel segment and progressively adhere to a monolayer of surface-bound platelets; this monolayer serves as the initial condition for thrombus growth (Figure 6A, Supplementary Figure 13). Thus, the model does not include a description of a specific injury mechanism or the initial events leading to platelet adhesion. This monolayer represents the ‘injury’ site, where thrombin is continuously generated and diluted by blood flow, forming a steep concentration gradient (see Supplementary Materials, Part 2: Modeling of thrombus formation).

**Figure 6.**
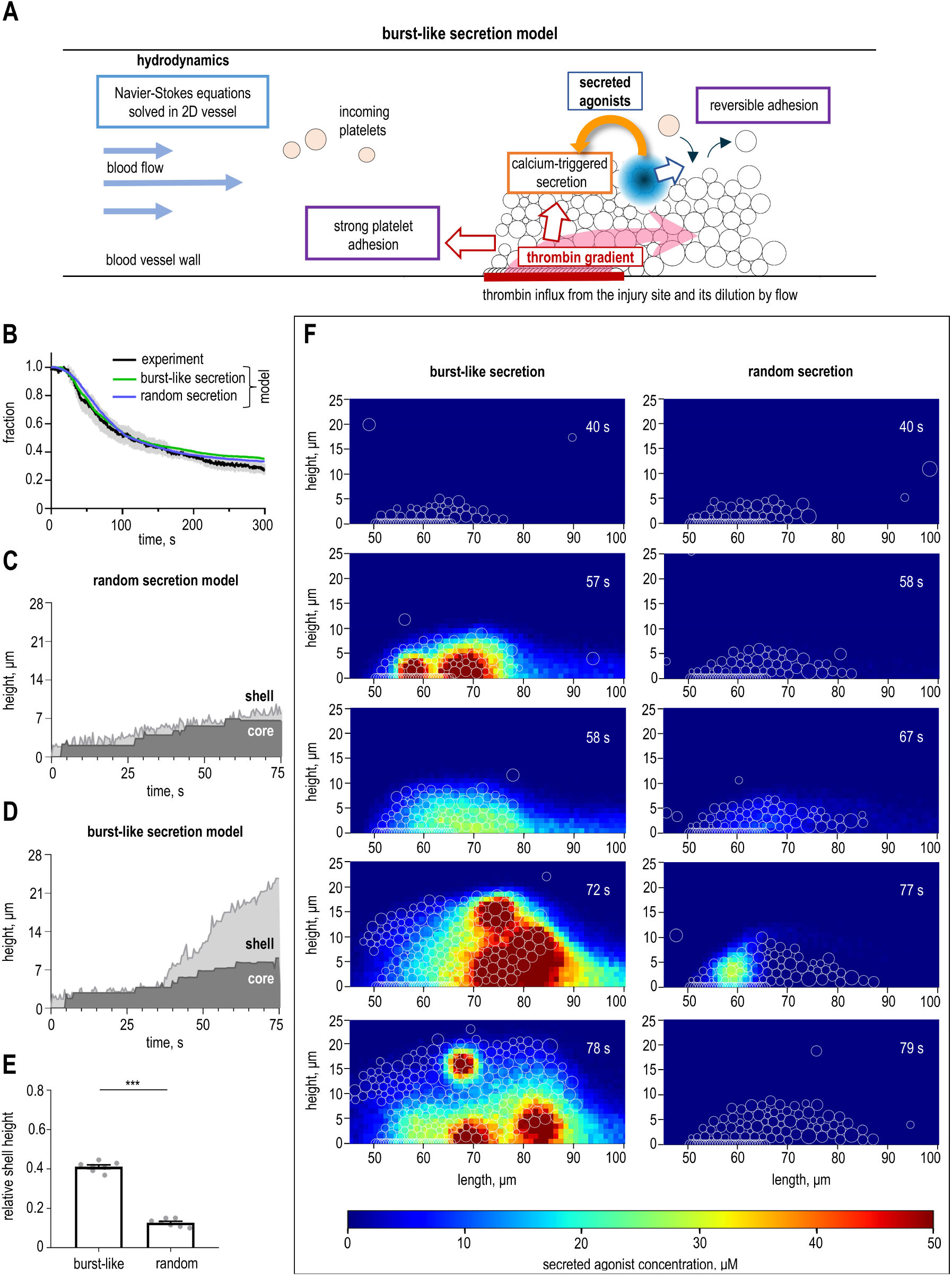
Computational model of thrombus growth. (A) Main model components. The concentration of thrombin and secreted agonist is calculated in real time for each platelet, determining its activation level, which in turn regulates DG secretion. Aggregation is controlled by both thrombin and secreted agonist. This schematic illustrates cooperative section, with positive feedback for secretion illustrated with orange arrow. (B) Population secretion curves represent time-dependent fraction of DGs in platelets activated by 0.1 U/ml thrombin. Simulated curves are mean result for 300 individual platelet secretion curves. for random secretion model (green) and bursting secretion model (blue). Experimental data (black) is the same as in Figure 2C. (C, D) Representative growth curves for thrombi in two models, depicting zoned composition. (E) The maximal shell height at the end of simulations (N=7 for each model) normalized by the vessel width. Bars are mean ± SEM. Mann-Whitney U test: *** is p < 0.001. (F) Still images show changes in the spatial agonist distribution in the growing thrombi. Numbers correspond to the time in seconds after the start of the simulation.

Two distinct platelet adhesion mechanisms were included: reversible interactions mediated by SC and thrombin-dependent irreversible adhesion^65,66^. The extent and type of platelet–platelet adhesion in the model are dictated by the local thrombin concentration, which is dynamically computed at each coordinate. Thrombin also induces intracellular calcium spikes in a concentration-dependent manner, triggering DG secretion with an experimentally determined probability. The local concentration of the released SC, modeled as a single effective molecular entity for simplicity, is determined in real time. DG secretion probability and calcium spike frequency are then adjusted according to the feedback mechanism observed experimentally (Figure 5A–D).

Using this quantitative framework, we examined the physiological implications of different secretion patterns for thrombus formation. When calcium triggered random secretion of individual DGs, thrombus size increased steadily, with most platelets aggregating in response to thrombin rather than SC (Figure 6B,C, Supplementary Figures 14, 15A–C, Supplementary Movies 5, 6). In contrast, when secretion was burst-like, the dynamics of thrombus growth differed markedly. Although thrombi in both scenarios initially assembled at similar rates, burst-like secretion accelerated platelet aggregation, ultimately producing a significantly larger thrombus (Figure 6C–E, Supplementary Figures 14, 15D–F, Supplementary Movie 7). Notably, DG secretion kinetics remained similar between the two models under constant thrombin concentrations (Figure 6B), strongly indicating that the observed differences in thrombus growth arose specifically from distinct SC secretion patterns.

We analyzed the spatial patterns of platelet activation within these thrombi and found that their central cores, where thrombin-dependent activation dominates, remained similar in size (Supplementary Figure 15G–I). This finding suggests that platelet aggregation near the injury site is largely unaffected by the DG secretion pattern^9,15^. When DGs were secreted randomly, the overall extracellular concentration of SC remained low throughout the thrombus. Because agonists were released gradually rather than in bursts, their local accumulation rarely reached levels sufficient to sustain feedback activation, and blood flow readily dispersed them before they could significantly influence secretion in the same or neighboring cells (Figure 6F). In contrast, burst-like DG secretion produced transient but highly concentrated foci of SC that persisted longer, effectively engaging positive feedback regulation (Figure 6F). This effect was most pronounced in the shell layers, where incoming platelets still retained their DG stores and thus secreted profusely when triggered. Consequently, platelet activation in the outer thrombus layers was driven primarily by SC rather than thrombin (Supplementary Movie 8). The adhesion between outer platelet aggregates was transient, as evidenced in the model by platelet rolling and flow-induced deformation of the external layers (Supplementary Movie 7). Occasionally, the entire shell detached under flow (Supplementary Figures 15D–F), suggesting a self-limiting mechanism that may help prevent vessel occlusion. Collectively, these results indicate that the burst-like kinetics of DG secretion play a key role in thrombus growth by controlling expansion of the dynamic outer shell.

## Discussion

Exquisitely controlled DG secretion is central to physiological platelet function during hemostasis and thrombosis^13–16^, as well as in inflammation and immunity^67^. However, the mechanisms that coordinate DG release at the single-cell level, and how this activity is spatially and temporally organized within a growing thrombus, remain unclear. Here, we integrated single-cell imaging of human platelets with quantitative modeling of thrombus growth on a platelet monolayer to connect cellular secretion behavior with the collective dynamics of platelet assembly. Although our model does not explicitly capture all types of vascular injury, such as penetrating or profuse bleeding, the combined experimental–computational approach provides a mechanistic framework for linking local secretion control to thrombus-scale organization.

Our real-time imaging revealed that DG secretion occurs in discrete, transient bursts rather than as a continuous process. When individual secretion traces are averaged across cells, they reproduce the smooth, exponential profiles previously observed in population assays, reconciling our findings with earlier bulk studies^26–28^. The evidence for burst-like secretion arises from direct visualization of secretion dynamics in individual platelets and from quantitative changes in total mepacrine signal after activation. The presence of bursts in cells adhered to different molecular surfaces and activated by different agonists strongly suggests that burst-like secretion is an intrinsic feature of activated human platelets.

We found that this cooperative secretion mode operates across a broad range of agonist concentrations. Thrombin levels primarily determine the number of DGs released per event, whereas ADP levels exert a weaker but modulatory effect that is enhanced by platelet adhesion to fibrinogen (Figure 7A). These findings reveal an unexpected complexity in DG secretion control, indicating that different activators and their combinations regulate the frequency and size of secretion bursts rather than simply the overall amount released.

**Figure 7.**
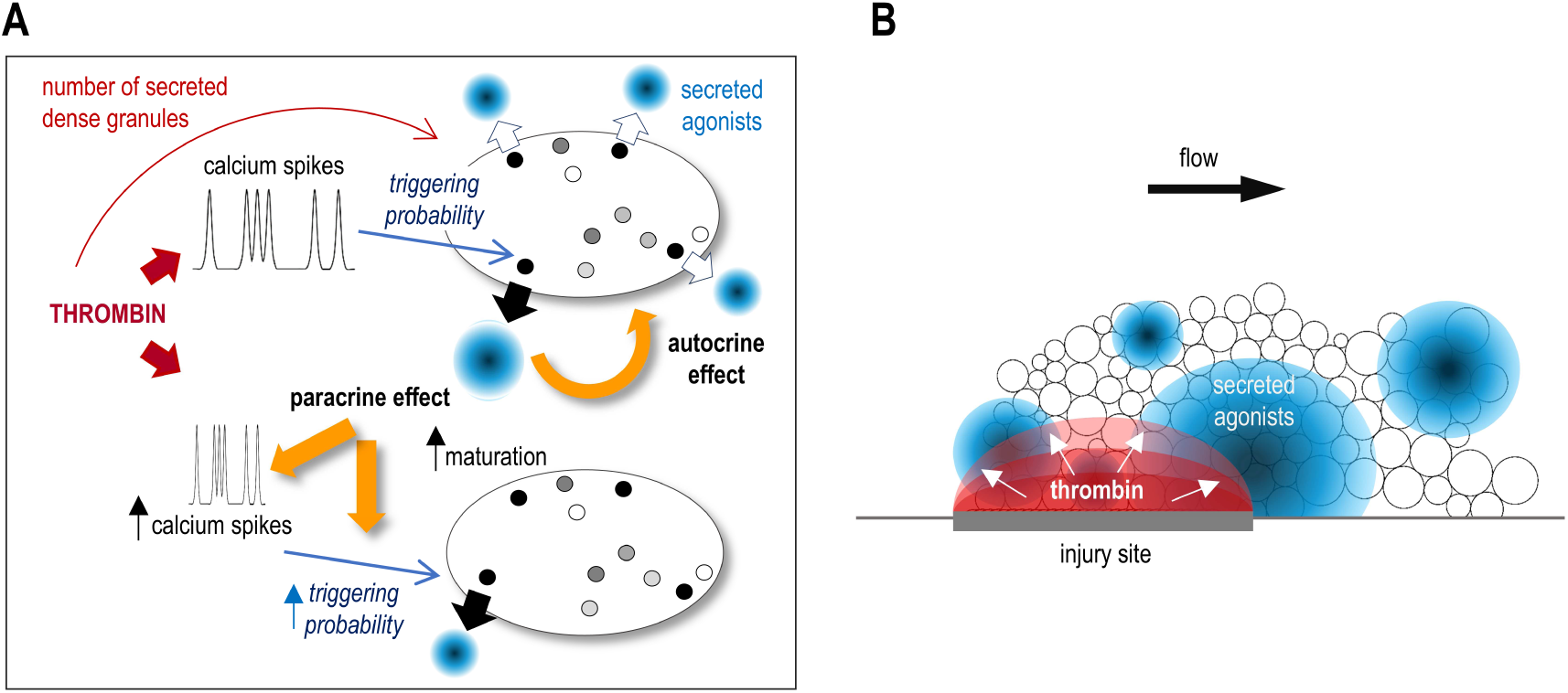
Graphical summary of main conclusions. (A) Regulatory logic of cooperative DG secretion governed by dual stochastic mechanisms. Platelet activation by thrombin induces calcium spikes, which trigger infrequent DG release (a black dot with a black arrow). The content of the first released granule activates positive feedback, promoting rapid release of the remaining mature granules within the same cell (autocrine effect; black dots with open arrows). In nearby cells (paracrine effect), the secreted agonists enhance secretion by increasing the frequency of calcium spikes, which in turn increases the probability of granule maturation, and independently increases the probability of triggering first-granule release. (B) Secreted agonists create transient, sporadic microdomains of high concentration within the growing thrombus. These localized agonist foci enhance platelet aggregation, counteracting the dilution effects of blood flow, and expanding the growing thrombus.

By simultaneously visualizing intracellular calcium and secretion dynamics, we established that stochastic calcium spikes trigger DG release with weak dependence on the specific activation pathway or agonist concentration. This stochastic calcium control parallels secretion mechanisms in other excitable cells, such as neurotransmitter release in neurons^33^. Interestingly, platelets typically release only a subset of their DGs during a given burst. This limitation may stem from the transient nature of calcium-induced secretion potentiation, a preference for releasing only the “mature” DGs, or both. In our model, secretion kinetics were best reproduced by assuming that each DG exists in either an immature or mature state, with the probability of maturation increasing monotonically and plateauing following platelet activation. Mature granules can then be released stochastically during brief, calcium-spike–linked time windows. Although direct experimental evidence linking DG maturation and stochastic triggering to burst size is currently lacking, these mechanisms provide testable hypotheses for future studies.

Mechanistic studies in other cell types offer insights into processes that could similarly operate in platelets. In many secretory cells, granule maturation renders vesicles competent for exocytosis, for example via transitions from docked to primed states that prepare them for rapid calcium-triggered release^68–71^. In neurons, vesicle secretion is influenced by proximity to the plasma membrane and the priming of a “readily releasable” pool^34^. In platelets, we did not observe obvious spatial constraints on DG secretion; however, given the small cell size and the resolution limits of fluorescence microscopy, subtle effects related to granule positioning—potentially involving the open canalicular system—cannot be excluded. Calcium microdomains, which regulate secretion in some cell types^31,33,35^, are also unlikely to play a dominant role in coordinating DG release in platelets, as we observed a relatively uniform cytoplasmic calcium distribution that fluctuates strongly upon activation^72^. In this regard, DG release regulation may resemble pulsatile insulin secretion in pancreatic cells, which is coordinated with global cytoplasmic calcium oscillations^42,73^. Granule proximity to the open canalicular system^74–76^, SNARE complex assembly and docking^47^, and other factors could influence the timing and number of secreted DGs and merit further investigation.

A key advance of this study is the characterization of a positive feedback mechanism that drives cooperative DG secretion. In thrombin-activated platelets, the released granule content accelerates secretion onset and amplifies its magnitude (Figure 7A). This feedback can be reproduced by a mixture of ADP, ATP, and serotonin, suggesting that the DGs themselves reinforce their own secretion. Mechanistically, these agonists increase both the frequency of calcium spikes and the sensitivity of secretion machinery to calcium, elevating the probability and amplitude of secretion bursts. They also enhance the number of DGs released per burst, consistent with an increase in the feedback factor K, which could reflect modulation of downstream exocytic machinery. The molecular mechanisms underlying this modulation, as well as how granule secretion is synchronized on such short timescales, remain unclear. One plausible pathway involves membrane depolarization via the ATP-gated P_2_X_1_ channel^77^, which could rapidly propagate activation signals across the platelet membrane to synchronize granule release. Analogous electrical coupling has been described in neurons and neuroendocrine cells^63,78^, suggesting a conserved role for membrane potential in coordinating vesicular secretion. Future work combining temporally resolved DG imaging with membrane potential measurements will be crucial to test this idea.

The integration of these quantitative findings into a mechanistic thrombus model provides new insight into how platelet-level secretion dynamics shape thrombus architecture. In silico simulations revealed that burst-like DG secretion generates transient, localized zones of high agonist concentration that persist long enough to activate nearby platelets and sustain positive feedback. These microdomains emerge stochastically, challenging the classical view of a smooth agonist gradient dictated solely by thrombin diffusion (Figure 7B). Instead, localized bursts produce self-reinforcing patches of activation that promote recruitment of loosely adhered platelets and drive expansion of the thrombus shell. This mechanism provides a physically consistent explanation for how the outer thrombus layers remain dynamically active despite low thrombin concentrations and flow-mediated dilution.

In summary, these mechanistic insights provide a new framework for understanding the spatiotemporal dynamics of clot growth, an aspect of hemostasis that cannot be directly studied in humans. While our model does not capture all types of vascular injury, such as deep puncture wounds, the principles identified here—cooperative secretion, calcium-dependent stochastic triggering, and extracellular feedback amplification—are likely relevant across diverse hemostatic contexts. The extent and regulation of cooperative DG secretion, including its modulation by blood flow and vascular geometry, remain important directions for future investigation. More broadly, recognizing burst-like DG release as a distinct hemostatic mechanism highlights opportunities to selectively modulate platelet secretion in thrombosis and bleeding disorders, addressing a long-standing therapeutic gap.

## Materials and Methods

### Human ethics

Collection of human blood were approved by the by the University of Pennsylvania Institutional Review Board (№ 855449). Informed consent was obtained from all healthy donors in accordance with the Declaration of Helsinki.

### Reagents preparation

Fibrinogen (Sigma Aldrich, F4883, USA) was added at 50 mg/ml to PBS (137 mM NaCl, 2.7 mM KCl, 10 mM Na_2_HPO_4_, 1.8 mM KH_2_PO_4_, pH 7.4) and incubated for 2 h at room temperature on a shaker. The mixture was centrifuged at 16,800 g for 30 min, and 90% of the top solution was retained. This solution (1 ml) was dialyzed overnight against 400 ml PBS at 4°C, and centrifuged as above. Final fibrinogen solution was aliquoted, flash frozen in liquid nitrogen and stored at −80°C. On the day of the experiment, one aliquot of fibrinogen was thawed at room temperature, diluted to 1 mg/ml in PBS, and centrifuged as above, and used at 1 mg/ml because high density of fibrinogen has been shown to minimize background platelet activation^79^. Calcium-specific dye Calbryte-590^AM^ (AAT Bioquest, 20702, USA) was dissolved at 10 mM in DMSO, aliquoted, flash frozen in liquid nitrogen and stored at −80°C. On the day of experiment, one aliquot of Calbryte-590^AM^ was thawed at room temperature, diluted to 40 μM in Buffer A (150 mM NaCl, 2.7 mM KCl, 1 mM MgCl_2_, 0.4 mM NaH_2_PO_4_, 20 mM HEPES, 5 mM glucose, pH 7.4) and stored on ice in the dark until using in experiment. WM59 monoclonal antibody against human CD31 (Bio-Rad, MCA1738T, USA) was diluted to 400 µg/ml in ice-cold PBS containing 20% glycerol (MP Biomedicals, 151194, USA), aliquoted, and stored at −80°C. On the day of the experiment, one aliquot of WM59 was thawed at room temperature and further diluted to 40 µg/ml in PBS. Annexin V-Alexa Fluor^647^ (BioLegend, 640912, CA) and CD62P-Alexa^647^ (Biolegend, 304918, USA) were stored at 4°C. Stock solution of 200 mM ethylene glycol tetraacetic acid (EGTA, Sigma-Aldrich, E4378, USA) in 1 M NaOH was aliquoted, frozen and stored at −20°C. Stock solutions of 10 mM mepacrine (Aldrich, Q3251, USA), thrombin at 20 NIH units (Sigma-Aldrich, T8885, USA), 20 mM ADP disodium salt (MP Biomedicals, 100056, USA), 100 mM ATP disodium salt (Sigma-Aldrich, A7699, USA), 4.7 mM serotonin hydrochloride (Sigma-Aldrich, H9523, USA) were prepared in milliQ water, aliquoted, flash frozen in liquid nitrogen, frozen and stored at −80°C. On the day of the experiment, the relevant aliquots were thawed at room temperature, spun briefly and kept on ice until using in experiment but no longer than 3 h.

### Flow chamber preparation

#### Master mold and PDMS preparation

Master mold was prepared using standard soft lithography technology at the Quattrone Nanofabrication Facility, University of Pennsylvania, USA. The mold has 2 parallel channels, each with dimensions 1 mm, 13 mm and 50 μm for width, length and height, respectively. Each channel was used sequentially as an independent experimental repeat. Using a Silicone Elastomer Kit (Electron Microscopy Sciences, 24236-10, Sylgard™ 184, USA,) the Silicone Elastomer Base was added to the Silicone Elastomer Curing Agent (10:1 ratio) to prepare polydimethylsiloxane (PDMS) according to the manufacture’s protocol. To remove air bubbles, the mixture was centrifuged at 200 g for 15 min, and poured into a mold to prepare a PDMS layer 6 – 8 mm in height. The mixture was covered and incubated at room temperature for 20 – 25 min to remove small bubbles, then placed in a thermostat at 75°C for 2 h. Solidified PDMS was carefully cut from the mold using a scalpel, and vertical holes (diameter 1 mm) were made at the ends of each channel using a puncher to serve as sites for tubing connections.

#### Flow chamber assembly and functionalization

Glass coverslips (VWR, 48366-089, USA) and PDMS slabs were cleaned by rinsing with isopropanol and then milliQ water, dried and treated for 30 s with oxygen plasma using a Harrick Plasma Cleaner (Harrick Plasma, USA) at 400 mTorr. Immediately after plasma cleaning, one treated coverslip was attached to the PDMS slab to form a tight seal. To form a channel, two 21G needles were connected to polyethylene tubes (Medsil, TMPE-9, Russian Federation). Needle tips were smoothed by manual filing and carefully inserted into the holes in the PDMS slab. The free end of each “inlet” tube (length 6 cm) was placed into the Eppendorf tubes with solutions, whereas the “outlet” tubes (length 75 cm) were connected with a 3 ml syringe, which was used to manually withdraw solutions from the chamber (Supplementary Figure 1A). After the flow chamber was assembled, the coverslip was functionalized by drawing in the fibrinogen solution or WM59 antibodies and incubating for 1 h at room temperature. The channels were washed by perfusing each with 0.5 ml PBS for 5 min, and the surfaces were then passivated by incubation with Buffer A supplemented with 4% BSA for 20 min. Flow chamber was placed in a custom-made holder (Supplementary Figure 1A), which was printed at the 3D printing service at Biotech Commons, University of Pennsylvania, USA. The fully assembled chamber was placed on a microscope stage and used immediately to study platelets activation.

### Microscopy procedures

Images of mepacrine-stained DGs were acquired in epifluorescence mode with an Axio Observer.Z1 microscope (Carl Zeiss, Germany) equipped with a Yokogawa spinning disc confocal device (CSU-X1; Yokogawa Corporation of America, USA), QuantEM camera (Photometrics, USA) and 100×oil objective. Mepacrine excitation was achieved with a 385 nm diode, exposure time 125 ms. Images were acquired every 10 s for 10 min. Iris diaphragm was partially closed to limit illumination only by the imaging field. Simultaneous imaging of DGs and calcium was carried out analogously but using a Nikon Eclipse Ti-E inverted microscope, see Chakraborty et al.^80^ for details. Mepacrine and Calbryte-590^AM^ were excited by rapidly switching between 488 nm and 561 nm lasers (100 mW Sapphire lasers by Coherent, USA). No crosstalk was observed between the fluorescence channels for cells stained with these dyes. The exposure time for each laser was 50 ms at 2 frames per second. Mepacrine brightness data obtained with the Zeiss microscope are shown in Figure 2, whereas all other brightness measurements were acquired using the Nikon microscope. High resolution imaging was carried out using Olympus IX71 inverted microscope (Olympus Corporation, Japan) equipped with Visitech VT iSIM scan head (VisiTech International, UK), Hamamatsu ORCA Quest qCMOS camera (Hamamatsu Photonics, Japan). Mepacrine excitation was achieved with a 488 nm laser, exposure time 200 ms. Z-stacks were acquired with 150 nm steps.

### Fluorescence-based DG secretion assay and calcium visualization

On the day of experiment, blood was collected in S-Monovette tubes (Sarstedt, 05.1071.100, USA) using 3.2% (w/v) sodium citrate as an anticoagulant. All preparations and experiments were done at room temperature. To ensure reproducibility, blood was stored for no longer than 0.5 – 1.5 h after collection. To stain DGs, blood was incubated for 20 min with 10 μM mepacrine. In chambers with fibrinogen-coated coverslips, blood was perfused at 1,000 s^-1^ shear rate for 20 s by withdrawing the liquid with a syringe pump (New Era, NE-4000, USA). Platelets attachment was confirmed using a differential interference contrast (DIC). Experiments with WM59 antibodies used reduced shear stress of 200 s^-1^ to ensure stable platelet attachment on WM59-coated surface. The chamber with attached platelets was washed twice by perfusing buffer A supplemented with 1% BSA and 1 μM mepacrine for 13 min. Fluorescent imaging started 1 min before platelet activation and continued for 11 min. DIC images of platelets were collected before and after platelet activation. Platelet activation was induced by the addition of buffer A with 2 mM CaCl_2_, 1% BSA, μM mepacrine and indicated activators. It took 20 s for an activator to flow through connecting tubes and reach cells, as determined in control experiments with a fluorescent dye (Annexin V-Alexa^647^ or CD62P-Alexa^647^). Time of activator’s contact with cells is indicated as 0 time on all graphs, which we refer as the time of activator addition.

To examine intracellular calcium levels, blood was incubated with 10 μM mepacrine and 1 μM calcium-specific dye Calbryte-590^AM^ for 20 min, and platelets were prepared as above. Extracellular calcium was chelated by addition of EGTA. Briefly, blood was perfused through a channel and adhered platelets were washed for 13 min with buffer A supplemented with 1% BSA, 1 μM mepacrine, 5 mM EGTA and 5 mM MgCl_2_. Platelet activation was induced by the addition of buffer A supplemented with 1% BSA, 1 μM mepacrine, 5 mM EGTA, 5 mM MgCl_2_ and 0.1 U/ml thrombin.

### DG secretion in the presence of different blood preparations

“Secreted content” (SC) was obtained by centrifuging freshly drawn citrated blood at 100 g for 8 min. Platelet rich plasma was centrifuged at 400 g for 5 min and 10 μl of plasma was used for control activation. The platelet pellet was resuspended in Buffer A supplemented with 1% BSA, cells were activated by incubating with 0.1 U/ml thrombin and 2 mM CaCl_2_ for 5 min, centrifuged at 400 g for 5 min, and supernatant was collected, corresponding to SC. Supernatant from unactivated platelets was prepared analogously, except the platelet pellet was incubated in Buffer A with 1% BSA for 5 min.

### Experimental data analysis

Images were analyzed with ImageJ (National Institute of Health, USA). Data was analyzed using Origin (OriginLab Corporation, USA), and Prizm (GraphPad Software, USA) or Matlab R2014a (MathWorks, USA) programs. Whenever possible, data are presented for *N* independent trials, i.e. experiments carried out in separate channels of a flow chamber. Each experiment used at least 3 different donors on at least 3 different days. In figure legends, small *n* denotes the total number of analyzed platelets in all trials for indicated conditions. Unless stated otherwise, the mean of values obtained in independent trials are reported as mean ± SEM.

#### Quantification of the number of DGs in human platelets

For SIM-generated images, Z-stacks were deconvoluted using ImageJ Microvolution plugin and examined in ImageJ. Only single platelets that did not move significantly during acquisition were analyzed. Spots were counted as DGs regardless of their brightness if they were consistently visible across multiple planes and had round or oval shape. Dumbbell-shaped spots were counted as two granules, and three-lobed spots as three granules. Because several granules may be located close together without changing round shape, the reported DG number may be an underestimation.

#### Quantification of the kinetics of DG secretion in single human platelets

To determine kinetics of DG secretion, approximately 10 platelets were selected from the imaging sequence representing one independent experiments. The cells were selected based on the following criteria: cells were fully present in the field of view; they did not overlap and did not move significantly during the experiment. Platelet images collected using regular epifluorescence microscopy were analyzed based on the total cell brightness. A central area of each cell containing all mepacrine-stained dots was selected using a hand drawn region with a minimal area to reduce background (Supplementary Figure 1B), and the integrated intensity was collected over time. The same region was then shifted to an area adjacent to the cell to collect integrated background intensity. To reduce noise, each curve for changes in background intensity was smoothed using the Savitzky-Golay method^81^ with a sliding window of 100 points. The resultant background intensity curve was subtracted from the integrated intensity curve of the corresponding cell to generate the mepacrine intensity curve for each cell, representing the total pool of DGs. The intensity curves from multiple cells were further analyzed using a custom written MATLAB program “Quantitative analysis of kinetics of DG secretion and calcium oscillations in individual platelets” (Supplementary Figure 3B, see Data Source File), which is available at https://www.med.upenn.edu/grishchuklab/protocols-software.html. Briefly, the release events were identified by an abrupt drop in mepacrine fluorescence. Each release event was confirmed visually using the corresponding imaging series to avoid false events. The duration of a release event was determined as a time between the frame immediately prior to the fluorescence drop and the frame when the fluorescence plateaued. Time intervals between the activator’s addition (time 0) and the 1^st^ release event, as well as the intervals between subsequent release events, were collected. Their frequency distributions were fitted using the exponential function 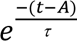, where *t* is experimental time, τ is characteristic time and *A* is time to a maximum. The value of parameter A was used to determine the “lag time” for secretion. The percent of secreted granules during each release event was calculated by dividing the change in cell brightness during the event by the cell brightness prior to this event.

To generate population secretion curves representing platelets behavior for specific experimental conditions, the mepacrine intensity curves for cells from one independent experiment were averaged. The resultant curves for *N* independent experiments carried under identical conditions were averaged and plotted with SEM. To compare population secretion curves for different agonist concentrations, the curves were normalized based on the maximum intensity at time zero. To obtain the cumulative frequency of platelets undergoing a 1^st^ release event, we calculated the percent of platelets that exhibited their 1^st^ release event by each time point.

#### Quantitative analysis of irregular calcium oscillations in individual platelets and testing of “null hypothesis”

To assess the dynamic level of calcium signal in individual cells, the intensity of Calbryte-590^AM^ was measured over time inside the cell. Due to the fact that platelets have irregular shapes and spread upon activation, we needed a standardized approach for quantification. Therefore, for each platelet, a circular region that included the maximum number of bright pixels was selected to measure the intensity of Calbryte-590^AM^ over time (Supplementary Figure 6A), which we refer to as the Calbryte intensity curve. The temporal sequence of calcium spikes was analyzed using a custom written MATLAB program “Quantitative analysis of kinetics of DG secretion and calcium oscillations in individual platelets” (Supplementary Figure 7A). First, for each cell, a baseline curve was determined as following. On the Calbryte intensity curve, six points were manually selected between calcium spikes. These six dots were fitted linearly by the program, creating the baseline curve. The baseline curve was then subtracted from the Calbryte intensity curve. Next, the “cut-off” level was visually determined above the baseline curve but below the peaks of the calcium spikes. The time moments of the calcium spikes were determined using local maximum condition. The “time to the preceding spike” was defined as the time difference between the preceding calcium spike and the start of the 1st release event.

To test the “null hypothesis” that the closeness of calcium spikes to release events is not coincidental, a Bootstrap simulation was performed. The MATLAB program “Quantitative analysis of kinetics of DG secretion and calcium oscillations in individual platelets” selected 100 random time points in the Calbryte intensity curve using a homogeneous probability density function and calculated the “time to the preceding spike” between each random time point and the preceding calcium spike.

#### Quantitative analysis of calcium oscillations in relation to DG secretion

To investigate whether the spikes that preceded secretion had any special features that may control timing of secretion or number of secreted DGs, two distinct data groups based on their timing relative to the 1^st^ DG release event were created. The first group included data for the time interval from the activator addition until the 1^st^ release event. The second group included data for the specific time interval prior to the beginning of the 1^st^ release event. The specific time intervals chosen are described below and in Supplementary Figure 7B-D.

#### Correlation of timing of release events with calcium spike amplitudes

To test a hypothesis that a calcium spike of a specific amplitude induces release of DGs we quantified calcium spikes amplitudes of two data groups described above. Using the MATLAB program “Quantitative analysis of DG secretion and calcium oscillations in platelet population” the amplitudes of calcium spikes were defined the spike’s maximum. The amplitudes of calcium spikes varied among different platelets, presumably because of variations in platelet volume and the amount of Calbryte-590^AM^ loaded per cell. To overcome this difficulty, we normalized the amplitude of each calcium spike by dividing it by the average value of all spike amplitudes before the 1^st^ release event. This average value was determined independently for each platelet:

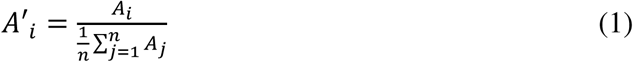

where A′_i_ is the relative amplitude of spike number *i* following platelet activation, A_i_ is the absolute value of spike amplitude derived from the Calbryte intensity curve, and the denominator corresponds to the average value of absolute spike amplitudes before the release event.

Then, we combined data for the platelet population and compared relative amplitudes of calcium spikes that preceded the 1^st^ release event, with relative amplitudes of all calcium spikes that occurred before the 1^st^ release event (Supplementary Figure 7B).

#### Correlation of timing of release events with intervals between calcium spikes

In order to examine whether the release event occurred following decreased intervals between calcium spikes, we compared the distributions of time intervals between calcium spikes of two data groups described above. Using the MATLAB program “Quantitative analysis of DG secretion and calcium oscillations in platelet population” intervals between calcium spikes were defined between spikes’ maximums (Supplementary Figure 7C). The first interval distribution was plotted for the intervals between the two spikes preceding the 1^st^ release events. The second interval distribution was plotted for the intervals between all spikes that occurred before the 1^st^ release event. Lastly, we quantified intervals between calcium spikes after the 1^st^ release event to determine whether DG release quickly increases calcium spike frequency of the same platelet.

#### Correlation of brightness of released DGs with calcium spike amplitudes and cumulative calcium signal

To investigate whether brightness of secreted DGs depends on amplitude of the preceding calcium spike, we plotted brightness of released DGs in the 1^st^ release event against the relative amplitude of the preceding calcium spike (Supplementary Figure 7B). The correlation was analyzed using Pearson correlation analysis built in Prizm. The analysis revealed absence of any dependence of brightness of released DGs in the 1^st^ release event on the amplitude of the preceding calcium spike. To investigate whether the brightness of released DGs depends on cumulative calcium signal preceding the release event, we measured the relative area under the Calbryte intensity curve per cell prior to the DG release event:

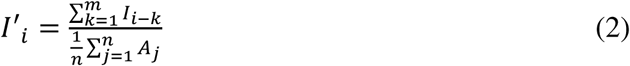

where the numerator represents the sum over a specific number of time frames (*m*) of the Calbryte intensity curve (I_i-k_). These time frames correspond to the measurements preceding the time frame (*i*) for which the relative calcium integral I′_i_ was calculated. The denominator represents the average value of the absolute calcium spike intensities before the DG release event. This normalization was used to compare results obtained for different platelets.

The correlation between brightness of released DGs during the 1^st^ release event and the area under the Calbryte intensity curve during the 10 s time interval was estimated using Pearson correlation analysis built in Prizm.

#### Theoretical modeling

Theoretical approaches are described in Section 1 of Supplementary Materials. Part 1 describes our probabilistic model (written in Python) of calcium spiking and DG secretion, whereas Part 2 describes a multi-scale model of thrombus formation in blood flow, which was implemented using C++ language (OpenFOAM).

## Data availability

A Data Source file containing the quantitative data underlying all graphs is provided with the Supplementary Materials. Due to their large size, raw imaging datasets (time-lapse microscopy sequences) are not included but are available from the corresponding author upon reasonable request.

## Code availability

The algorithms used for computer simulations of secretion are described in Section 1 of the Supplementary Materials. The source codes and associated files are available at https://gitlab.com/masaltseva/particle-thrombosis.

## Acknowledgements

We gratefully acknowledge the invaluable input from D. Yu. Nechipurenko, who helped with computer modelling and contributed to discussions. We thank Anna Parkhaeva for technical assistance, Dr. W. Luo and other Grishchuk lab members for discussions and assistance. We are grateful to Drs. M.S. Marks and N. Chanaday for critical reading of the manuscript, and to members of the Upenn Platelet Club and Dr. K. Foskett for insightful discussions. This work was supported by the Perelman School of Medicine at the University of Pennsylvania through a discretionary fund awarded to ELG. Masks for flow chambers were prepared at Singh Center for Nanotechnology (University of Pennsylvania, USA), which is supported by the NSF National Nanotechnology Coordinated Infrastructure Program under grant NNCI-2025608, and chamber holders were made at 3D printing service at Biotech Commons (University of Pennsylvania, USA).

## Authorship contributions

T.O.S performed all experiments with platelets and analyzed data, A.A.M and F.I.A carried out mathematical modeling, R.R.K assisted with data analysis and investigated secretion in cells immobilized with PECAM-1 antibodies, T.O.S, F.I.A. and E.L.G interpreted data and designed research, T.O.S. and E.L.G. wrote the manuscript with input from all authors.

