## Supplement for "Burst-like Secretion of Platelet Dense Granules Promotes Thrombus Shell Expansion"

### TABLE OF CONTENT


1. **Theoretical approaches**

**Part 1. Modeling of DG secretion in platelets**

1.1. Model framework
1.2. General description of platelets and DGs
1.3. Calcium-induced DG maturation and secretion
  1.3.1. Description of calcium spikes
  1.3.2. DG maturation
  1.3.3. Calcium-induced triggering of the DG release event
  1.3.4. Release of DGs during a calcium-induced secretion event
1.4. Modeling of platelets activators
1.5. Numerical simulation algorithm
1.6. Model calibration
  1.6.1. Parameters that characterize secretion in response to thrombin
  1.6.2. Parameters that characterize secretion in response to secreted agonists
1.7. Verification of the developed model of bursting DG release during a calcium-triggered secretion event
1.8. Modeling algorithm, calibration and verification for the random secretion scenario
1.9. Modeling platelet activation in the presence of the secreted content (SC)

**Part 2. Modeling of thrombus formation**

1.10. General approach
1.11. Model description
  1.11.1. Model geometry
  1.11.2. Modeling blood hydrodynamics
  1.11.3. Modeling soluble platelet activators
  1.11.4. Modeling platelet interactions
  1.11.5. Modeling DG secretion
    1.11.5.1. DG secretion in response to calcium spiking
    1.11.5.2. Modeling different scenarios of secretion in the growing thrombus
1.12. Numerical simulations of thrombus formation
1.13. Processing of Model-Generated Data for Visualization and Quantification

2. **Supplementary Notes**

Supplementary Note 1. Quantification of DG Count in Human Platelets.
Supplementary Note 2. Spatial DG Distribution Does Not Govern Temporal Secretion Coordination.
Supplementary Note 3. Neither Calcium Spike Amplitude nor Frequency Determines DG Release Timing.
Supplementary Note 4. Estimating ATP, ADP, and Serotonin Concentrations in Secreted Content.

3. **Supplementary Tables**
4. **Supplementary Figures and Figure Legends**
5. **Supplementary Movie Legends**
6. **Supplementary References**

### 1. Theoretical approaches

To seek mechanistic insights into our experimental results on platelet activation, we used mathematical modeling. Part 1 of this section describes our model of calcium-induced DG secretion in platelets. Part 2 describes a combined model, which incorporates the above model of DG secretion at a single cell level and our previously developed model of thrombus formation in a blood flow[1].

#### Part 1. Modeling of DG secretion in platelets

#### 1.1. Model framework

We use a stochastic approach to provide a simplified description of the kinetics of DG release. Platelets are modeled as virtual spherical objects characterized by their radius, a set of DGs and intracellular calcium concentration. The model does not consider the internal structure of a cell, including spatial distribution of DGs, or biochemical pathways that lead to generation of calcium spikes and DG release. In the model, platelets exist in two states: resting and activated. In the resting state, the DGs are not secreted and calcium concentration is low. Transition from this initial state into the activated state is triggered by various activators. In the activated state, the platelets generate calcium spikes, which induce DG maturation and stochastic secretion of DGs. The calcium spike frequency, DG maturation and secretion are described using the concentration-dependent functions for thrombin and ADP. The value of parameters for these functions are determined by calibration based on our experimental findings. As the modeling output, we calculate the number of DGs secreted over time as a function of activator concentration. Furthermore, the model allows to examine kinetics of DG release in response to a combination of thrombin and other activators.

#### 1.2. General description of platelets and DGs

Because real platelets differ in size, our model assumes that platelet radius $r$ is varied. For simplicity, $r$ is distributed normally:

$$r = |r_o + r_\sigma \zeta|,\ \zeta \sim [0,1] \qquad (3)$$

where $r_o$ is the mean platelet size, $r_\sigma$ is the width of size distribution and $\zeta$ is a random number within the range from 0 to 1.

Values of parameters $r_o$ and $r_\sigma$ are chosen to reproduce the experimental distribution of platelet volumes in ref[2]: $r_o$ = 1 µm and $r_\sigma = 0.22$ µm (Supplementary Figure 9E).

We note that the distribution of the number of intracellular DGs in the model (Supplementary Figure 9E) is similar to the published distribution of platelet volume in ref[2] (Supplementary Figure 9E). For this reason, the number of DGs in a resting platelet, $N_{dg}$, in our model is assumed to be proportional to cell volume:

$$N_{dg} = N_o \left(\frac{r}{r_o}\right)^3 \qquad (4)$$

where $N_o$ is the factor that characterizes the number of DGs in unactivated platelets. In our model, this unitless normalizing factor $N_o$ = 19. With this value, the model described well the experimental intensity distribution of the minimally-sized DGs with average brightness 30,000 a.u. (Supplementary Figure 9E, Supplementary Note 1).

### 1.3. Calcium-induced DG maturation and secretion

#### 1.3.1 Description of calcium spikes

The addition of activators in the model induces the activated state and leads to calcium spikes, i.e. the transient peaks in intracellular calcium concentration. In the model, calcium ions are not described explicitly, and all spikes have the same amplitude. The latter simplification is justified by our finding that the spike's amplitude does not correlate with the occurrence of secretion events (Supplementary Figure 7B). For simplicity, the duration of each spike is assumed to equal 1time step $\Delta t$. The timing of spikes is stochastic and takes place with probability $p_{Ca}$, which is linked to the experimentally measured calcium spike frequency $f$: $p_{Ca} = \Delta t f$. In the model, the dependency of $p_{Ca}$ on thrombin concentration $T$ is described with the following expression:

$$p_{Ca}(T) = \Delta t\, a(b + \log T) \tag{5}$$

where $a$ characterizes the slope of the dependency and $b$ is the probability of calcium oscillations in resting platelets. Our experimentally determined function for the frequency of spikes in response to thrombin concentration is fitted using Wolfram Mathematica, Version 12 (Wolfram Research, USA). With $a$ = 1.38 min$^{-1}$ and $b$ = 9.79, Equation (5) provides a good fit (Supplementary Figure 9A).

In our experiments, calcium spikes can also be triggered in platelets by the addition of ADP, but the frequency of spikes is less sensitive to ADP concentration than thrombin (Supplementary Figure 9B). We therefore assume that probability of calcium spikes in the presence of ADP, $p_{Ca}(ADP)$, is constant for the range of examined ADP concentrations. We estimate this probability from the average frequency of calcium spikes in platelets activated with 1 µM and 100 µM ADP:

$$p_{Ca}(ADP) = 2.2 \cdot 10^{-4} \tag{6}$$

#### 1.3.2 DG maturation

DG maturation is modeled as a stochastic, two-state process with maturation probability $p_m$: each DG is either immature or mature, and only mature granules can be secreted. In non-activated cells $p_m$= 0. After addition of an activator, $p_m$ gradually increases. This gradual rise was included because experimentally we observed a lag period before the onset of DG secretion. In the model, $p_m$ eventually reaches a plateau (Supplementary Figure 9F). The plateau is justified by the observed constant probability of DG secretion across different release events once the lag period has ended.

To define this $p_m$ function quantitatively for a general case, we considered that the lag time—the interval between activation and the first secretion event—depends on the type and concentration of the activator (Figure 4E). However, the number of calcium spikes during the lag period shows little dependence on activator type or concentration (Figure 4F). Therefore, we modeled the change in $p_m$ during the lag period as a function of the number of calcium spikes rather than explicit time, which allows for a universal formulation of the maturation probability after n calcium spikes:

$$p_m = \frac{1}{1+\left(\frac{n_{lag}}{n}\right)^3} \tag{7}$$

Here, $n_{lag}$ is a constant that defines the number of calcium spikes required for $p_m$ to reach its half-maximum value. The choice of $n_{lag}$ is discussed in Section 1.6.1.

#### 1.3.3 Calcium-induced triggering of DG release events

Experiments reported here indicate that DG release events occur following calcium spikes. To model DG release specifically after a calcium spike, we assumed that activated platelets can secrete mature DGs only during a brief period called the “window of opportunity.” This window opens immediately after a calcium spike and, in the model, lasts for $t_{Ca}$ = 3 s. This duration was chosen because in our experiments, approximately 95% of platelets secreted DGs within 3 s after a calcium spike (Figure 4D).

Experimental data also indicate that calcium spikes trigger DG release stochastically. To represent this, each calcium spike in the model induces a window of opportunity during which any mature DG can be secreted with probability $\pi$ during one modeling time step. When the window of opportunity closes, $\pi$ = 0.

Thus, two independent stochastic processes determine the probability $p_1^i$ that a given DG becomes secreted in an activated cell: the probability $p_m$ that this DG has transitioned to the mature state, and the probability $\pi^i$ that the mature DG becomes secreted during the calcium-induced window of opportunity:

$$p_1^i = p_m \pi^i \tag{8}$$

where superscript $i$ corresponds to the DG index (1 to $N$), and $N$ is the total number of DGs in the modeled cell.

Finally, we took into account that not all DGs are released within the observation time of 600 s (Supplementary Figure 5G). This incomplete release is modeled by assuming that the probability $\pi^i$ for a mature DG to become secreted varies among granules. This variation is described mathematically using the following distribution for $\pi^i$ (Supplementary Figure 9G):

$$\pi^i = \frac{1-\sigma^4}{1+\left(\frac{\sigma}{\xi^i}\right)^4} \tag{9}$$

Here, parameter $\xi^i$ is uniformly distributed between 0 and 1. Each DG in the resting cell is randomly assigned its own $\xi^i$, which remains constant throughout the simulation, leading to different values of $\pi^i$ among DGs. DGs with low $\xi^i$ have correspondingly low secretion probabilities $p_1^i$. Parameter $\sigma$ in Equation (9) is a scaling factor that modulates DG release probability in response to different activators, as described in Section 1.6. Smaller values of $\sigma$ correspond to higher secretion probabilities in the modeled cell.

At the population level, equation (9) yields a residual fraction of non-secreted DGs, consistent with experimental observations (Figure 6B, Supplementary Figure 10B).

To summarize, the overall probability for one DG to become secreted, $p_1^i$, is given by:

$$p_1^i = p_m \pi^i = \frac{1}{1+\left(\frac{n_{lag}}{n}\right)^3} \cdot \frac{1-\sigma^4}{1+\left(\frac{\sigma}{\xi^i}\right)^4} \tag{10}$$

#### 1.3.4 Release of DGs during a calcium-induced secretion event

We consider two scenarios of DG release during a calcium-induced secretion event: bursting granule release and random granule release. In both scenarios, secretion of the first DG, by definition, corresponds to the start of the release event. Therefore, for both scenarios, the probability of the secretion event equals the probability of the secretion of the first DG.

In the random release scenario, after a calcium spike opens the window of opportunity, all DGs can be secreted according to their maturation-dependent probabilities $p_1^i$, as in Equation (10).

In the bursting release scenario, positive feedback is initiated after the first DG is released. For simplicity, we assume that this positive feedback acts on all remaining DGs ( $p_m$ = 1). Owing to this feedback, the probability of the release of the remaining DGs, $p_2^i$ , is larger than probability of the release of the first DG by a feedback parameter $K > 1$:

$$p_2^i = K p_1^i \tag{11}$$

The choice of $K$ value is discussed in Section 1.6.

Because the first DG is released stochastically during the calcium-induced time window, this release may happen late during the window of opportunity, thereby reducing the feedback effect. To compensate for this effect, after the first DG is secreted the window of opportunity is extended by $t_K$ = 3 s, during which all remaining DGs can be secreted with the probability $p_2^i$, as in Equation (11).

**1.4. Modeling of platelet activators**

In our model, different activators regulate DG secretion in two distinct ways. First, the activators initiate calcium spikes, as described in Section 1.3.1. Second, the activators modulate the probabilities of DG release during the calcium-induced window of opportunity. This is achieved mathematically via parameter $\sigma$ , which adjusts the release probabilities of the first DG ($p_1$) and the remaining DGs ($p_2$) in accordance with Equations (10) and (11). The value of parameter $\sigma$ in different scenarios was determined via model calibration procedure, see Section 1.6.

**1.5. Numerical simulation algorithm**

Stochastic simulations of calcium spike generation and DG release are carried out using the custom script written in Python v3.0 (Python Software Foundation, USA), which is freely available at https://www.med.upenn.edu/grishchuklab/protocols-software.html. For all simulations, the time step $\Delta t$ is 2 ms and the total simulation time is 600 s.

The following stochastic simulation algorithm was implemented:

1. In a resting platelet, the initial number of DGs is generated and individual DGs are assigned their randomly generated values of parameter $\xi^i$. These values do not change during the simulation of this platelet.

2. After addition of an activator at $t = 0$, calcium spikes are generated with the probability $p_{Ca}$, which depends on the type and concentration of an activator. Specifically, at each time step a random number $\hat{p}$ from the range [0, 1] is chosen. If $\hat{p}$ > $p_{Ca}$, a spike is generated, a calcium-induced window of opportunity opens, and the algorithm proceeded to step 3. If $\hat{p} \leq p_{Ca}$, a spike is not produced. If the window of opportunity induced by the previous spike is still open, the algorithm proceeded to step 3. If there are no open windows from any prior spikes, the algorithm proceeded to step 5.
3. In the bursting secretion scenario, the DG release during the opened window is calculated as follows.

a. At each time step, the program evaluates whether any DGs have already been released during this opened window. If there have been no released DGs, probability $p_1^i$ is calculated for each DG using Equation (10). For the DG with the largest $p_1^i$, a random number $\hat{p}$ from the range [0, 1] is generated. If $\hat{p} > p_1^i$, the DG is not released at this time step, and the algorithm proceeds to step 4. If $\hat{p} < p_1^i$, this DG is released, and the window of opportunity is extended by the time interval $t_K$, as described in Section 1.3.4, and the algorithm proceeded to step 4.

b. If a DG has been released during the window of opportunity before this time step, probability $p_2^i$ is calculated for all remaining DGs in the platelet in accordance with Equation (11). For the DG with the largest $p_2^i$, a random number $\hat{p}$ from the range [0, 1] is generated. If $\hat{p} > p_2^i$, the DG is not released at this time step, and algorithm proceeds to step 4. If $\hat{p} < p_2^i$, the DG is released. If the original calcium-induced window of opportunity has lapsed, the window of opportunity is extended by the time interval $t_K$, as described in Section 1.3.4, and the algorithm proceeded to step 4.

4. Calculations at subsequent time steps proceed as in step 3 until the window of opportunity and its extensions close. The algorithm proceeds to step 5.

5. Steps 2-5 are repeated until the end of simulation for this cell.

As the outcome, for each platelet the program calculates the number of DGs recorded at each time step, the number of DGs released during different events, the number of spikes before the 1st release event and time intervals for all events. Calculations were carried out for $n$ = 100 cells for different types and concentrations of activators. The total number of DGs at each time step for all cells was normalized to the initial number of DGs in these cells and plotted as a function of time to obtain a population secretion curve (as in Figure 6B).

#### 1.6. Model calibration

The above-described model for bursting secretion has 15 parameters, which are listed in Table 1. The values of 8 parameters: $r_o$, $r_\sigma$, $N_o$, $a$, $b$, $p_{Ca}(ADP)$, $t_{Ca}$ and $t_K$ were chosen based on the published results or the experimental findings reported in this work, as explained in Sections 1.2, 2.3.1 and 2.3.3-4. However, values of the remaining 7 parameters could not be derived in this way. Therefore, these parameters were determined by model calibration using experimental data, as described below.

##### 1.6.1 Parameters that characterize secretion in response to thrombin

This section describes the calibration procedure to determine the feedback parameter $K$, parameter $n_{lag}(T)$ and parametric function $\sigma$ from Equations (10) and (11) for different thrombin concentrations. Parameters $K$ and $n_{lag}$ do not depend on thrombin concentration, so their values were determined by fitting the population secretion curve in response to thrombin 0.1 U/ml (Figure 6B). The same fitting also defines parametric function $\sigma$ for this thrombin concentration. Values $K$ = 7.7647, $n_{lag}(T)$ = 7 and $\sigma$ = 0.7 provide a good fit of the experimental data for 0.1 U/ml thrombin activation (Figure 6B, Supplementary Figure 10A).

To determine the parametric function $\sigma(T)$ across the thrombin concentration range, we fitted the model to recapitulate the experimental population secretion curves (Supplementary Figure 9H). Smoothing of $\sigma(T)$ was achieved by extrapolation with exponential function (Supplementary Figure 9J):

$$\sigma(T) = a_1\, e^{-(b_1 T)^{0.5}} \quad (12)$$

where $a_1$ = 0.851 and $b_1$ = 0.26 ml/U.

**1.6.2 Parameters that characterize secretion in response to ADP**

Calibration procedure to determine the values of parameters $K$, $n_{lag}$ and $\sigma$ (eq. 10) for different ADP concentrations was carried out analogously to thrombin. Parameters $K$ and $n_{lag}$ do not depend on ADP concentration, so their values were determined by fitting the population secretion curves for ADP activation, see Supplementary Figure 5G. For all concentrations of ADP, $n_{lag}(ADP)$ = 0 and $K$ = 7.7647 provided the best description of the population secretion curves (Supplementary Figure 9I). Smooth function $\sigma(ADP)$ was determined by extrapolation (Supplementary Figure 9J):

$$\sigma(ADP) = a_2\, e^{-(b_2 ADP)^{0.2}} \quad (13)$$

where $a_2 = 1$ and $b_2 = 2.4 \cdot 10^{-5}\mu M^{-1}$.

**1.7. Verification of the developed model of bursting DG release during a calcium-triggered secretion event**

After establishing the model framework and determining all parameter values, we tested the model by using it to predict the outcomes of experiments that have not been used during model calibration. The values of all model parameters were fixed (Supplementary Table 1), ensuring unbiased and rigorous model testing. Unless stated otherwise, calculations for each condition were carried out for *n* = 100 platelets using the algorithm described in Section 1.5.

1) The model was used to determine the apparent probability $p_s$ of the 1st release event for a range of thrombin concentration ($10^{-3}$ – 1 U/ml). This probability was calculated as $p_s = 1/n_s$, where $n_s$ is the number of spikes preceding the 1st release event. The model correctly predicted that probability of the 1st release event is largely insensitive to thrombin concentration (Supplementary Figure 9K), as in our experiments (Figure 4F). The mean value of $p_s$ for all thrombin concentrations (0.103 ± 0.04) is in agreement with the experimentally determined probability 0.11 ± 0.04 for the same conditions (Figure 4F).

2) The model was used to calculate parameters of DG release which were not taken into account during model training. A. The simulated distribution of the number of remaining granules at 300 s after the activation by 0.1 U/ml thrombin is in agreement with the experimental result (Supplementary Figure 10B). B. Average time intervals between the activator's addition and the 1st release event, and between the subsequent events in the model are also similar to analogous experimental results (Supplementary Figure 10C). C. The average number of the released DGs per event in the model shows a declining trend, which is similar to experimental results (Supplementary Figure 10D). However, in the model, the percent of released DGs during subsequent events is declining, whereas in our experiments it is approximately constant (Supplementary Figure 10E). This minor discrepancy between the model and experiment suggests that the framework of our simplified model may need further refinement.

3) The model was used to calculate DG release in response to four concentrations of thrombin: $10^{-3}$, $10^{-2}$, 0.1, 1 U/ml. We performed numerical simulations for *n* = 100 platelets for each concentration. Following output characteristics were determined: the percent of secreting platelets, the percent of secreted DGs during entire simulation time, time before the 1st release event, time

between the 1st and 2nd release events, total number of release events, and the number of released DGs during the 1st event. Theoretical results show the same trends as observed in the experiments (Supplementary Figure 11): the percent of secreting platelets (Supplementary Figure 11A), percent of released DGs (Supplementary Figure 11B), total number of release events and the number of DGs in the 1st release event (Supplementary Figure 11E,F) increase with increasing thrombin concentration, whereas the time before the 1st release (Supplementary Figure 11C) and time between the 1st and the 2nd release (Supplementary Figure 11D) decrease with increasing thrombin concentration.

4) The model was used to calculate DG release in response to five concentrations of ADP ($10^{-2}$, 0.1, 1, 10, $10^{-2}$ μM), using the same approach as described for thrombin. The results in Supplementary Figure 11 show the same trends as observed in experiments. The percent of secreting platelets (Supplementary Figure 11G), percent of secreted DGs (Supplementary Figure 11H), total number of release events the and number of DGs in the 1st release (Supplementary Figure 11K,L) show increase with increasing ADP concentration, whereas the time before the 1st release decreases with increasing ADP concentration (Supplementary Figure 11I). Several minor discrepancies between the model and experiment were noted. First, the model predicts that the time between the 1st and 2nd release events should remain constant, whereas in our experiment this time decreases slightly and within the margin of error in response to increasing ADP concentration (Supplementary Figure 11J). Second, the number of released DGs in the 1st release event slightly increases with increasing ADP concentration in the simulations, but it remains fairly constant in the experiment (Supplementary Figure 11K), suggesting that the future refined model should improve the definition of probability $p_2^i$. Since at each ADP concentration the number of DGs predicted by the model is close to experimental values, we considered this discrepancy minor and did not pursue further model refinements.

Overall, modeling results for four different experimental approaches, each of which provides a rigorous and independent test for the model, show robust consistency, thereby verifying the model and the chosen parameter values.

#### 1.8. Modeling algorithm, calibration and verification for the random secretion scenario

The modeling algorithm, calcium triggering, and all model parameters in this scenario were identical to those in the bursting secretion model, except for the description of DG release during the calcium-induced window of opportunity. Specially, in the random secretion scenario, positive feedback was excluded by setting the feedback parameter K = 1 (Eq. 11). As a result, when calcium spike triggers the window of opportunity, all DGs have an equal probability of being released.

To implement this scenario, the calculation algorithm described in Section 1.5 was modified by replacing step 3 with the following clause, which mathematically defines the incorporation of randomness:

"The probability $p_1^i$ is calculated for each DG using Equation (10). For the DG with the largest $p_1^i$, a random number $\hat{p}$ is drawn from the range [0, 1]. If $\hat{p}$ exceeds $p_1^i$, the DG is not released at this time step, and the algorithm proceeds to step 4. Otherwise, if $\hat{p}$ is smaller than $p_1^i$, the DG is released, and the algorithm continues to step 4."

Importantly, this modeling scenario was calibrated using the same experimental data as the bursting secretion scenario (see Section 1.6.1). Specifically, the DG release probability was set based on $\sigma_{rand}$ = 0.8, which ensured a good fit to the population secretion curve at 0.1 U/ml thrombin (Figure 6B). In the bursting scenario, the probability of DG release decreases at lower thrombin concentrations. When a similar dependency was applied to the random secretion scenario, the model failed to adequately fit the experimental results. To address this limitation, we explored an alternative modification in which the DG release probability remained constant, regardless of thrombin concentration, and was maintained at the same high level observed at 0.1 U/ml thrombin. However, as described in the main text, even with this increased secretion sensitivity, the model still failed to support robust shell expansion in silico.

#### 1.9. Modeling platelet activation in the presence of the secreted content (SC)

The model was applied to describe experiments, in which platelets were activated by a combination of 0.1 U/ml thrombin and the SC. As described in the main text, cells activated by thrombin and SC showed enhanced activation relative to thrombin alone, as seen from the several criteria:

1) the brightness of DGs released during the 1st release event increased from 27.3 ± 2.4 × $10^4$ a.u. to 42.3 ± 3.6 × $10^4$ a.u. (Figure 5B);

2) time of the 1st release event decreased from 49.2 ± 5.2 to 15.8 ± 2.3 s (Figure 5C);

3) number of calcium spikes before the 1st release event decreased from 9.5 ± 0.9 to 4.5 ± 0.5 (Figure 5D);

4) frequency of calcium spikes before the 1st release event increased from 12.1 ± 0.5 to 18.1 ± 2.0 spikes/min (Supplementary Figure 8E).

Due to the complexity of platelet activation mechanisms, it is not immediately evident whether these specific changes can be consistently and quantitatively described within the proposed cooperative (bursting) mechanism. This mechanism posits that the release of the first DG(s) during a secretion event enhances the probability of subsequent DG release, establishing a positive feedback loop that drives the coordinated release of multiple DGs within a single event. If this feedback is mediated by the released content of the DG(s) themselves, then adding the SC preparation to thrombin-activated cells should amplify secretion triggering.

To test this prediction quantitatively, we incorporated SC into the bursting secretion model. Specifically, the observed secretion amplification in the presence of SC was modeled by introducing (1) an increased probability of DG release and (2) an elevated calcium spike frequency.

1) The probability of the 1st DG release (also called event triggering) in the presence of SC, $p_{1SC}^{i}$, was amplified by factor K:

$$p_1^i(SC) = K\, p_1^i \tag{14}$$

where $K$ is the parameter for positive feedback in our model, and $p_1^i$ is the probability of DG release in platelets activated by thrombin alone, as described in Section 1.3.4. Thus, in the presence of SC, the first DG is secreted with the same enhanced probability as all subsequent DGs:

$$p_1^i(SC) = p_2^i \tag{14}$$

2) Experimentally determined frequency of calcium spikes in the presence of SC was used to infer the probability of spike generation, $p_{Ca}(T, SC)$. This probability increased 1.4 times compared to the probability $p_{Ca}(T)$ with same concentration of thrombin in the absence of SC.

All other model assumptions, equations, and parameter values remained unchanged from Section 1.7.

Numerical simulations were conducted for 300 s for each of the 100 virtual platelets, and the resulting population secretion curves were plotted (Supplementary Figure 12A). In the model, the number of DGs released during the first secretion event increased (Supplementary Figure 12B), while the timing of the first release and the number of calcium spikes preceding it decreased (Supplementary Figure 12C,D). Notably, these theoretical results closely align with the population secretion curve and the characteristics of the first release event (Supplementary Figure 12). Thus, the bursting secretion model effectively reproduces experimental results with SC, providing strong support for the model framework.

### Part 2. Modeling of thrombus formation

#### 1.10 General approach

To investigate whether the character and kinetics of DG secretion plays a role in thrombus formation, we used the previously developed model of activator-driven thrombus formation in a microvessel[1,3,4]. This published model was combined with the model of platelet secretion described in Theoretical Approaches, Part 1 of Supplementary Materials. The model of platelet secretion described in Part 1 of this Supplementary text was incorporated with no modifications other than required by system geometry, as described in detail in Section 1.11.5. The general framework of the combined model is described in Section 1.11 and Supplementary Figure 13A, and values of parameters were determined as described in Section 1.11.

Briefly,

- Thrombus forms at the site mimicking a blood vessel injury. The injury site is a source of thrombin, which diffuses away from the site and is diluted by flow. The site is further assumed to be covered in a monolayer of platelets, which simplifies the model by bypassing the reactions that lead to these initial binding interactions.
- Platelets covered with integrins and GPIb receptors flow in the vessel and adhere to each other and to platelets already bound to the injury site in a thrombin-dependent manner.
- Hydrodynamic forces are imposed on the growing thrombus. As the thrombus increase in size, the flow through the vessel is affected.
- Platelets are activated by thrombin in a time and concentration-dependent manner, depending on their spatial location within the dynamic gradient of thrombin concentration.

Activated platelets:

1) establish integrin-dependent contacts
2) generate calcium spikes and release DGs. For simplicity, we model the release and transport of SC as a single molecular entity rather than a mixture of agonists. In the bursting secretion scenario, SC triggers positive feedback in secretion.
3) SC contributes to platelet activation by increasing the amount of available integrins and induces DG release in other platelets. These responses are modeled using our experimentally determined dependencies for ADP, which represents the major DG component.

As the modeling output, we tracked the changes in thrombus size in time, the coordinates of platelets within the thrombus, DG release events, and local concentrations of thrombin and SC. With this combined model we investigated how thrombus growth depends on the character and timing of DG secretion. Specifically, three scenarios were examined: 1) secretion with feedback (bursting secretion), 2) secretion without feedback (random secretion), and 3) no DG secretion.

#### 1.11 Model description

##### 1.11.1 Model geometry

Blood vessel segment is represented by a two-dimensional rectangular area with stiff boundaries (height $H$ = 35 μm, length $X$ = 200 μm). A depth (1 μm) is introduced for calculation of fluid dynamics and concentrations of activators. Platelets are modeled as rigid discs with radius $r$ as described in Theoretical Approaches, Part 1, Section 1.2. The injury site is modeled as a line $L$ = 15 μm long. This line has a monolayer of irreversibly attached platelets, which are used in this simplified model as a nucleation seed for the aggregation of platelets introduced with blood flow. This approach bypasses the need to mathematically represent tissue-dependent activation, which typically induces cell adhesion to wounded tissue. Here, it allows the model to focus on thrombus growth dynamics directly.

##### 1.11.2 Modeling blood hydrodynamics

We adopt the approach to model blood hydrodynamics from ref[1] without modification. In brief, blood plasma is modeled as a non-compressible Newtonian fluid described by Navier-Stokes and continuity equations. The flow in the vessel is from left to right and it has the wall shear rate $\dot{\gamma}$ = 1,000 $s^{-1}$. Explicit pressure and velocity fields are obtained for a part of the vessel that includes the injury site (computational domain). Attached platelets are impermeable for the flow. Vessel walls and platelet surfaces are treated as no-slip boundaries. The probability of platelet entering the computational domain depends only on the flow velocity and is chosen to provide realistic concentration of floating platelets. For detailed description of this framework and choice of parameter values, see ref[1].

##### 1.11.3. Modeling soluble platelet activators

Soluble activators are virtual particles, which are modeled as in ref.[1] with minor modifications. Briefly, these particles are dimensionless and massless, they do not interact with each other, and their motions are described by Langevin dynamics. In the computational domain, the particles are subjected to advection and diffusion with diffusion coefficients for thrombin and SC. The diffusion coefficient for SC is assumed to be the same as for ADP, see ref[1]. Thrombin particles are generated at the injury site, forming a constant flux $J_T$. To mimic volumetric inhibition of thrombin in plasma, thrombin particles are eliminated randomly with a probability $p_i$, as described in ref.[1]. The value of $J_T$ was reduced 10-fold compared to the flux in ref[1] to ensure that the thrombin-activated part of the modeled thrombus occupies not more than half of the total thrombus area, in correspondence with experiment[5].

At the computational step which led to DG release, the SC particles are generated at the location of the secreting cell *in situ*. The number of activator particles in the vicinity of the cell is used to determine the local concentration of SC, which is then used in the model to amend the platelet's activation, as described in Sections 1.11.4.2 and 1.11.5. The choice of parameters values is discussed in Section 1.12.

#### 1.11.4. Modeling platelet-platelet adhesion

We employed the same approach for modeling mechanics of single platelets and their interactions, as in Kaneva et al., ref[3]. In brief, the flow exerts force and torque on the free-flowing platelets in accordance with Stokes law. These forces pull platelets away from the injury site, and they are counteracted by platelet adhesion to each other and to the platelets immobilized at the injury site. The strength of platelet adhesion depends on their state of activation described with the parameter $\alpha$, which is determined by the local concentrations of activators. The total level of activation for each platelet at a given time is a sum of thrombin-induced and SC-induced activation: $\alpha = \alpha_T + \alpha_{SC}$. The activation level of a resting platelet (in the absence of any activators) is zero. Activation by thrombin leads to an irreversible increase of $\alpha_T$ as a function of current local thrombin concentration and total time of exposure to thrombin. Parameter $\alpha_T$ determines irreversible platelet-platelet binding, which is modeled via Morse potential; these bonds were modeled based on $\alpha_{IIb}\beta_3$-fibrinogen connections that link adjacent platelets in a real thrombus. Exposure to SC leads to reversible SC-induced activation $\alpha_{SC}$, which is calculated in our model as in ref[1]. The value of $\alpha_{SC}$ depends on the local SC concentration, and it defines reversible platelet binding, which is modeled with stochastically associating and disassociating springs. These weak interactions roughly correspond to the platelet-platelet interactions mediated by the GPIb receptor and the WF. All forces acting on each platelet in the computational domain are determined at every moment, and the equations of motions are solved to calculate platelet dynamics.

#### 1.11.5 Modeling DG secretion

*1.11.5.1. DG secretion in response to calcium spiking*

The model for DG secretion in isolated platelets (Theoretical approaches, Part 1 of Supplementary Materials) was incorporated without any modifications into the model of thrombus formation. In the resulting real-time model of the growing thrombus, local thrombin and SC concentrations change quickly owing to their production (or secretion), diffusion, and dilution by flow. Thus, equations (12) and (13) for single cell activation are used with the local concentrations of thrombin and SC. These concentrations are calculated at each time step for each platelet as in Section 1.11.3. Second, the model assumes that the effect of thrombin and SC on DG release is additive. Thus, if both activators are present simultaneously, the probabilities $p_1^i$ and $p_2^i$ for DG secretion are determined as a sum of probabilities to be released in response to thrombin and SC:

$$p_1^i = p_1^i(T) + p_1^i(SC) \quad (15)$$

$$p_2^i = Kp_1^i \quad (16)$$

Furthermore, when SC and thrombin act on platelets simultaneously, the frequency of calcium spikes was assumed to increase relative to the frequency at the same thrombin. Equation (5) for the probability of calcium spikes was modified accordingly:

$$p_{Ca}(T, SC) = \Delta t\ a\ \log(c\,T + d) \quad (17)$$

where $c$ is the enhancement factor from the SC, $d$ is the factor that determines response to SC in the absence of thrombin, and other parameters are as in Equation (5). Parameter values $c = 4.3 \cdot 10^9$ ml/U and $d = 117.9$ provide 18.1 ± 2.0 spikes/min in response to a mixture of 0.1 U/ml thrombin and SC, as in experiment (Supplementary Figure 8E).

Lastly, to avoid inappropriate triggering of secretion, we introduced the threshold concentration $C_{eff}$ for SC. If local SC concentration was below $C_{eff}$, DG secretion proceeded as described in Theoretical Approaches, Part 1 of Supplementary Materials, causing autocrine secretion amplification which involved only the secreting cell. If the local SC concentration exceeded the $C_{eff}$ threshold, DG secretion proceeded as described in Section 1.9 for platelets activated by thrombin and SC. Thus, at high SC concentrations, the amplification mechanism included the paracrine effect, in which cells that have not yet released any DGs have increased their release probability owing to the close proximity to actively secreting cells. The value of $C_{eff}$ = 5 µM was chosen based on the concentration of ADP measured immediately after DG secretion by activated platelets[6,7].

*1.11.5.2. Modeling different scenarios of secretion in the growing thrombus*

To investigate how DG secretion affects thrombus dynamics, we used the combined model to compare three scenarios of thrombus formation: 1) bursting secretion, 2) random secretion, and 3) without DG secretion.

*1. Bursting secretion*

This scenario was analyzed using approaches and parameter values described in Section 1.11.5.1.

*2. Random secretion*

In this scenario, to simulate the absence of both autocrine and paracrine effects of SC, positive secretion feedback was eliminated by setting $p_1^i(SC)$ to 0 and adjusting the secretion algorithm described in Section 1.5. Specifically, in step 3a, the DG with the largest $p_1^i$ was released with this probability. Values of parameters of secretion were determined as described in Section 1.8. This secretion scenario features enhanced secretion sensitivity to thrombin, see Section 1.8

*3. Without DG secretion*

In this scenario, the number of DGs $N_{DG}$ = 0 in all platelets, so the platelets forming the thrombus do not secret any granules. Thrombus formation in this case is determined exclusively by thrombin gradient.

#### 1.12. Numerical simulations of thrombus formation

Program code was written in C++ and calculations were performed using Lomonosov-2 supercomputer resources at Lomonosov Moscow State University, Russia. CFD calculations are performed using simpleFoam solver provided by open-source OpenFOAM software (OpenFOAM Foundation, UK). Equations of motions were parallelized using OpenMP and solved using a modified Verlet algorithm. Langevin equations for soluble platelet activators were solved on CUDA-enabled GPUs. Coordinates of soluble particles were used to create a coarse-grid (1x1x1 µm) concentration map, which was used to determine local activator concentrations. The minimally resolved concentration for SC was 150 nM, and 0.042 U/ml for thrombin.

#### 1.13. Processing of Model-Generated Data for Visualization and Quantification

We conducted 7 stochastic simulations, each spanning 120 seconds of model time, for each secretion scenario. The simulated platelet coordinates were used to generate computational movies 5 and 7. In these visualizations, platelets are color-coded according to their activation parameter, $\alpha_T$. When $\alpha_T$ exceeds 15%, indicating relatively strong activation, the platelet is classified as part of the thrombus core (Supplementary Figure 13B). This empirically derived threshold effectively differentiates the thrombus structure into core and shell regions, as validated by visual comparison with experimental results in ref[8]. Computational movies 6 and 8 were generated using computed local SC concentration.

To quantify thrombus dynamics, we measured the relative thrombus height over time by dividing the distance from the vessel wall to the most distal platelet in the thrombus by the vessel width. The relative core height was calculated similarly, using the distance to the most distal platelet within the core. The relative shell height was then determined as the difference between the relative thrombus height and the relative core height.

### 2. Supplementary Notes

#### Supplementary Note 1. Quantification of DG count in human platelets.

Dense granules (DGs) exhibit variable sizes and morphologies, making it challenging to determine their exact number with certainty. Previous studies have used mepacrine staining of platelet DGs, assuming each mepacrine-stained spot represents an individual granule[9–14], leading to an estimated average of 5–6 granules per platelet[9,12–14]. Other methods for measuring and quantifying DGs have estimated between 3 and 8 DGs per platelet[15–22]. Variability in DG estimates is not unique to human platelets. A study using mepacrine labeling in rabbit platelets reported 16 DGs per platelet[13], while single-cell voltammetry, which detects the release of individual granules, estimated 20–25 DGs[23]. Additionally, mouse platelets have been reported to contain approximately 6 DGs[13,24], whereas a recent study using FIB/scanning electron microscopy and 3D reconstruction found about 18 granules per platelet[25].

In our study, we observed that mepacrine-stained dots in human platelets varied in brightness, likely because some granules were closely packed and could not be resolved individually by epifluorescence microscopy. To quantify DG numbers in platelets, we employed two approaches. Both utilized direct counting of mepacrine-positive spots, and images were collected with epifluorescence or structured illumination microscopy (SIM). These approaches resulted in on average 5 DGs with regular epifluorescence or 10 DGs with SIM microscopy. The latter estimate using high resolution method is consistent with the upper range of the reported number of DGs in human cells[13] (Figure 1C,D). The slightly higher DG count observed in our study compared to previous reports[15–22] may be explained by our use of live platelets and minimal handling to reduce artificial activation. Previous studies may have underestimated DG numbers due to the high sensitivity of platelets to handling during preparation. Prior studies often used washed and, in some cases, permeabilized platelets (e.g., for transmission electron microscopy, focused ion beam-scanning electron microscopy, or super-resolution microscopy)[18,22], or dried platelet suspensions (e.g., whole-mount transmission electron microscopy)[15,17]. Additionally, defocused or blurred images of intracellular granules may contribute to underestimation due to the diversity of dense structures[26].

Finally, we used the total cell brightness and the intensity of single granules imaged with epifluorescence microscopy to estimate the upper limit of the number of DGs, as determined with mepacrine brightness. We reasoned that the individual dots with a spherical shape and minimal brightness likely corresponded to single granules. The following criteria for selecting the smallest dots were used: 1) each dot could be fit within a circular region 0.54 μm in diameter (corresponding to 4 pixels); 2) the dots were at some distance from each other, so there was no overlap in fluorescence; 3) upon activation, the fluorescence of each dot dropped to the background level during one frame of 0.5 s, as expected for a single granule secretion (Supplementary Figure 4F,G). The median value of the intensity distribution of the smallest mepacrine dots corresponded to the integrated intensity of a single minimally-sized DG (Supplementary Figure 4H). The number of DGs per cell was then determined from the ratio of this cell intensity and intensity of the smallest fluorescent dots determined for cell population ($n$ = 80 – 120 cells) imaged under identical conditions. With this method, we estimate that the maximum average number of DGs is ~ 19 per cell (Supplementary Figure 4I). This estimate exceeds the average number of DGs by only a factor of 2, indicating that size heterogeneity is not very significant. Importantly, the major conclusions of our study regarding secretion in discrete, temporally correlated sets are based on direct visual observations of individually resolved DGs and are therefore independent of the exact DG count.

**Supplementary Note 2. Spatial DG distribution does not correlate with temporal secretion coordination.**

Cooperative DG secretion may be determined, at least in part, by DG spatial organization of secretion. Spatial confinement of exocytosis events has been reported is many cell types[27–31], and DG secretion has also been suggested to exhibit spatial localization[32], potentially explaining temporally-coordinated secretion. Alternatively, several DGs may secrete simultaneously owing to their spatial proximity, although unlike α-granules the DGs do not appear to undergo compound fusion[22]. Testing these hypotheses in platelets, which are among the smallest cells in the human body[33], is difficult because many DGs are localized together and individual granules cannot be resolved using light microscopy. Our measurement of distance between co-released granules (see Results Chapter 1 and Figure 1G) showed that co-secreting granules were found all over the cell, ruling out presence of preferred sites of exocytosis. Thus, close proximity between DGs or their specific localization is not required for the synchronous release.

Prior studies found that a large fraction of DGs in resting platelets is located close to the plasma membrane[34,35]. Such granules could represent a readily releasable pool, as seen for secreting synaptic vesicles in neuronal cells[36]. Thus, we examined if the DG release depends on their proximity to plasma membrane. We surmised that if DGs located within this membrane-proximal area are more prone to secretion, most of them will be released during the very first secretion event. However, the percent of the released granules recorded via TIRF was only ~ 38%, which is similar to the percentage of granules released in platelets imaged by epifluorescence (see Results Chapter 1 and Supplementary Figure 3C). We concluded that although the close proximity of granules to the membrane may play a role, it is not the main factor leading to synchronized granule secretion.

**Supplementary Note 3. Neither calcium spike amplitude nor frequency determine DG release timing.**

Because not all calcium spikes were followed by the DGs release, we wondered if the spikes that preceded secretion had any special features that may control timing of secretion or number of secreted DGs.

First, we tested that a calcium spike of a specific amplitude induces release of DGs. Our data indicates that the average amplitude of preceding calcium spikes (1.17 ± 0.71 a.u.) was 22% larger than that in other spikes (0.96 ± 0.83 a.u.). However, their significant variability and overlapping of amplitude distributions indicated a lack of direct correlation between the occurrence of the release events and the preceding spike amplitude (Supplementary Figure 7A,B).

Second, the average interval between two preceding spikes and the other spikes was also similar (Supplementary Figure 7C). These results are consistent with our finding that the total interval between the consecutive release events is distributed exponentially (Figure 2D). Thus, calcium-induced event triggering is stochastic in nature.

**Supplementary Note 4. Estimating ATP, ADP, and serotonin concentrations in secreted content.**

We estimated concentration of ADP secreted in our experiments by human platelets using published data. Previous work by Holmsen et al.[37] demonstrated that $10^{11}$ platelets secrete $3 \cdot 10^{6}$ moles of ADP. We assume that platelet concentration in human blood is $2.5 \cdot 10^{5}$ platelets/μl, which is in the middle of the reported physiological range: $1.5 \cdot 10^{5}$ – $4 \cdot 10^{5}$ platelets/μl (ref.[38]). During preparation of platelet-rich plasma (PRP), platelet concentration increases by at least 1.5 times (ref.[38]), resulting in $3.75 \cdot 10^{5}$ platelets/μl. Furthermore, in our protocol, 1.4 ml PRP was centrifuged to pellet the cells, which were then resuspended in 800 μl. Assuming no cell loss, this procedure produces a suspension with $6.5 \cdot 10^{5}$ platelets/μl, containing in total $5.2 \cdot 10^{8}$ cells. Thus, platelets activated in our experiments generate $15.6 \cdot 10^{-9}$ moles of ADP or ~ 20 μM. To estimate concentration of ATP, we took into account the previously established ratio of ATP/ADP concentrations in platelets: ~0.7 (ref.[37]). The quantity of added ATP is 13.3 μM. Furthermore, the serotonin concentration (ref.[39]) is analogous to the ADP concentration (ref.[37]) in DGs. Consequently, the added serotonin concentration is 20 μM.

### 3. Supplementary Tables

**Table 1.** Parameters and variables for the model of DG secretion in platelets.

| Parameter symbol | Parameter name | Value | Units | Source |
|---|---|---|---|---|
| **General model parameters** | | | | |
| $r_o$ | parameter in Equation (3) | 1 | μm | section 1.2 |
| $r_\sigma$ | parameter in Equation (3) | 0.22 | μm | |
| $N_0$ | parameter in Equation (4) | 19 | unitless | section 1.2 |
| **Calcium spikes** | | | | |
| $p_{Ca}(T)$ | parameter in Equation (5) | variable | unitless | section 1.3.1 |
| $a$ | parameter in Equation (5) | 1.38 | $min^{-1}$ | section 1.3.1 |
| $b$ | parameter in Equation (5) | 9.79 | unitless | |
| $p_{Ca}(ADP)$ | parameter in Equation (6) | $2.2 \cdot 10^{-4}$ | unitless | section 1.3.1 |
| $p_{Ca}(T,SC)$ | parameter in Equation (17) | variable | unitless | section 1.9 |
| c | parameter in Equation (17) | $4.3 \cdot 10^{9}$ | unitless | section 1.9 |
| d | parameter in Equation (17) | 117.9 | $min^{-1}$ | |
| **DG release** | | | | |
| $t_{Ca}$ | window of opportunity | 3 | s | section 1.3.3 |
| $t_K$ | release event time | 3 | s | section 1.3.4 |
| $n_{lag}(T)$ | delay parameter | 7 | unitless | calibrated, section 1.6.1 |
| $K$ | feedback coefficient | 7.76 | unitless | |
| $n_{lag}(ADP)$ | delay parameter | 0 | unitless | section 1.6.2 |
| $\sigma(T)$ | parameter in Equation (12) | variable | unitless | variable, section 1.6.1 |
| $a_1$ | parameter in Equation (12) | 0.85 | unitless | calibrated, section 1.6.1 |
| $b_1$ | parameter in Equation (12) | 0.26 | ml $U^{-1}$ | |
| $\sigma(ADP)$ | parameter in Equation (13) | variable | unitless | variable, section 1.6.2 |
| $a_2$ | parameter in Equation (13) | 1 | unitless | calibrated, section 1.6.2 |
| $b_2$ | parameter in Equation (13) | $2.4 \cdot 10^{-5}$ | $\mu M^{-1}$ | |

### 4. Supplementary Figures and Figure Legends

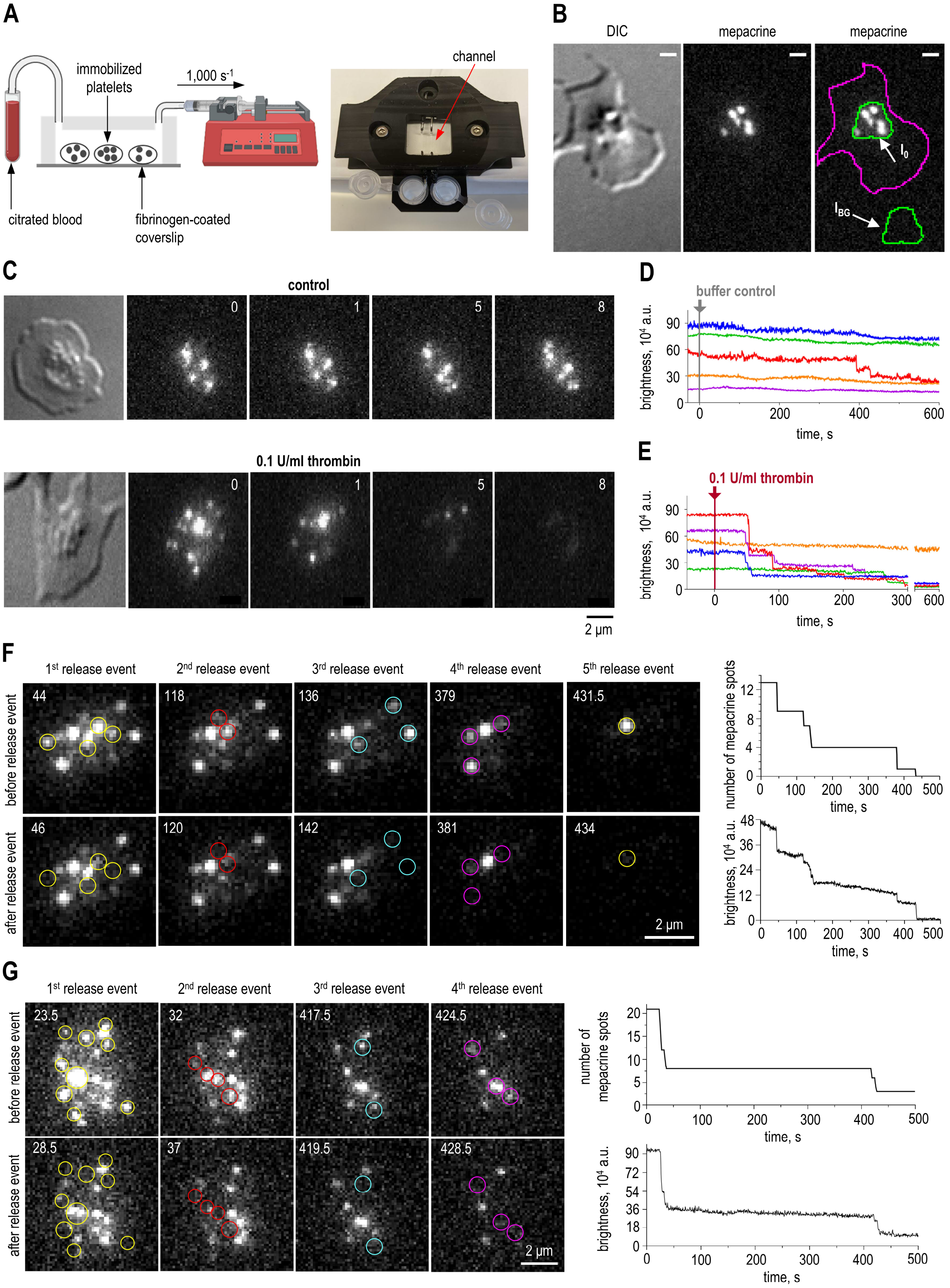


**Supplementary Figure 1. Quantification of DG number and secretion kinetics in single platelets.** (A) Schematics of the flow chamber with a pump-driven blood perfusion to immobilize platelets for subsequent activation (left). Photograph of microfluidic flow chamber in 3D printed holder (right). (B) Example of a fibrinogen-adhered platelet imaged via DIC and mepacrine fluorescence. The purple line shows cell border determined from the DIC image. The green line shows the area containing all mepacrine stained dots inside the platelet, which was used to measure integrated intensity ($I_0$). The same area was then shifted to an area adjacent to cell to collect the integrated background intensity ($I_{BG}$). Visually counted number of mepacrine spots is 6. Scale bar is 1 μm. (C) Still images of a typical control platelet and a thrombin-activated platelet. Numbers correspond to the time in minutes after the addition of a buffer (control) or thrombin. The disappearance of small fluorescent dots and decreased brightness of the large dots indicate DG release. (D) Changes in the total cell brightness in untreated platelets. Each curve corresponds to a single platelet imaged for 10 min. (E) Analogous quantification as in panel D but for thrombin-activated cells. (F)-(G) Still images and kinetics of secretion of a platelet activated by 0.1 U/ml thrombin at 0 s. The epifluorescence images show mepacrine-labeled DGs. The signal was not saturated and identical linear brightness adjustments were applied to all images in the panel. The numbers show time in seconds after thrombin addition. Each pair of images, from left to right, show start and end of a release event, with circles highlighting mepacrine spots released during the same release event. The graphs on the right quantify visually counted number of mepacrine spots (top) and total cell brightness (bottom) over time for the same cell.

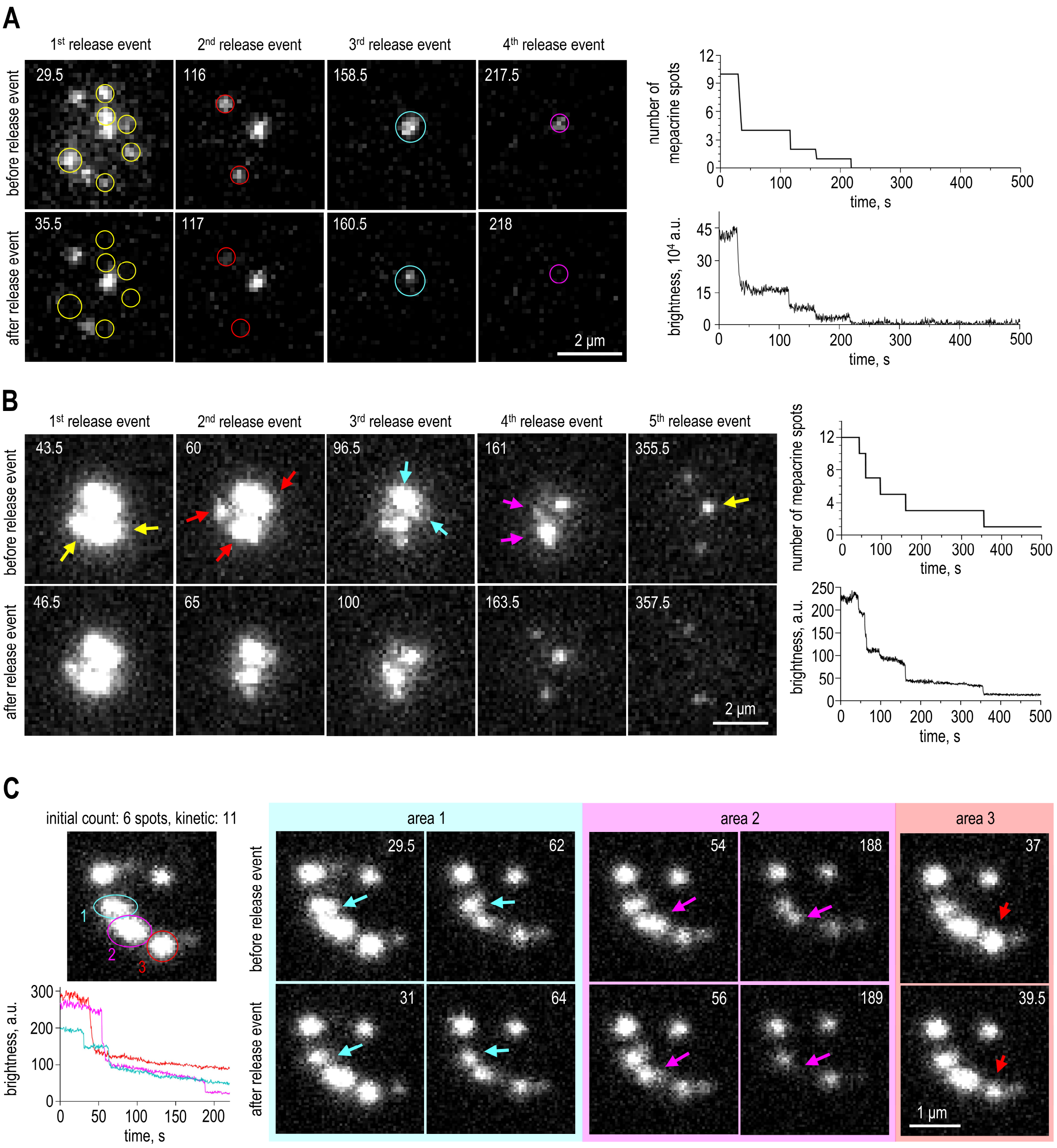


**Supplementary Figure 2. Examples of secretion kinetics in single platelets.** (A) Still images and kinetics of secretion of a platelet activated by 0.1 U/ml thrombin at 0 s. The epifluorescence images show mepacrine-labeled DGs. The signal was not saturated and identical linear brightness adjustments were applied to all images in the panel. The numbers show time in seconds after thrombin addition. Each pair of images, from left to right, show start and end of a release event, with circles highlighting mepacrine spots released during the same release event. The graphs on the right quantify visually counted number of mepacrine spots (top) and total cell brightness (bottom) over time for the same cell. (B) Still images and kinetics of secretion of a platelet activated by 0.1 U/ml thrombin at 0 s. See legend to panel A for more details. The images demonstrate DGs that are not optically resolvable. However, the secretion kinetics suggest the presence of multiple granules within these spots. This is inferred from observing changes in the shape and brightness of mepacrine spots, alongside the kinetics of total cell fluorescence over time, which shows a partial decrease in the brightness of mepacrine-labeled spots. Arrows in the images indicate the specific locations where brightness decreases. (C) SIM microscope measurement of a platelet activated by 0.1 U/ml thrombin. The first image on the left shows the platelet image prior to activation. Highlighted in three distinct colors, designated areas within the cell are monitored for changes in fluorescence intensity, representing regions where total brightness is quantitatively measured over time, as shown in the graph at the bottom. The numbers show time in seconds after thrombin addition at 0 s. DG releases, observed by fluorescent drops, are presented as still images before and after release for each area. Arrows indicate the specific locations where brightness decreases. Note that the brightness units are different in this panel as experiment was conducted using a SIM microscope under different settings and photobleaching was not corrected, see Materials and Methods for details.

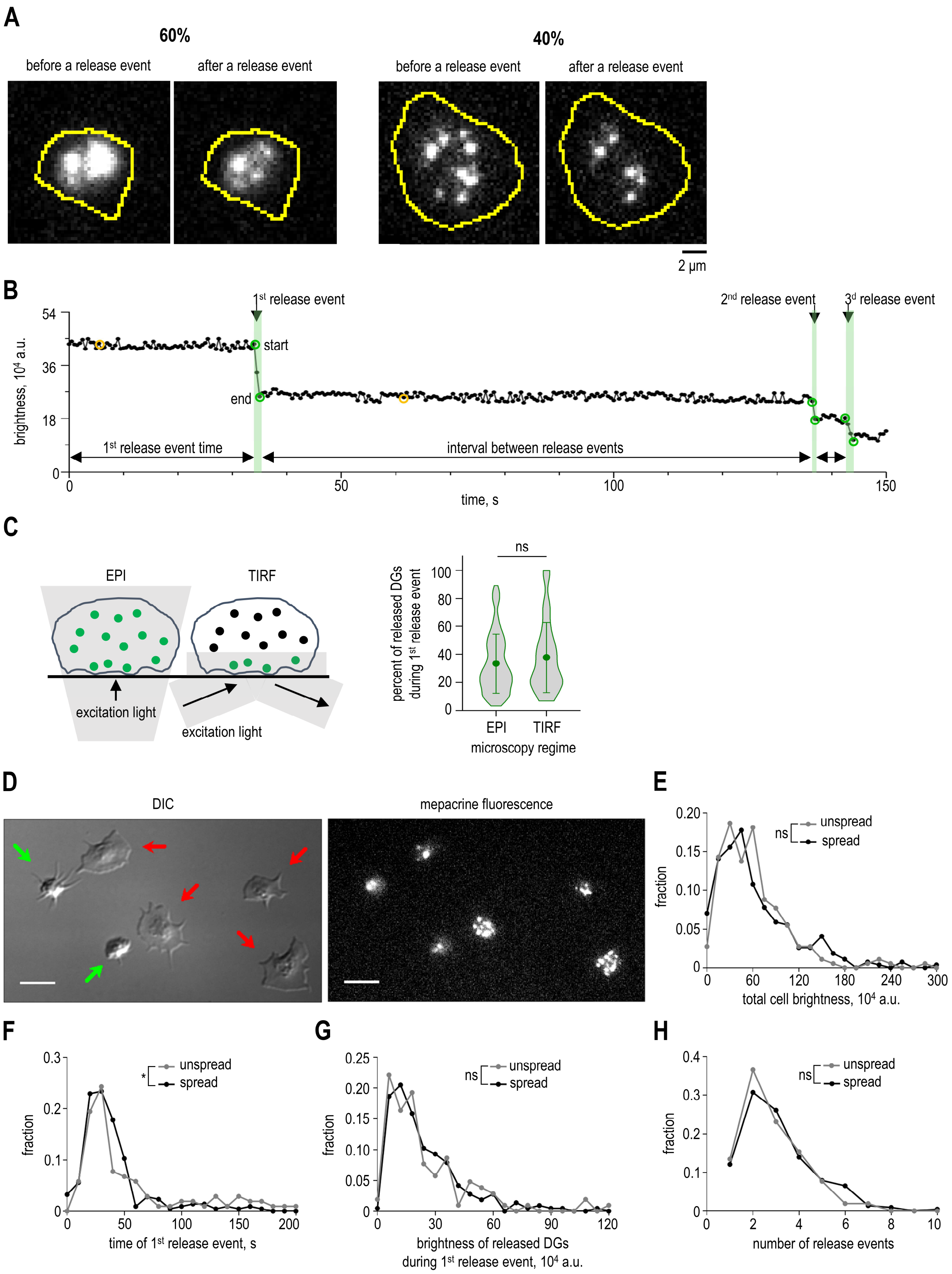


**Supplementary Figure 3. Analysis of secretion under different illumination regimes and in platelets of different shape.** (A) Example images of platelets immediately prior to and after the secretion. The numbers indicate occurrences in the total platelet population in experiments with 0.1 U/ml thrombin. As in the cell shown in the right images, granules were scattered far enough to enable pairwise distance analysis between them. Yellow lines show cell borders determined from DIC images. (B) Example of analysis of an integrated intensity curve. Release events, indicated by black arrows, were identified by an abrupt drop in fluorescence. Green circles show the start and end of a release event, which were determined visually using the corresponding imaging series.

The time between activator addition (0 s) and the start of the 1st release event is further called "1st release event time". Then, time intervals were measured analogously between other release events. (C) The schematic illustrates two illumination modes to visualize dense granules, epifluorescence (EPI) and total internal reflection fluorescence (TIRF), which are characterized by different depth of light penetration. With TIRF-based illumination, granules located within ~ 200 nm of the coverslip-attached plasma membrane are visualized (shown as green dots). Epifluorescence visualizes granules located farther away from the cell membrane. The graph shows the percent of released DGs in cells activated by 0.1 U/ml thrombin based on $N = 3$ and $n = 42$ for TIRF, $N = 20$ and $n = 230$ for EPI. Mann-Whitney U test: ns – not significant. (D) Example field of the platelets imaged via DIC and mepacrine fluorescence on a fibrinogen-coated coverslip. Platelets with filopodia are indicated by the green arrows, platelets with lamellipodia are indicated by the red arrows. Before activation, platelets were categorized based on their shape by visual observations. Platelets with round-like shape, with or without filopodia, were categorized as "unspread", while platelets with lamellipodia were categorized as "spread". Scale bar is 5 μm. (E) Distributions of the total cell brightness in spread and unspread platelets. Here and in panels F-H: platelets were activated by 0.1 U/ml thrombin, $N = 36$, $n = 452$, see Data Source File for details. Kolmogorov-Smirnov test: ns – not significant. (F) Distribution of time of the 1st release event of spread and unspread platelets. Kolmogorov-Smirnov test: * is p-value $< 0.05$. (G) Distribution of the brightness of released DGs during 1st release event of spread and unspread platelets. Kolmogorov-Smirnov test: ns – not significant. (H) Distribution of the number of release events of spread and unspread platelets. Kolmogorov-Smirnov test: ns – not significant.

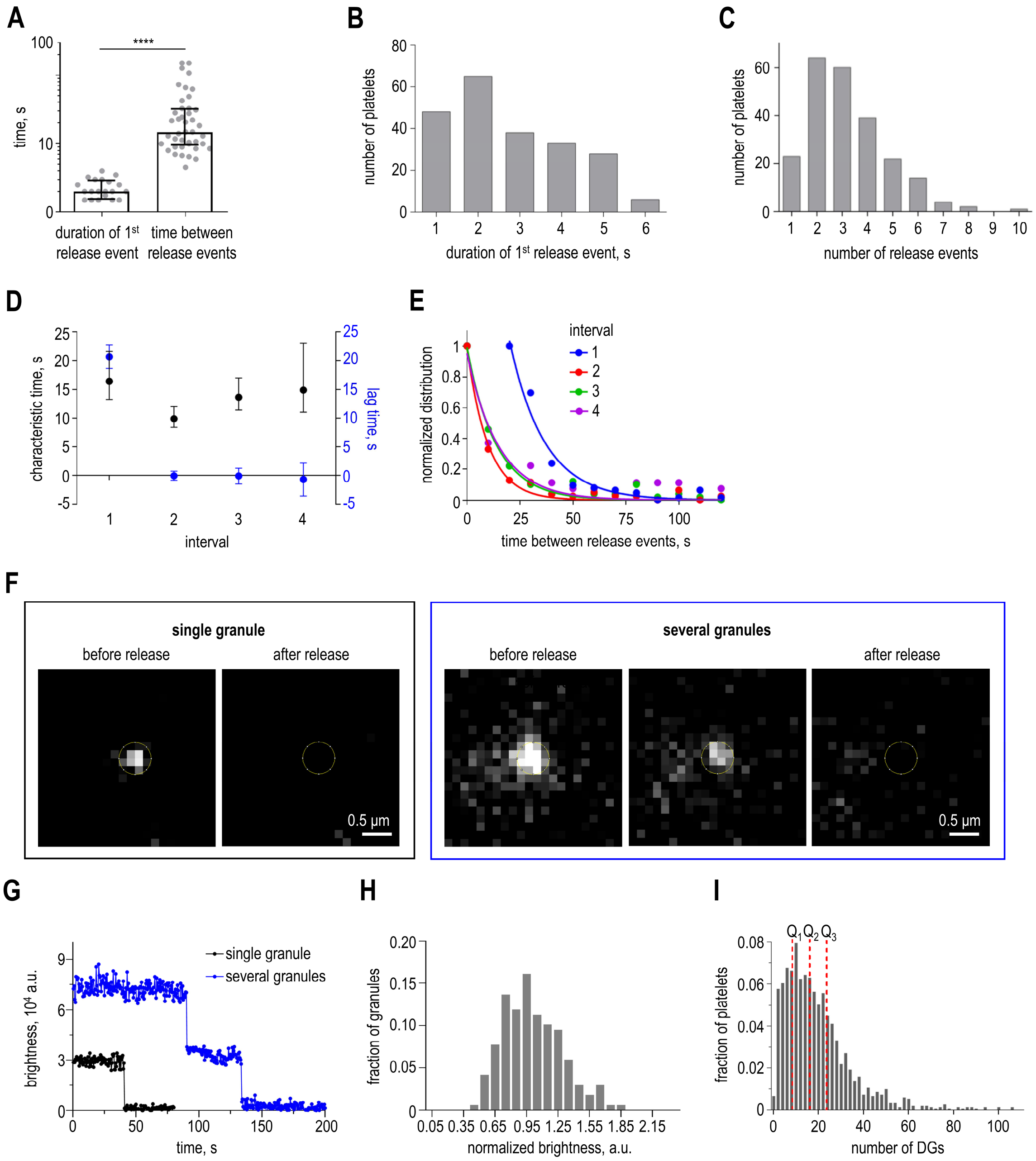


**Supplementary Figure 4. Quantification of DG secretion kinetics in response to thrombin activation and estimation of DG number.** (A) Comparison of the duration of a release event with the time of the intervals between release events in cells activated by 0.1 U/ml thrombin. Here and in other graphs, dots represent the median results from *N* independent experiments, bars are their means ± SEM, and *n* is the total number of platelets. The left column presents the duration of the 1$^{st}$ granule release event of *N* = 20 and *n* = 230. The right column presents the time of the intervals # 2 – 4 of *N* = 14 and *n* = 171, 159, 129, and 89 platelets for consecutive events, see Data Source File for more details. Mann-Whitney U test: **** is p < 0.0001. (B) Distribution of the duration of the 1$^{st}$ release event. The average duration is 2.5 ± 0.1 s. Platelets were activated by 0.1 U/ml thrombin in *N* = 20, *n* = 230. Bin size is 1 s. (C) Distribution of the number of release events. Bin size is 1 release event. See legend to panel B for details. (D) Exponential fitting of distributions of time intervals. Distributions are shown for the time between the activator addition and the 1$^{st}$ release event (interval 1), between the 1$^{st}$ and 2$^{nd}$ (interval 2), 2$^{nd}$ and 3$^{d}$ (interval 3) and 3$^{d}$ and 4$^{th}$

(interval 4) release events. The dots represent the mean of $N$ = 14, $n$ = 171, 159, 129 and 89 platelets for consecutive events. The line represents the exponential fitting (see Materials and Methods and Data Source File for details). (E) Parameters of exponential fitting, plotted for different time intervals as described in panel D. (F) Example of a single granule (left images) and several granules (right images) imaged via mepacrine fluorescence. (G) Granule brightness over time from panel F. When a single granule (black curve) or several granules (blue curve) are released, brightness drops to zero. Note that when one mepacrine spot comprises several granules (blue), the release occurs in several steps. (H) Distribution of normalized single granule brightness. The data were acquired using a Zeiss microscope with a 385 nm diode (number of single granules is 117) and a Nikon microscope with a 488 nm laser (number of single granules is 79), see Materials and Methods for details. Distributions were normalized by their peak value and then averaged. (I) Distribution of the number of the minimally—sized DGs in platelets. The red dashed lines indicate the median ($Q_2$) = 16.14, $Q_1$ = 8.57, $Q_3$ = 25.5, based on $N$ = 99 independent experiments and $n$ = 1,508 total number of platelets. Bin size is 2 granules.

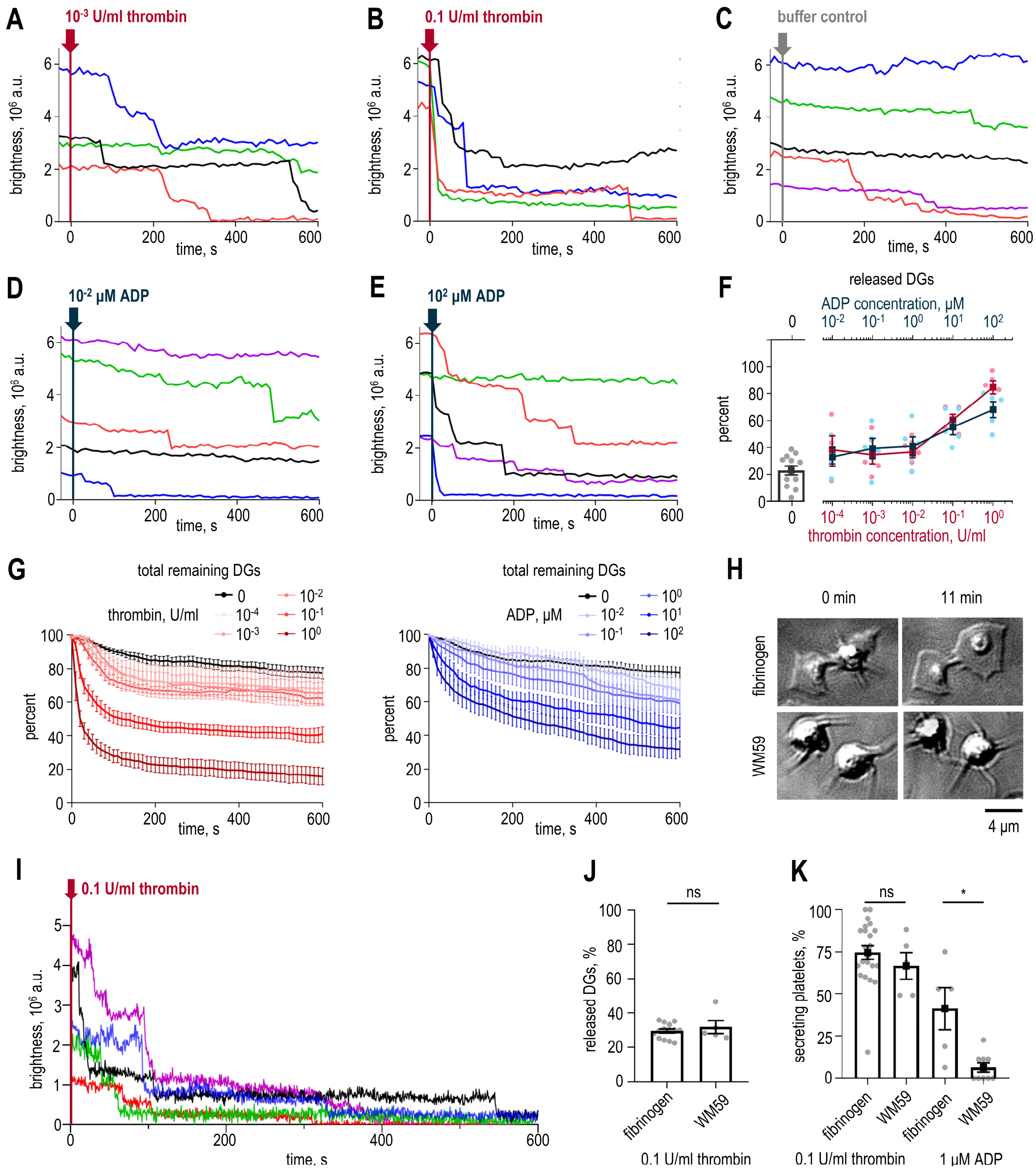


**Supplementary Figure 5. DG secretion kinetics upon thrombin or ADP activation in single platelets adhered on fibrinogen or WM59 antibodies.** (A)-(E) Changes in the brightness of DGs in platelets. Each secretion curve corresponds to one cell. (F) Percent of released DGs within 600 s after addition of the indicated activator: thrombin (bottom axis, red dots) or ADP (top axis, blue dots) concentrations are plotted on the semi-log scales. Here and in panel G: $N = 4 – 5$ and $n = 58 – 94$ for each ADP concentration, $N = 5 – 6$ and $n = 81 – 124$ for each thrombin concentration, $N = 11$ and $n = 164$ for control platelets with no ADP or thrombin. See Data Source File for results of Mann-Whitney U test. (G) Population secretion curves of the percent of remaining granules. The graphs show curves for the percent of remaining granules in control platelets (black curve, both graphs) compared to platelets activated by indicated concentrations of ADP (blue curves, right graph) or thrombin (red curves, left graph). (H) DIC images of platelets adhered to coverslips coated with fibrinogen (top row) or WM59 antibodies (bottom row), imaged at 0 and 11 minutes in the absence of activation. (I) Changes in total mepacrine brightness in platelets adhered to

WM59 antibodies and activated by thrombin at 0 min. Each secretion curve corresponds to one cell. (J) Percent of DGs released during the 1st release event. Circles represent the mean value from each individual experiment; bars indicate the mean ± SEM across experiments. Data for fibrinogen and thrombin platelets are the same as in Figure 2B. Number of independent experiments $N = 5$ and total number of analyzed cells $n = 43$ for WM59 and thrombin. Mann–Whitney U test: * $p < 0.05$; ns, not significant. (K) Percent of platelets that secrete DGs within 1 min of activation. Fibrinogen and thrombin ($N = 22$, $n = 241$), WM59 and thrombin ($N = 5$, $n = 43$), fibrinogen and ADP ($N = 5$, $n = 66$), WM59 and ADP ($N = 9$, $n = 30$).

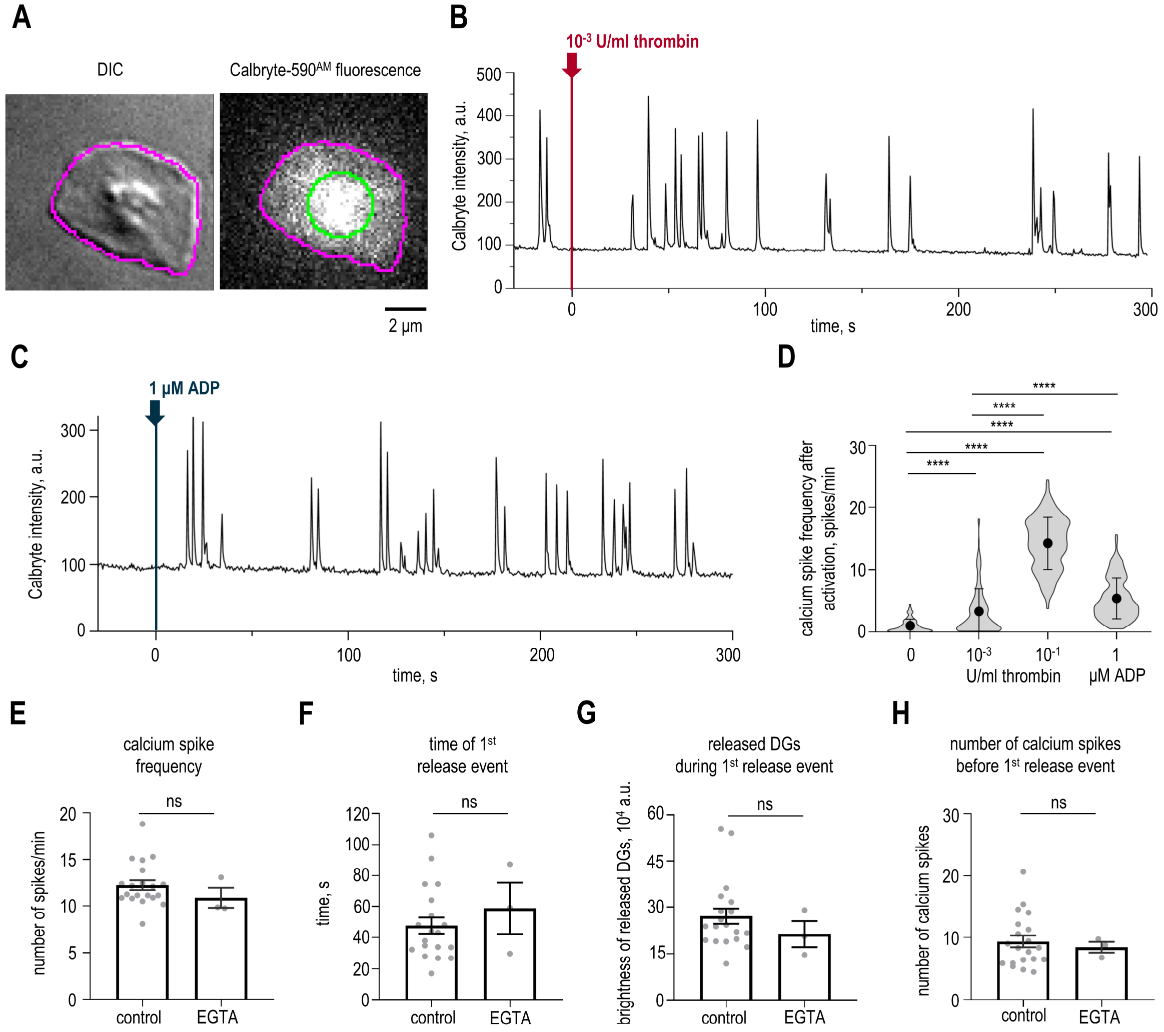


**Supplementary Figure 6. Quantification of calcium kinetics and the influence of extracellular calcium on DG secretion.** (A) Example of a representative platelet imaged via DIC and Calbryte-590$^{AM}$ fluorescence. The purple line shows cell border determined from DIC image. The green line represents the selected region within the platelet used to measure calcium intensity over time. (B) Example of Calbryte intensity curve of a single platelet activated by $10^{-3}$ U/ml thrombin. (C) Example of Calbryte intensity curve of a single platelet activated by 1 µM ADP. (D) Average frequency of calcium spikes. In the graph, dots with whiskers represent mean ± SD and contours show data distribution for *n* platelets in *N* individual experiments. Calcium spike frequency in control platelets was determined from measurement during 5 min before the addition of the activator ($N$ = 11, $n$ = 89). Calcium spike frequency in activated platelets was determined from measurement from activator addition until the end of the fluorescent measurement. Platelets were activated by different thrombin concentrations ($N$ = 17, $n$ = 166 and $N$ = 20, $n$ = 241 for $10^{-3}$ and 0.1 U/ml thrombin, respectively) and by 1 µM ADP ($N$ = 9, $n$ = 125). Note that data for both secreting and non-secreting platelets are included in the graph. Mann-Whitney U test: **** is p-value < 0.0001. (E) Calcium spike frequency before the 1$^{st}$ release event. Here and in panels F-H: dots represent the mean results from *N* independent experiments, bars are their means ± SEM. In the “control” experiments, platelets were activated by 0.1 U/ml thrombin in the presence of 2 mM $CaCl_2$ ($N$ = 20, $n$ = 230), while in the “EGTA” experiments platelets were activated by 0.1 U/ml thrombin in the presence of 5 mM EGTA ($N$ = 3, $n$ = 46). Mann-Whitney U test: ns – not significant. (F) Time of the 1st release event. See legend to panel E for more details. (G) Brightness of released DGs during the 1st release event. See legend to panel E for more details. (H) Number of calcium spikes before 1st release event. See legend to panel E for more details.

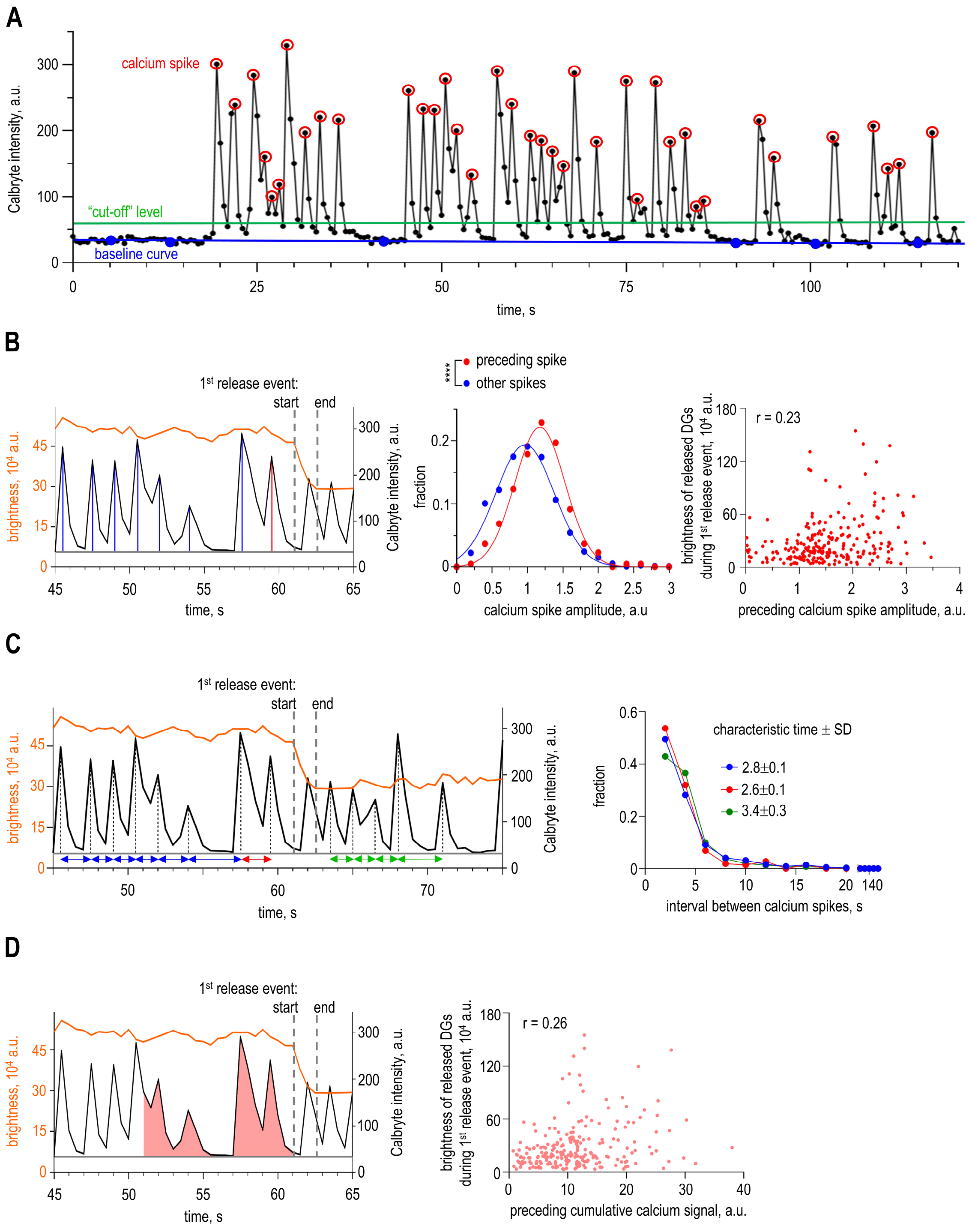


**Supplementary Figure 7. Quantitative analysis of calcium oscillations in relation to DG secretion.** (A) Representative trace showing whole-cell Calbryte-590$^{AM}$ fluorescent signal (indicative of $Ca^{2+}$ concentration; black curve) in a single platelet activated by 0.1 U/ml thrombin. Baseline curve (blue), "cut-off" level (green) and peaks of calcium spikes (red circles) are determined as described in Materials and Methods. (B) Example of calculation of calcium spike amplitudes (left panel), distributions of calcium spike amplitudes (middle panel) and correlation between brightness of released DGs and preceding calcium spike amplitude (right panel). Left panel shows an example of Calbryte intensity curve (black, right y axis) and brightness of DGs (orange curve, left y axis) of a single platelet activated by 0.1 U/ml thrombin. Calcium spikes amplitudes before the 1st release event (shown by blue lines and called "other spikes") and amplitude of preceding calcium spike (shown by red line and called “preceding spike”) were calculated as calcium spike’s maximum minus value with baseline curve value (grey curve) subtracted. Middle panel shows distributions of amplitudes of calcium spikes that were determined as shown in left panel. Since amplitudes of calcium spikes varied among different platelets, we normalized the amplitude of each calcium spike by dividing it by the average value of all spike amplitudes before the 1st release event. See Materials and Methods for details. Red dots show the distribution of relative amplitudes of preceding calcium spikes and the red line shows its Gaussian fitting. Blue dots show the distribution of relative amplitudes of all spikes that occurred from activator addition until the 1st release event. The blue line shows its Gaussian fitting. Here and in right panel, data are presented for platelets activated by 0.1 U/ml thrombin, $N = 20$, $n = 230$. Kolmogorov-Smirnov test: **** is $p < 0.0001$. Right panel shows correlation between the relative amplitudes of preceding calcium spikes and the brightness of released DGs during the 1st release event. Each dot represents data for a single activated platelet. The Pearson correlation coefficient (r) is plotted on the graph. See Data Source File for details. (C) Left panel shows an example of Calbryte intensity curve (black, right y axis) and brightness of released DGs (orange curve, left y axis) of a single platelet activated by 0.1 U/ml thrombin. Dual arrows below the baseline curve (grey) indicate intervals between the calcium spikes. Blue arrows show at the intervals between calcium spikes after activator addition and until 1st release event with the exception of the interval shown by red arrow, which is an interval between calcium spike preceding the 1st release event and calcium spike occurred before it. Green arrows show intervals between calcium spikes that occurred after the 1st release event ended. Right panel shows distributions of intervals between calcium spikes. Color code is the same as in left panel. The distributions were plotted for platelets activated by 0.1 U/ml thrombin, $N = 20$, $n = 230$, and fitted with a single decay exponent, see Data Source File for details. (D) Left panel shows an example of Calbryte intensity curve (black, right y axis) and number of DGs (orange curve, left y axis) of a single platelet activated by 0.1 U/ml thrombin. An example of the cumulative calcium signal for a 10 s window preceding the 1st release event is shown by red area under curve. Cumulative calcium signal was normalized dividing it by the average value of all spike amplitudes before the 1st release event for each platelet. See Materials and Methods for details. Right panel shows correlation between the relative preceding cumulative calcium signal for a 10 s window and the brightness of released DGs during the 1st release event. Each dot represents data for a single platelet activated by 0.1 U/ml, $N = 20$, $n = 230$. The Pearson correlation coefficient (r) is plotted on the graph. See Data Source File for details.

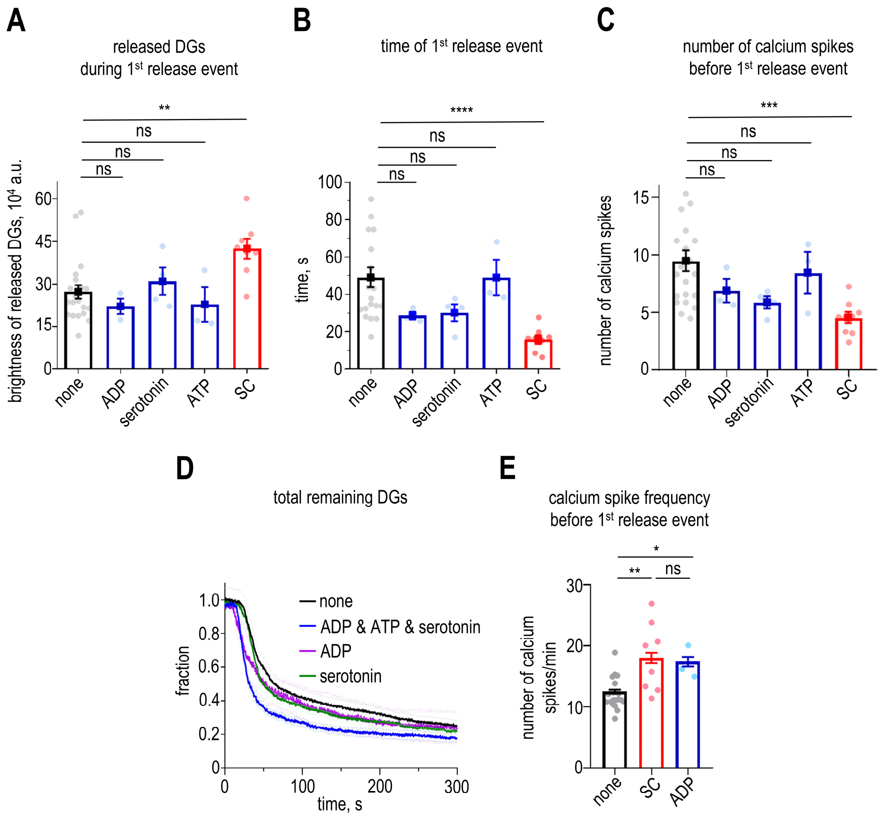


**Supplementary Figure 8. Experimental results of secretion in response to SC or components of DGs.** (A) Brightness of released DGs during the 1st release event in platelets activated solely by 0.1 U/ml thrombin or in combination with ADP, serotonin or ATP. Here and in panels B,C,E: the dots show the mean of individual experiments, while bars show their mean ± SEM. Platelets were activated using 0.1 U/ml thrombin in combination with: "none" (*N* = 21, *n* = 230), "ADP" (*N* = 3, *n* = 61), "serotonin" (*N* = 4, *n* = 66), "ATP" (*N* = 3, *n* = 37). Mann-Whitney U test: ** is p-value < 0.01, ns – not significant. (B) Time of the 1st release event. See legend to panel A for more details. Mann-Whitney U test: **** is p-value < 0.0001, ns – not significant. (C) Number of calcium spikes before the 1st release event. See legend to panel A for more details. Mann-Whitney U test: *** is p-value < 0.001, ns – not significant. (D) Population secretion curve of percent of the remaining granules over time for secreting platelets. Platelets were activated using 0.1 U/ml thrombin in combination with: "none" (*N* = 21, *n* = 230), "ADP & ATP & serotonin" (*N* = 5, *n* = 75), "ADP" (*N* = 3, *n* = 61), "serotonin" (*N* = 4, *n* = 66). Solid curve is the mean, while dashed curves are SEM. (E) Frequency of calcium spikes before the 1st release event. Platelets were activated using 0.1 U/ml thrombin in combination with: "none" (*N* = 21, *n* = 230), "SC" (*N* = 9, *n* = 98), "ADP" (*N* = 3, *n* = 61). Mann-Whitney U test: ** is p-value < 0.01, * is p-value < 0.05, ns – not significant.

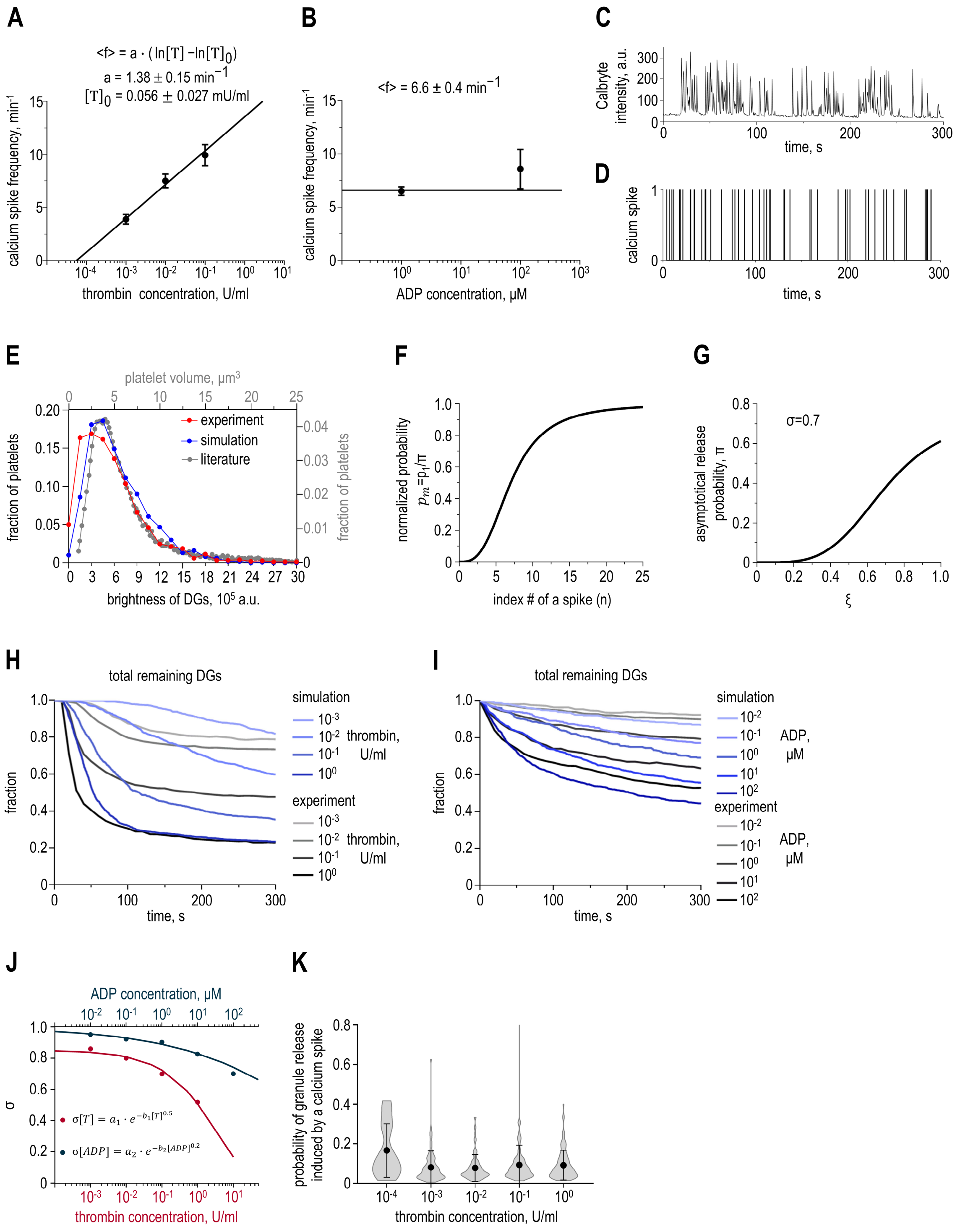


**Supplementary Figure 9. Modeling calcium oscillations and DG number and secretion in response to different activator concentrations.** (A) Dots are experimentally measured calcium spike frequencies (mean ± SEM) for platelets activated by thrombin plotted on a semi-log scale, also presented in Supplementary Figure 6D. Line shows theoretical dependency. (B) Dots are experimentally measured calcium spike frequencies (mean ± SEM) for platelets activated by ADP plotted on a semi-log scale. Line shows theoretical dependency. (C) Calbryte intensity curve of a single platelet activated by 0.1 U/ml thrombin at 0 s. (D) Simulated sequence of calcium spikes of a single platelet activated by 0.1 U/ml thrombin. In the graph, 0 indicates the absence of a calcium spike, and 1 indicates a calcium spike. (E) Distributions of platelet volume and DG brightness.

The red line corresponds to the experimental distribution of the total brightness of DGs, which is plotted in Supplementary Figure 4I as the number of minimally-sized DGs. The grey line corresponds to the distribution of platelet volumes plotted based on data in ref[2]. The blue line corresponds to the distribution of the number of DGs multiplied by 30,000 a.u. obtained in the simulations of 300 platelets. (F) Calcium-dependent increase in the normalized maturation rate embedded in the model. (G) Dependence of the asymptotic maturation rate on the random variable unique to a platelet. (H) Comparison of experimental and simulated secretion curves for indicated thrombin concentrations. Population secretion curves are plotted as fractions of remaining DGs over time. Experimental curves (shades of grey) are also presented in Supplementary Figure 5G. Simulated curves (shades of blue) are utilized for determining values of the σ parameter for corresponding activator concentrations. Each simulation secretion curve is an average of 100 platelet simulations. (I) Comparison of experimental and simulated population secretion curves for indicated ADP concentrations. See legend to panel H for more details. (J) Calibration of secretion response to different activator concentrations. The dots correspond to the values of the σ parameter employed to generate the curves in panels H and I. Fitting is employed to establish the continuous dependency of the σ parameter on activator concentration. The blue dots and curve correspond to thrombin activation, while the red dots and curve correspond to ADP activation. (K) Predicted apparent probability of a granule release in response to a calcium spike in platelets activated by indicated thrombin concentrations. In the graph, dots with whiskers represent mean ± SD and contours show data distribution based on 100 simulations of platelets for each thrombin concentration.

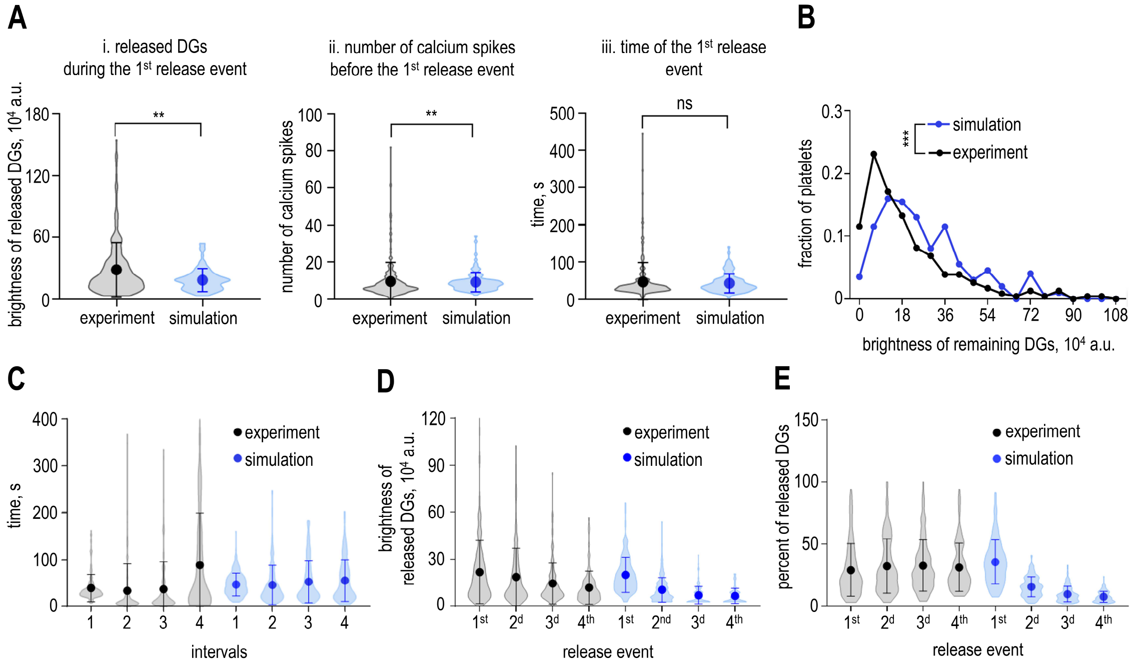


**Supplementary Figure 10. Validation of the bursting secretion model.** (A) Comparison of parameters of 1$^{st}$ release event obtained in experiment and in bursting secretion model. Dots represent the means; whiskers represent SD, and contours show the distribution of the data. The simulation was done for 100 platelets activated by 0.1 U/ml thrombin. Experimental data is also presented in Figure 5B-D. (i) is the brightness of released DGs during the 1$^{st}$ release event, (ii) is the number of spikes before the 1$^{st}$ release event, (iii) is the time before the 1$^{st}$ release event. Mann-Whitney U test: ** is p-value < 0.01, ns – not significant. (B) Distributions of remaining DGs obtained in the bursting secretion model (blue curve) and in the experiment (black curve) in platelets activated by 0.1 U/ml thrombin. DGs remaining in platelets were measured at 300 s of simulation or at 300 s after activator addition in the experiment. Experimental data are shown for $N$ = 20, $n$ = 230. The simulation was performed for 200 platelets. Kolmogorov-Smirnov test: *** is p-value < 0.001. (C) Time intervals between release events. Here and in panels D, E: dots are the means, whiskers are SD and contours show the data distribution. Experimental data are shown for $N$ = 14, $n$ = 171, 159, 129 and 89 platelets for consecutive events, see Data Source File for more details. Simulation of bursting secretion was done for 200 platelets. (D) Brightness of released DGs during release events. See legend to panel C for more details. (E) Percent of released DGs during release events. See legend to panel C for more details.

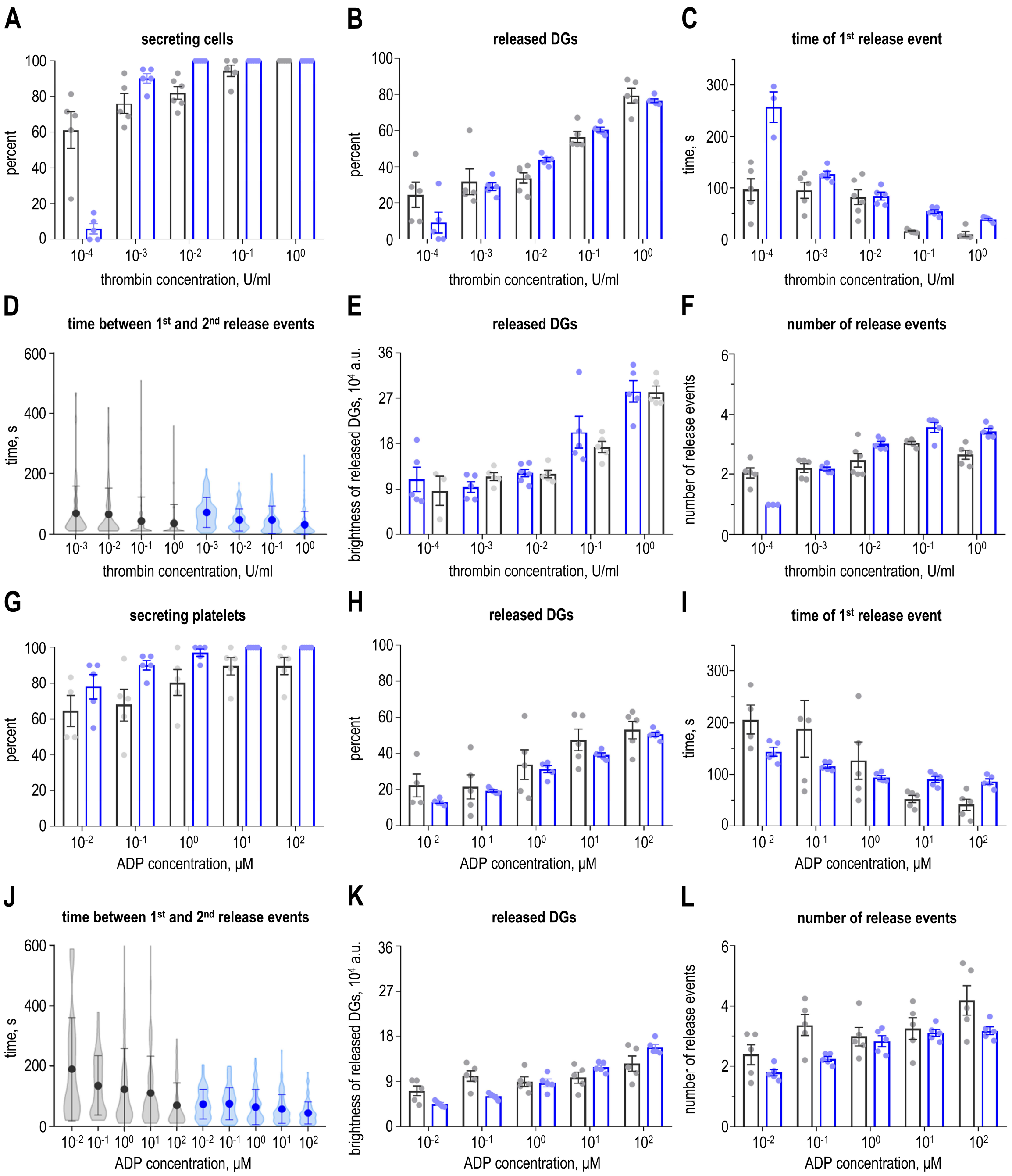


**Supplementary Figure 11. Model results of secretion in response to different activators.** Plots (A)-(F) show data for different thrombin concentrations, while plots (G)-(L) show data for different ADP concentrations. Concentrations are indicated on the x axis. The experimental data are shown in black, and the simulation results for bursting secretion model are shown in blue. The dots show the mean results of *N* individual experiments or the mean results of 5 groups of 20 simulated platelets in each group. The bars show the means ± SEM. The experimental data correspond to those presented in Figure 3 and Supplementary Figure 5, see legends for details. (A) and (G) Percent of cells that secreted DGs within 600 s after the addition of the activator in the experiment or following the start of the simulation. (B) and (H) Percent of released DGs within 600 s after the addition of the activator in the experiment or following the start of the simulation. (C) and (I) Time of the 1$^{st}$ release event after the addition of the activator in the experiment or following the start of the simulation. (D) and (J) Time between the 1$^{st}$ and 2$^{nd}$ release events. Dots represent the means; whiskers represent SD, and contours show the distribution of the data. (E) and (K) Brightness of released DGs during the 1st release event. (F) and (L) Number of release events within 600 s after the addition of the activator in the experiment or following the start of simulation.

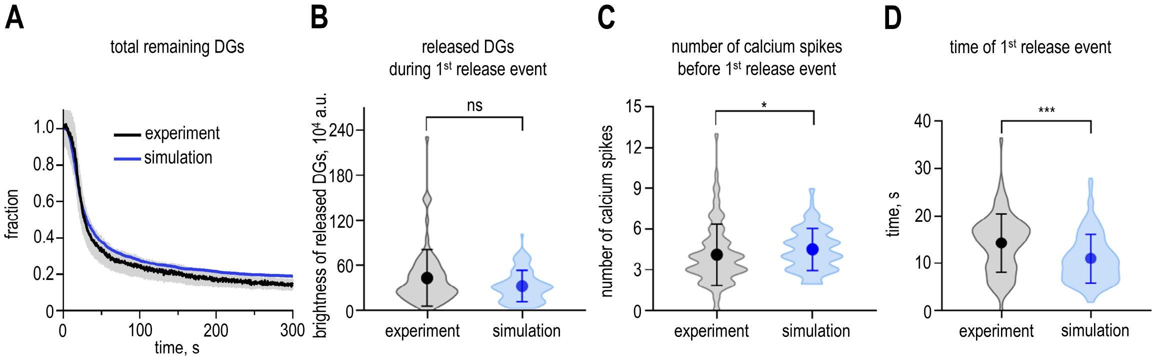


**Supplementary Figure 12. Model results of secretion in response to SC or components of DGs.** (A) Comparison of experimental and simulated population secretion curves for 0.1 U/ml thrombin activation with SC. The experimental curve (black) was plotted as the mean ± SEM of $N$ = 9, $n$ = 98. The simulated curve (blue) is an average of 300 platelet secretion curves. (B) Brightness of released DGs during the 1$^{st}$ release event. Dots represent the means; whiskers represent SEM and contours show the distribution of the data. The experimental data are also presented in Figure 5B. The simulated distribution is plotted for 100 platelets. Mann-Whitney U test: ns – not significant. (C) Number of calcium spikes before the 1$^{st}$ release event. The experimental data are also presented in Figure 5C. See legend to panel B for more details. Mann-Whitney U test: * is p-value < 0.05. (D) Time of the 1$^{st}$ release event. The experimental data are also presented in Figure 5D. See legend to panel B for more details. Mann-Whitney U test: *** is p-value < 0.001.

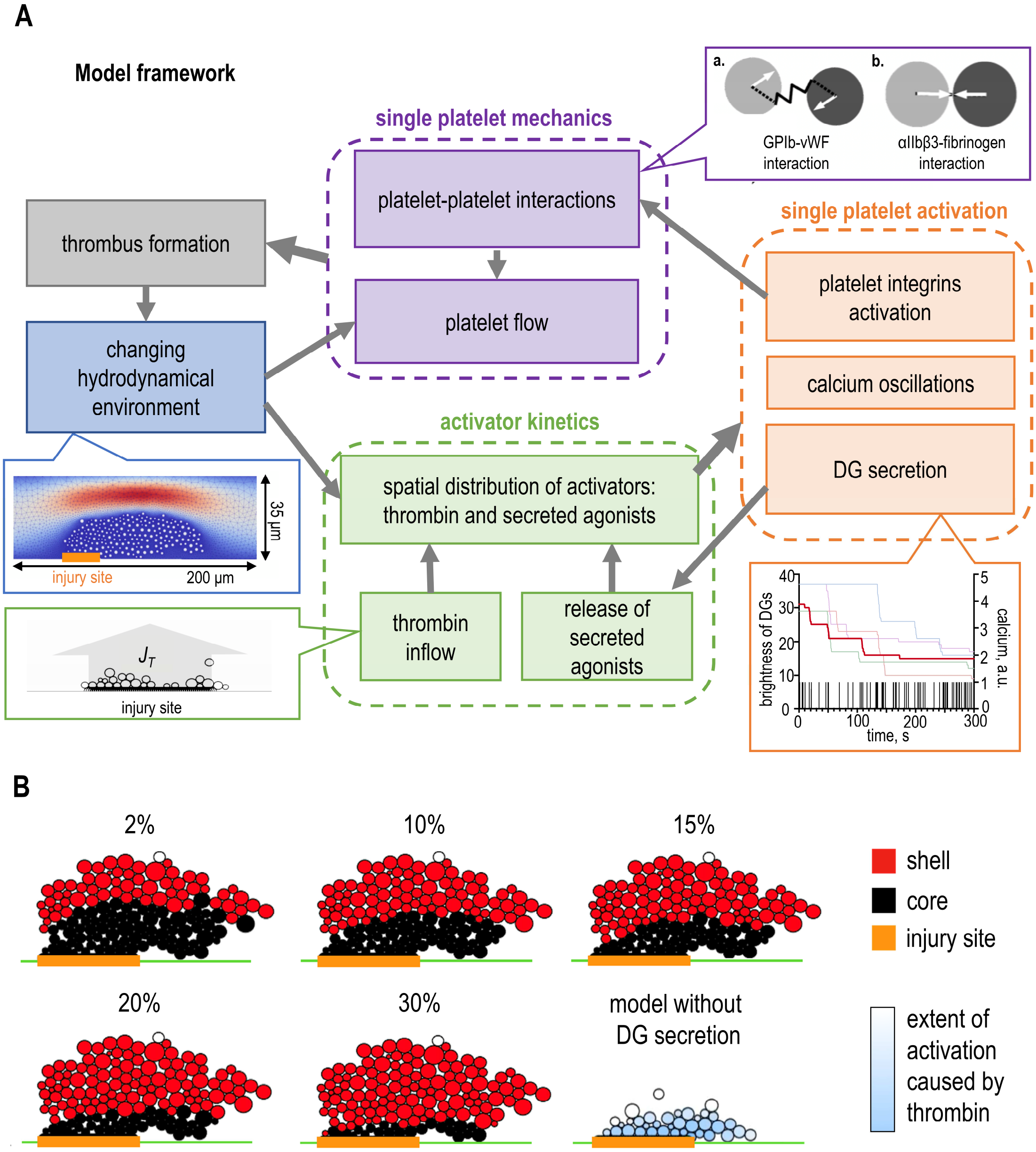


**Supplementary Figure 13. Modeling of the growing thrombus.** (A) Interdependence of different modules within the thrombus formation model. The processes contributing to thrombus formation in the model can be categorized into four groups: hydrodynamics of blood flow (blue box), mechanics of single platelets (purple box), kinetics of activators (green box), and activation of individual platelets (orange box). Thrombin enters the vessel through the injury site at a constant flux $J_T$ (shown in the green inbox). Thrombin activates cells by inducing mechanical interactions, calcium oscillations, and bursting secretion of DGs (an illustration of secretion kinetics for individual platelets is presented in the orange box), resulting in the release of secreted agonists that further activate platelets. Platelet interactions (demonstrated by two mechanisms in the purple box) and their transport via the flow lead to an aggregate formation. The growth of aggregate leads to the recalculation of the hydrodynamic conditions within the vessel (an example of a velocity field surrounding the thrombus is shown in the blue box). The updated blood flow transports platelets and activators. (B) Example distinguishing between the core and shell in the same simulated thrombus, determined by various values of the extent of platelet activation ($\alpha$). The extent of activation is provided as percent of the maximum possible extent of activation. The thrombus core is defined as an aggregate of platelets, each activated to an indicated percent by thrombin ($\alpha_T$), and is depicted in black. The thrombus shell is colored in red. The vessel wall is denoted by a green line, and the injury site is indicated by a bold orange line. The last image shows an example of a thrombus modeled without DG secretion. The blue color indicates the extent of activation solely caused by thrombin. A parameter $\alpha_T$ > 15% was selected as a threshold in the model to determine the core.

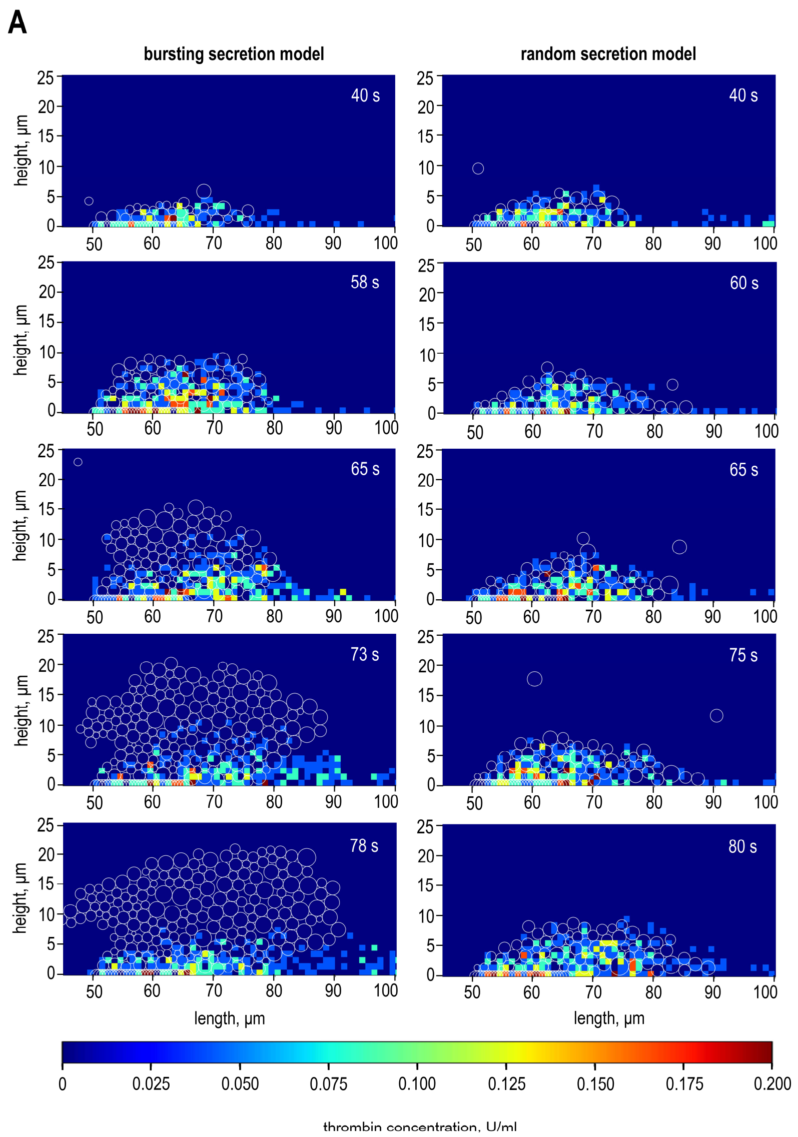


**Supplementary Figure 14. Thrombin distribution in growing thrombus.** (A) Thrombin distribution in the growing thrombus as simulated using the model with random (left) or bursting (right) secretion. The images show the identical region of the modeled vessel. The white empty disks are platelets. The platelets stacked at the bottom replicate the injury site, while flowing platelets adhere to the growing thrombus. The injury site is a source of thrombin, which diffuses away from the site and is diluted by flow. The color scale bar indicates thrombin concentration. Numbers correspond to the time in seconds after the start of the simulation.

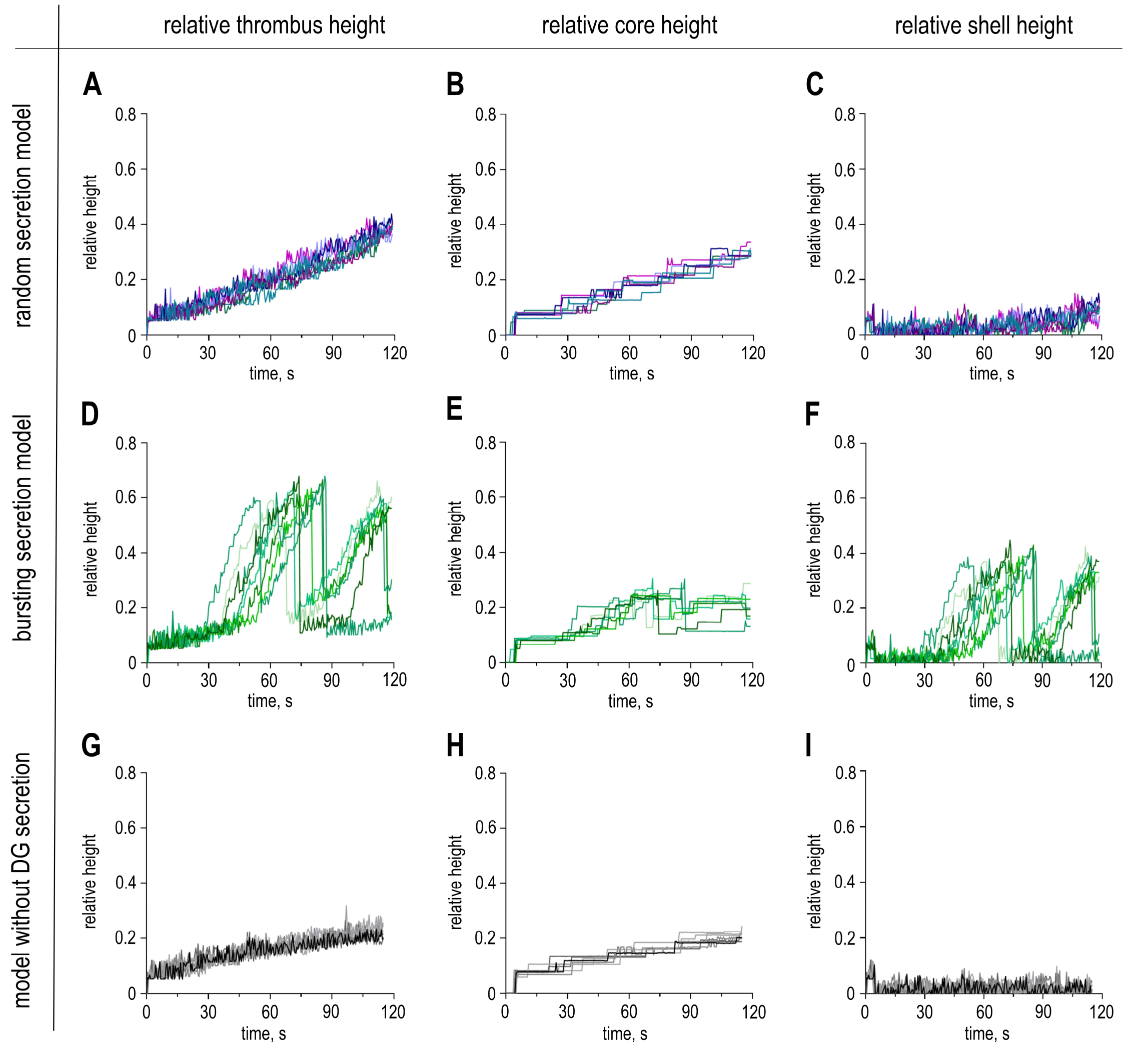


**Supplementary Figure 15. Thrombus growth kinetics in different simulation scenarios.** Rows correspond to different simulation scenarios from top to bottom: random secretion model, bursting secretion model and model without DG secretion. The thrombus core and shell are defined the same as in Supplementary Figure 13B. All heights are given relative to the vessel diameter. Each graph shows the curves of the 7 simulations for each scenario. (A), (D), (G) show the relative thrombus height. (B), (E), (H) show the relative thrombus core height. (C), (F), (I) show the relative thrombus shell height.

### 5. Supplementary Movie legends

**Supplementary Movie 1. Visualization of mepacrine-stained dense granules by SIM microscopy.**

This movie depicts a snapshot of a platelet adhered to a fibrinogen-coated coverslip in a flow chamber, with dense granules visualized using mepacrine staining. The initial segment presents a z-stack of super-resolution images, providing slice-by-slice optical sectioning of the stained granules. Numbers indicate the z-position (μm) relative to the coverslip. The second part of the movie features a 3D projection of the z-stack, offering a comprehensive view of granule localization within the platelet. The projection first rotates around the y-axis for a lateral perspective, followed by rotation around the x-axis for a top-down view, enhancing the visualization of spatial granule distribution. Numbers indicate the rotation angles. The movie plays at 1 fps. Scale bar: 0.5 μm.

**Supplementary Movie 2. Time-lapse visualization of mepacrine-stained dense granules in the absence of activators.**

This movie captures the time-dependent behavior of dense granules in a platelet adhered to a fibrinogen-coated coverslip in a flow chamber, recorded over 6.5 minutes using epi-fluorescence microscopy. The first frame presents the platelet under DIC illumination, with its boundary outlined in white in subsequent frames. Time stamps (minutes) are shown, with 0 min corresponding to buffer addition in this control experiment. Throughout the recording, dense granules exhibit small motions within the cell, but their overall brightness remains largely unchanged. The movie plays at 40 fps. Scale bar: 1 μm.

**Supplementary Movie 3. Kinetics of dense granule release following thrombin activation.** This movie depicts the same experiment as in Supplementary Movie 2, but with platelet activation by 0.1 U/ml thrombin at 0 min. The disappearance of small fluorescent dots and the gradual dimming of larger dots indicate dense granule release. The fluorescent image sequence is replayed second time showing arrows marking granules released during the same secretion event. The arrow colors and corresponding numbers denote the sequence of release events: (1$^{st}$ – yellow, 2$^{nd}$ – red, 3$^{rd}$ – cyan, 4$^{th}$ – magenta, 5$^{th}$ - yellow). The total imaging duration is 6 minutes, and the movie plays at 40 fps. Scale bar: 1 μm.

**Supplementary Movie 4. Dense granule release and changes in intracellular calcium in thrombin-activated platelets.**

Image sequences show two adjacent platelets adhered to a fibrinogen-coated coverslip. The first images were acquired with DIC microscopy to reveal cell morphology. Subsequent images acquired with epifluorescence show the secretion of mepacrine-labeled DGs (green) and intracellular calcium changes (Calbryte-590$^{AM}$ dye, red). Time stamps (minutes) are shown, with 0 min corresponding to addition of 0.1 U/ml thrombin. The total imaging duration is 7.5 min, and the movie plays at 40 fps. Scale bar is 2 μm.

**Supplementary Movie 5. Modeling result: thrombus formation by platelets with random dense granule secretion.**

The movie depicts platelet aggregation and activation in 2D thrombus model. The injury site, where thrombin is generated, is indicated with a thick grey bar along the horizontal axis. The vertical axis represents vessel width (μm), while numbers along the horizontal axis denote the distance along the vessel wall. The vertical axis is perpendicular to the vessel flow and shows the vessel width in microns. Time (s) is shown from the start of simulation. Platelets (white circles) flow from left to right at 1,000 $s^{-1}$ and adhere at the injury site due to activation by thrombin. Adhered platelets forming the core are shown as grey circles, while platelets in the shell layer are shown as green circles. The simulation duration is 80 s, and the movie plays at 10 fps.

**Supplementary Movie 6. Modeling result: Secreted agonist levels in a thrombus formed by platelets with random dense granule secretion.**

This movie depicts the same *in silico* experiment as in Supplementary Movie 5 but visualizes the spatio-temporal distribution of the secreted agonists using a color-coded scale. All platelets are represented as white empty circles. The colored bar on the right indicates the secreted agonist concentration in μM. The secreted agonists level remains low due to rapid dilution by flow. The simulation duration is 80 s, and the movie plays at 10 fps.

**Supplementary Movie 7. Modeling result: platelets with bursting dense granule secretion form a robust thrombus shell.**

This movie presents the results of an *in silico* experiment where bursting dense granule secretion is facilitated by positive feedback amplification from secreted agonists. Initially, thrombus growth resembles platelet aggregation without positive feedback. However, one minute into the simulation, the thrombus shell begins to expand rapidly. The shell comprises loosely adhered cells, as indicated by its deformation under flow. The total simulation duration is 80 seconds, with the movie playing at 10 frames per second.

**Supplementary Movie 8. Modeling result: Bursts of secreted agonists in a thrombus formed by platelets with bursting dense granule secretion.**

This movie depicts the same *in silico* experiment as in Supplementary Movie 7 but showing the spatiotemporal dynamics of secreted agonists. Explosive shell growth correlated with the appearance of a local burst with secreted agonists. The simulation duration is 80 s, and the movie plays at 10 fps.
